# Smooth muscle expression of RNA editing enzyme *ADAR1* controls activation of RNA sensor MDA5 in atherosclerosis

**DOI:** 10.1101/2024.07.08.602569

**Authors:** Chad S. Weldy, Qin Li, João P. Monteiro, Tim S. Peters, Hongchao Guo, Drew Galls, Wenduo Gu, Paul P. Cheng, Markus Ramste, Daniel Li, Brian T. Palmisano, Disha Sharma, Matthew D. Worssam, Quanyi Zhao, Amruta Bhate, Ramendra K. Kundu, Trieu Nguyen, Michal Mokry, Clint L. Miller, Sander W. van der Laan, Jin Billy Li, Thomas Quertermous

## Abstract

Mapping the genomic architecture of complex disease has been predicated on the understanding that genetic variants influence disease risk through modifying gene expression. However, recent discoveries have revealed that a significant burden of disease heritability in common autoinflammatory disorders and coronary artery disease (CAD) is mediated through genetic variation modifying post-transcriptional modification of RNA through adenosine-to-inosine (A-to-I) RNA editing. This common RNA modification is catalyzed by ADAR enzymes, where ADAR1 edits specific immunogenic double stranded RNA (dsRNA) to prevent activation of the double strand RNA (dsRNA) sensor MDA5 (*IFIH1*) and stimulation of an interferon stimulated gene (ISG) response. Multiple lines of human genetic data indicate impaired RNA editing and increased dsRNA sensing by MDA5 to be an important mechanism of CAD risk. Here, we provide a crucial link between observations in human genetics and mechanistic cell biology leading to progression of CAD. Through analysis of human atherosclerotic plaque and culture of human coronary artery vascular smooth muscle cells (SMCs) we implicate the SMC to have a distinct requirement for RNA editing, and that MDA5 activation regulates SMC phenotypic modulation. Through generation of a conditional SMC specific *Adar1* deletion mouse model on a pro-atherosclerosis background with additional constitutive deletion of MDA5 (*Ifih1*), and with incorporation of single cell RNA sequencing cellular profiling, we further show that Adar1 controls SMC phenotypic state by regulating Mda5 activation, is required to maintain vascular integrity, and controls progression of atherosclerosis and vascular calcification. Finally, we further corroborate our findings in a large human carotid endarterectomy dataset (Athero-Express) where we show that ISG activation is strongly associated with decreased plaque stability, increased SMC phenotypic modulation, and increased plaque calcification. Through this work, we describe a fundamental mechanism of CAD, where cell type and context specific RNA editing and sensing of dsRNA mediates disease progression, bridging our understanding of human genetics and disease causality.

**One Sentence Summary:** Smooth muscle expression of RNA editing enzyme ADAR1 regulates activation of double strand RNA sensor MDA5 in novel mechanism of atherosclerosis.

## INTRODUCTION

As coronary artery disease (CAD) is the worldwide leading cause of death^1^, there is enormous interest to map the genomic mechanisms of disease, establish biological causality, and accelerate therapeutic drug development. Despite the discovery of hundreds of genomic loci associated with CAD^2–5^ the promise of genome wide association studies (GWAS) to identify new mechanisms for treatment outside of lipid lowering therapy has not been fulfilled. Quantitative trait loci (QTL) studies mapping gene expression QTLs (eQTLs) have helped bridge GWAS variants to their molecular mechanisms^6,7^. However, post-transcriptional processes, such as RNA editing, have critical functions in health and disease and genetic determinants of RNA modification are important and poorly appreciated disease mechanisms^8–10^. Catalyzed by adenosine deaminase acting on RNA enzymes (ADARs), adenosine-to-inosine (A-to-I) RNA editing is a highly frequent RNA modification that is evolutionarily conserved across all metazoan species^8,11^. We have recently reported that common genetic variants that modify the frequency of A-to-I RNA editing, loci termed ‘editing-QTLs’ (edQTL), significantly increase CAD risk as well as numerous other autoinflammatory disorders^12^. This discovery raises the important possibility that genetically determined RNA editing function is a causal mechanism of CAD, yet the biological underpinning of this relationship is unclear.

Foundational work has revealed that ADAR1 (gene symbol *ADAR*) mediated RNA editing is a novel mechanism of inflammation and disease^13–16^. A principal function of ADAR1 is to bind double-stranded RNA (dsRNA) in non-coding repetitive elements and convert adenosines to inosines to facilitate the clearance of dsRNA molecules^17^. This contrasts with ADAR2 (gene symbol *ADARB1*) that serves to edit coding region of transcripts to modify transcript function^11,18^. ADAR1 is ubiquitously expressed across human tissues and its function to edit dsRNA suppresses the sensing of “non-self” dsRNA that is mediated by the cytosolic sensor of dsRNA, MDA5 (gene symbol *IFIH1*) (Fig 1A)^14,19,20^. Mice deficient in Adar1 editing are embryonic lethal due to activation of an interferon-stimulated gene (ISG) response, which is rescued to full life span when Mda5 (*Ifih1*) is also knocked out^14^. In humans, rare variants resulting in loss of function in ADAR1 or gain of function in MDA5 result in two severe and clinically overlapping Mendelian interferon mediated disorders, Acardi Goutières Syndrome and Singleton Merton Syndrome, both of which are characterized by a profound early onset vascular calcification phenotype demonstrating a link between RNA editing and dsRNA sensing in vascular disease^21,22^.

**Figure 1.**
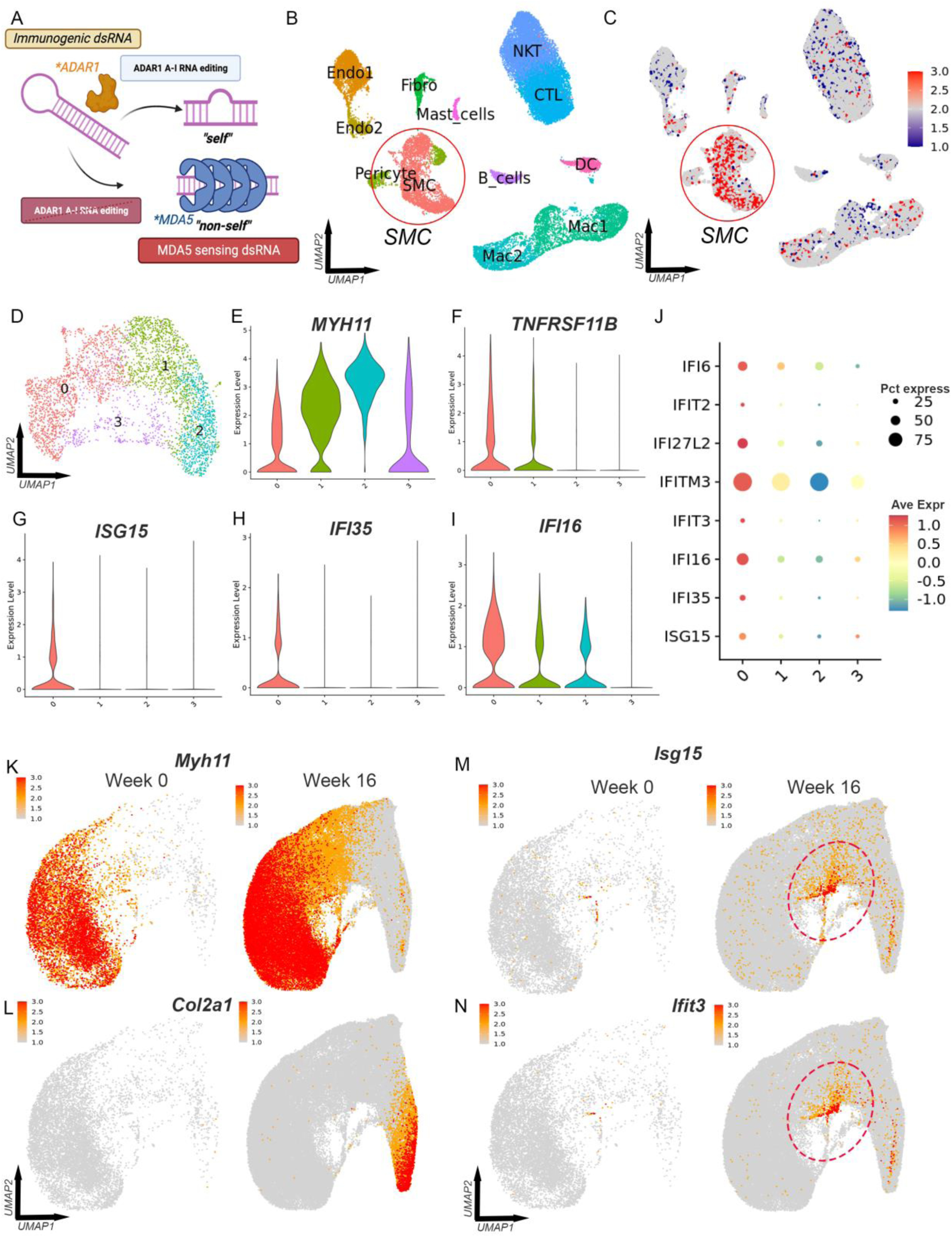
Smooth muscle cells in atherosclerosis express immunogenic RNA and show evidence of ISG induction in phenotypic modulation. (A) Schematic of ADAR1 RNA editing of dsRNA. (B) UMAP of scRNAseq data from human carotid atherosclerotic plaque (Alsaigh et al., 2022). (C) Module score of immunogenic RNA expression. (D) Clustering of SMC subset with (E) violin plot showing SMC marker *MYH11* with (F) fibromyocyte/chondromyocyte marker *TNFRSF11B*. (G-I) Violin plots of ISGs *ISG15* (G), *IF135* (H), and *IFI16* (I). (J) Dotplot of top ISGs within SMC clusters. (K-N) In mouse comprehensive integrated dataset of lineage traced SMCs (from Sharma et al., 2024), featureplot of *Myh11* (K) and *Col2a1* (L) at 0 and 16 weeks of high fat diet. (L-M) Featureplots of *Isg15* (L) and *Ifit3* (M) in SMC phenotypic modulation.

Common genetic variants that decrease RNA editing and increase CAD risk have similar effects on risk for autoinflammatory disorders such as inflammatory bowel disease and lupus^12^. To further link RNA editing to ISG response in human coronary arteries, risk allele-specific RNA editing was found to be lower and closely predicted an increased interferon gene response (implicating MDA5 activation) in the coronary arteries of patients with CAD^12^. Although prior work has identified ADAR1 as mediating atherosclerotic disease risk through RNA editing of specific RNA transcripts such as *NEAT1* and *CTSS*^23,24^, human genetic data corroborates a separate ‘axis’ of global ADAR1 RNA editing of immunogenic dsRNA to regulate MDA5 activation (ADAR1-dsRNA-MDA5 axis) as a mechanism of atherogenesis (Fig 1A). The relationship of this axis to CAD is further supported by the notable discovery that loss of function variants in *IFIH1* (MDA5) are protective against CAD^25^ and that the *IFIH1* locus meets genome wide level significant for association to CAD in GWAS^5^.

Here, by analyzing human atherosclerotic plaque single cell RNA sequencing (scRNAseq) data, we identified that vascular smooth muscle cells in atherosclerosis express RNA that require ADAR1 mediated RNA editing to evade MDA5 activation at a higher level than other cell types. Through *in vitro* culture of primary human coronary artery smooth muscle cells (HCASMCs) in relevant models, generation of genetic mouse models with conditional deletion of *Adar1* in vascular smooth muscle cells (SMCs) in homozygous loss of function and haploinsufficiency in atherosclerosis, extensive single cell RNA sequencing and histological characterization, with concurrent deletion of *Ifih1* (Mda5), in addition to a large human dataset to evaluate ISG activation and plaque phenotype (Athero-Express), we provide data to support the causality behind the genetic determinants of impaired RNA editing and CAD through MDA5 activation. We identify RNA editing in SMC as a mediator of atherosclerosis and by doing so create a new paradigm linking human genetics, epigenomics, and vascular biology to CAD.

## RESULTS

### Expression of immunogenic RNA within vascular SMCs in human atherosclerotic plaque

Activation of MDA5 is dependent on the concentration of specific RNA molecules that are prone to the formation dsRNA structures and form a pattern recognized by MDA5^26^. By combining our data from experimental *in vitro* models^27^ along with human genetic edQTL analysis^12^, we generated a putative list of 629 high probability human RNA molecules that are ‘immunogenic’ and require RNA editing to evade sensing by MDA5 (Table S1). Given the relationship between edQTLs and CAD, we evaluated human scRNAseq data from carotid atherosclerotic plaque^28^ to determine if there is a cell type specific expression of these immunogenic RNA. We determined cell clusters and cell types (Fig 1B) and generated a cell module ‘score’ that summates the expression of all 629 high probability ‘immunogenic’ RNA. Human SMCs expressed these immunogenic RNA at a higher level compared to all other cell types (Fig 1C). This observation may suggest a distinct role for SMCs in mediating RNA editing within the vessel wall.

### SMC phenotypic modulation and ISG induction in atherosclerosis

The formation of atherosclerotic plaque and vascular calcification involves SMCs undergoing a process of epigenetic reprogramming, termed “phenotypic modulation”^29,30^. In this process, SMCs dedifferentiate, down-regulate lineage marker expression, and migrate into the atheroma where these cells then proliferate and differentiate into different progenies that affect the biology of the plaque. Using SMC lineage tracing mouse models of atherosclerosis with single cell RNA and ATAC sequencing of the aortic root, we have previously shown that numerous CAD risk genes mediate disease risk through modification of this phenotypic modulation, including *TCF21*, *ZEB2*, *AHR*, *PDGFD* and *SMAD3*^31–35^. Mature SMCs will undergo phenotypic modulation to a more ‘fibroblast like’ fibromyocyte (FMC) and then into a calcification promoting chondromyocyte (CMC)^33^ phenotype in the setting of disease.

Given SMCs express immunogenic RNA at a higher level than other cell types, we hypothesized that insufficient RNA editing and dsRNA sensing could be identified as expression of key ISGs in the SMC population. We performed a subset analysis of the SMC population in this dataset^28^ where clustering revealed 4 major clusters (cluster ‘0’, ‘1’, ‘2’, and ‘3’)(Fig 1D). SMCs undergo a process of downregulation of the mature SMC marker *MYH11* (Fig 1E, cluster 2 vs 0 — -2.92 log2 fold change, adjusted *p* value 1.98E-290) with upregulation of FMC/CMC markers including *TNFRSF11B*, *HAPLN1* and *LUM* (ex. *TNFRSF11B* Fig 1F, cluster 2 vs 0 — 1.52 log2 fold change, adjusted *p* value 3.07E-26) as SMCs transition from mature SMC cluster ‘2’ to FMC/CMC cluster ‘0’ — a finding we have previously reported^33^. However, findmarker analysis of these clusters identified numerous ISGs as top markers of the FMC/CMC cluster ‘0’, including *ISG15*, *IFI35*, *IFI16*, *IFI27*, *IFIT3* (ex. *ISG15*, *IFI35*, *IFI16;* adj *p* value 3.94E-21, 2.60E-27, 2.63E-64, respectively; Fig 1G-J) (FindMarker results in Table S2). This finding may suggest dsRNA sensing by MDA5 with phenotypic modulation.

To investigate if ISG induction with SMC phenotypic modulation is conserved in mouse models of atherosclerosis, we evaluated our previously published comprehensive integrated dataset of multiple single cell transcriptomic experiments^36^. In this dataset, we determined the transcriptomic profiles in the aortic root of lineage traced SMCs in hyperlipidemic models of atherosclerosis^36^. We observed that in comparing between 0 and 16 weeks of high fat diet, SMCs underwent significant phenotypic modulation, marked by downregulation of SMC contractile marker *Myh11* (Fig 1K; -0.73 log2 fold change, *p* < 2E-16), with upregulation of *Col2a1* (Fig 1L; 3.29 log2 fold change, *p* < 2E-16), a marker of chondromyocyte formation. With this phenotypic transition, we observed significant upregulation of ISGs including *Isg15* (Fig 1M; 0.29 log2 fold change, *p* < 2E-16) and *Ifit3* (Fig 1N) in these phenotypically modified SMCs. These cells lie at the juncture between FMC and CMC formation and expression was absent in mature SMCs prior to phenotypic modulation.

### SMC ADAR1 and MDA5 expression in atherosclerosis in mice

To understand the expression patterns of *ADAR* and *IFIH1* (MDA5) in atherosclerosis SMC phenotypic modulation, we evaluated the expression of *ADAR* and *IFIH1* in this human carotid atherosclerosis dataset where we observed no differences in the expression of either *ADAR* or *IFIH1* across the different SMC subsets (Fig S1A-B)(*p* > 0.05). We then evaluated if expression of *ADAR* and *IFIH1* varied significantly across cell types in this human carotid atherosclerosis dataset, where we observed that expression of *ADAR* appeared largely consistent between SMCs, Endos, Fibros, and other immune cells, however *IFIH1* expression appeared highest in endothelial cells, followed by macrophage and SMCs (Fig S1C).

With evaluation of our comprehensive SMC lineage traced mouse dataset comparing week 0 and week 16, we observed a small increase in *Adar* and *Ifih1* at 16 weeks compared to the 0-week baseline (Fig S1D-F, *Adar* 1.26 fold *p*=2.2e-7; *Ifih1* 2.01 fold *p<2e-16*). At the 16-week timepoint, *Adar* expression appears largely consistent across SMC populations with phenotypic modulation, however *Ifih1* (Mda5) increases significantly at the FMC/CMC junction as observed with other ISGs (Fig S1D-F).

### ADAR1 controls global RNA editing and ISG response in human coronary artery smooth muscle cells

To investigate ADAR1 in human SMCs, we used an *in vitro* model of phenotypic modulation with primary human coronary artery smooth muscle cells (HCASMCs) and knocked down (KD) *ADAR* (ADAR1) and *IFIH1* (MDA5) with siRNA transfection paired with bulk RNA sequencing (RNAseq). In this model (Fig S2A), we cultured HCASMCs to confluency, treated with siRNA for 48 hours, and then transitioned to serum free medium (serum starvation) to induce a mature SMC phenotype marked by high expression of SMC markers (*e.g. ACTA2*, *CNN1*). Following 48 hours of serum starvation, cells were stimulated with either serum replete medium or maintained in a serum free medium for 48 hours. With serum stimulation, HCASMCs subsequently downregulated expression of mature SMC markers, a phenotype akin to phenotypic modulation in the SMC to FMC transition in atherosclerosis^33^.

With RNAseq data we inferred total RNA editing by detection of G base sequences at RNA editing sites as we have done previously (Fig S2B)^12,37^. We observed a notably strong linear relationship between *ADAR* expression and global RNA editing frequency (r^2^=0.9509, *p<0.0001*) (Fig S2B&C). Intriguingly, RNA editing frequency significantly decreased with serum stimulation alone (Fig S2C).

*ADAR* siRNA transfection in serum starved conditions induced a significant gene expression response with 6,351 differentially expressed genes (*FDR <0.05*) (Fig S2D). Top upregulated genes were interferon stimulated genes (ISGs) including *ISG15, IFI6*, *IFI44L*, *IFIT2*, and *IFIT1* (Fig S2D). With transfection of both *ADAR* (ADAR1) and *IFIH1* (MDA5) siRNAs we observed that co-KD of *IFIH1* blocked the majority of the gene expression response (Fig S2E).

With serum stimulation (representing phenotypic modulation) we observed *ADAR* KD induced a much greater effect than seen with serum starvation with 12,747 differentially expressed genes (*FDR <0.05*) (Fig 2F). This effect was characterized with a dominant ISG response and appeared nearly entirely dependent on activation of MDA5 (Fig 2G). When using principal component analysis (PCA) of bulk RNAseq data for each treatment group, we observed that the two dominant principal components tracked with MDA5 activation (PC1) and serum stimulation (PC2), where serum starved cells with *ADAR* KD produced an outwards shift on PC1, and then returned to near baseline with co-KD of *IFIH1* (Fig 2H). With serum stimulation, there was an upwards shift on PC2, and an even greater shift on PC1 with *ADAR* KD, and a return to baseline with co-KD of *IFIH1* (Fig 2H). These findings highlight that the dominant effect of impaired RNA editing in HCASMCs is driven by activation of MDA5 and SMC cell state.

**Figure 2.**
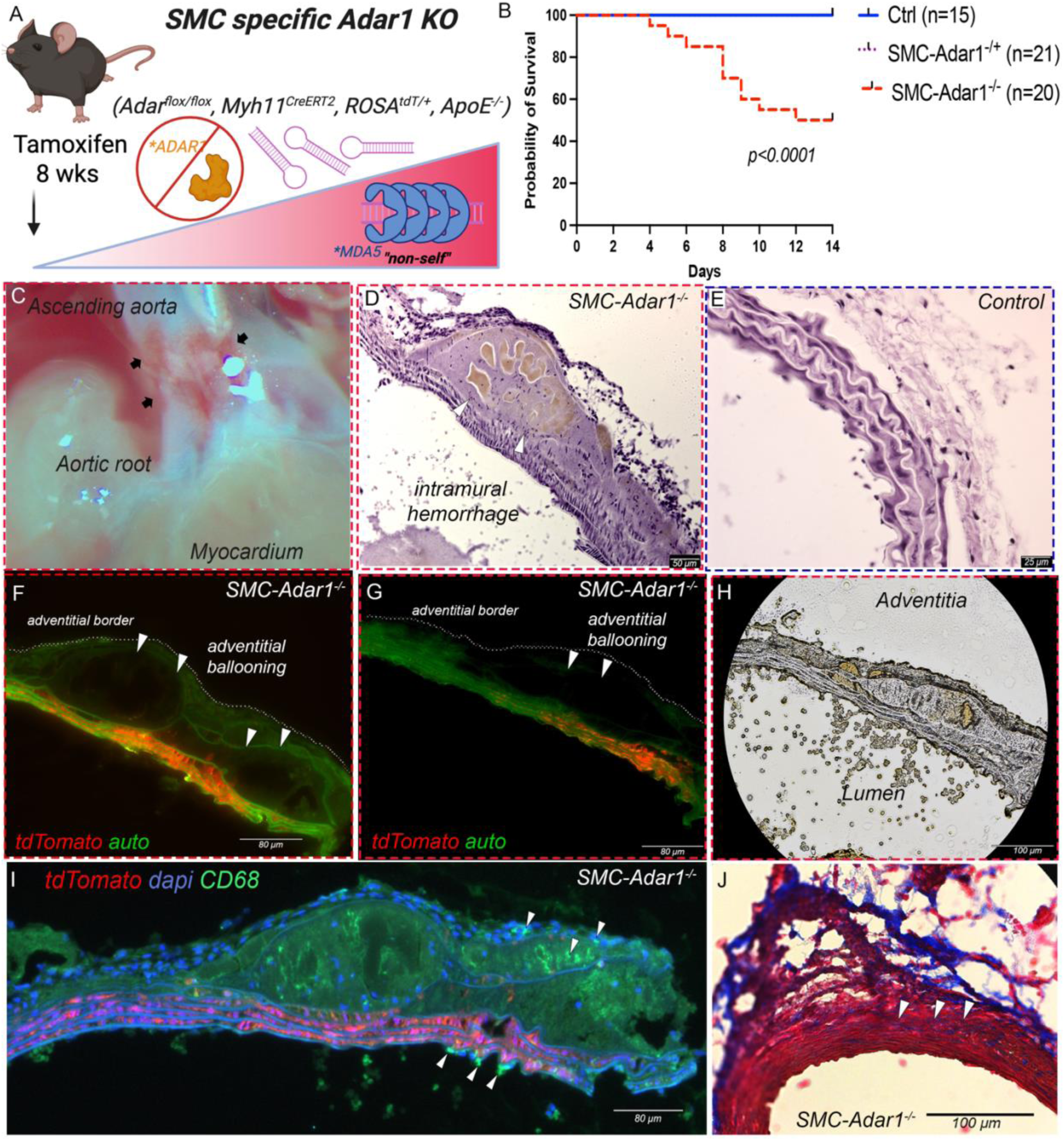
SMC specific ADAR1 is required to maintain vascular integrity and survival. (A) Schematic of SMC-*Adar1^-/-^* model and accumulation of dsRNA and MDA5 activation. (B) Survival curves for 56 mice treated with tamoxifen at 8 weeks of age, comparing *SMC-Adar1^-/-^*, *SMC-Adar1^-/+^* (*Adar1*^fl/WT^, *Myh11^CreERT2^*, *ROSAtdTomato*, *ApoE^-/-^*), and control (*Adar1*^WT/WT^, *Myh11^CreERT2^*, *ROSAtdTomato*, *ApoE^-/-^*) genotypes. (C) Gross image of aortic root and ascending aorta of *SMC-Adar1^KO^* mouse at 2 weeks following tamoxifen treatment. (D-E) Representative images of hematoxylin and eosin (H&E) staining from *SMC-Adar1^-/-^* (D) and control genotype mice (E). (F-G) Fluorescent imaging with autofluorescence indicating elastin (green) and Myh11 derived cells labeled with tdTomato (red) in two mice with adventitial ballooning in *SMC-Adar1^-/-^* (F-G). (H) Bright field imaging of aorta cross section from *SMC-Adar1^-/-^* mouse corresponding to fluorescent imaging in panel G. (I) Cd68 immunostaining of aorta cross section from *SMC-Adar1^-/-^* mouse showing macrophage infiltration. (J) Masson trichrome stain of aorta cross section from *SMC-Adar1^-/-^* mouse showing intra-medial collagen deposition. *P-*value represents comparison of survival curves, Log-rank (Mantel-Cox) test (B).

### ADAR1 and MDA5 regulate SMC maturation and key CAD transcription factors in phenotypic transition

In this HCASMC model of phenotypic transition, we observed that serum stimulation downregulated mature SMC markers including *ACTA2* and *CNN1* (Fig S2I&J). However, we also observed that *ADAR* KD alone downregulated these SMC markers, an effect that was accentuated with serum stimulation (Fig S2I&J). Using *ISG15* expression as a marker of ISG activation, *ADAR* KD induced an increase in *ISG15* expression (Fig S2K), an effect that was enhanced with serum stimulation and entirely blocked with co-KD of *IFIH1* (MDA5) (Fig S2K). Interestingly, numerous other key transcription factors with known functions in SMC maturation and risk of CAD (*i.e. KLF4*, *EGR1*, and *ATF3*)^38^ were similarly affected with *ADAR* KD (Fig S2L-2M).

Notably, classical interferon genes interferon α, β, and γ (i.e. *IFNA1*, *IFNB1, IFNG*) were either not expressed (*IFNA1* & *IFNG*), or minimally affected by *ADAR* KD in an MDA5 dependent mechanism (*IFNB1*) (Fig S3A-C) with similar minimal effect on the key IFN receptor, *IFNAR1* (Fig S3D). Top KEGG pathways of upregulated genes indicated a global viral sensing and inflammatory response, including “TNF signaling pathway”, “Influenza A”, “NOD-like receptor pathway”, and “EBV infection”, as well as “Lipid and atherosclerosis” (Fig S3E). GO biological processes similarly revealed “Defense Response to Virus” as the top process (Fig S3F). Downregulated pathways highlighted “Ribosome”, “Focal Adhesion”, and “ECM-receptor interaction” (Fig S3G) and GO terms suggested downregulation of RNA and protein processing with “Cytoplasmic translation” and related terms (Fig S3H).

### Global RNA editing is decreased in HCASMC following phenotypic modulation in vitro

In the evaluation of global RNA editing frequency in HCASMC with phenotypic modulation (serum stimulation) with control, *ADAR,* or *ADAR* + *IFIH1* KD, we observed that serum stimulation alone appeared to decrease total RNA editing (Fig S2C). Given we see serum stimulation to increase MDA5 activation with *ADAR* KD, this suggests a potential interaction between phenotypic modulation, decreased RNA editing, and dsRNA formation. To better understand this relationship, we performed a genome wide RNA editing site specific comparison in serum starved and serum stimulated HCASMC (Fig S4). In comparing the mean RNA editing level for each RNA editing site in control siRNA treated cells in serum starved and serum stimulated conditions, we observed a significant shift where serum stimulated cells have an overall decrease in RNA editing (Fig S4A). Although no specific RNA editing site met genome wide level significance, there was a significant overall net effect to decrease RNA editing frequency across hundreds of RNA editing sites. To evaluate the effect of *ADAR* KD we performed this editing site specific analysis in the *ADAR* KD with concurrent *IFIH1* KD (minimize the effect of MDA5 activation on ADAR1 editing) that revealed a massive reduction in RNA editing with *ADAR* KD in both serum starved, and serum stimulated conditions (Fig S4B&C). From this analysis, we then identified 1,101 differentially edited sites (DES) across 219 genes (Fig S4D&E) that are ADAR1 dependent. Most of these DES are shared between serum starved and serum stimulated conditions (Fig S4D&E), however we identified 226 sites within 38 genes that are distinct to serum stimulated conditions, and 357 sites within 49 genes that are distinct to serum starved conditions. Evaluation of these genes show no consistent biological pattern where gene ontology enrichment analysis shows no significant pathways (data not shown), however several genes (*TCTA, SWAP70, FN1, GGCX*) have been identified to reside at loci that meet genome wide level significance for CAD^5^. These data implicate specific genes that may be responsible for this enhanced MDA5 activation seen with serum stimulation.

To understand how serum stimulation may influence the overall expression of immunogenic RNA, we analyzed the expression change of the 629 previously identified immunogenic RNA (Table S1) and performed differential gene expression analysis. We observed that in HCASMC, serum stimulation elicited a notable change in gene expression of these selected genes with 82% differentially expressed (Fig S4G), however there appeared to be a relative balance between upregulated (249) and downregulated (239) genes. We hypothesized that the overall expression of these combined immunogenic RNA may have significance, where we measured the overall expression of all 629 immunogenic RNA by normalized RNAseq read counts. We observed that serum stimulation did not change the overall expression of the total immunogenic RNAs by combined RNA read count, however we did observe that with *ADAR* KD and MDA5 (*IFIH1*) activation, the expression of these RNAs trended down in the serum starved cells (p=0.11), and then significantly decreased in the serum stimulated cells (p=0.0022) in an MDA5 dependent mechanism (Fig S4H). We then looked at two distinct immunogenic RNA that colocalize with CAD GWAS where serum stimulation results in upregulation, *SWAP70* and *TCTA*. We observe that serum stimulation leads to upregulation, however *ADAR* KD blocks this effect in an MDA5 (*IFIH1*) dependent mechanism (Fig S4I&J). These data may suggest that MDA5 activation can regulate the global expression of immunogenic RNA.

### TGFβ attenuates MDA5 and ISG activation in HCASMC

As we observed *ADAR* and *IFIH1* (MDA5) to have distinct effects on HCASMC response to serum stimulation (a “synthetic” response), we aimed to further understand how *ADAR*, *IFIH1*, and *PKR* — a key ISG that acts to control cellular signaling following MDA5 activation^39^ — may further regulate HCASMC response to TGF*β* as a ‘contractile’ response. We cultured HCASMC to confluency, treated with siRNA for 48 hours, and transitioned cells to a serum free condition for 72 hours. We then stimulated cells with TGF*β* (10ng/mL) or control for 72 hours before we isolated RNA and performed quantitative PCR. When evaluating expression of key ISGs such as *ISG15*, we observed that TGF*β* treatment significantly blunted the effect of *ADAR* KD, where a 25-fold upregulation of *ISG15* with *ADAR* KD in serum starved conditions was limited to less than 10-fold upregulation with TGF*β* treatment (Fig S5A). Upregulation of *ISG15* was entirely blocked with KD of *IFIH1* (MDA5) and markedly reduced with *PKR* KD (Fig S5A). This finding suggests activation of *PKR* itself further activates the ISG response. In our prior serum stimulation study, *ADAR* KD induced a marked upregulation of *IFIH1* (MDA5) that was exacerbated with serum stimulation in a pattern that was consistent with all other ISGs (Fig S5B). However, TGF*β* treatment reduced this effect of *ADAR* KD (Fig S5C). In control cells, we observed that concurrent *PKR* KD further blunted the activation of *IFIH1* (MDA5)(Fig S5C). We then evaluated expression of *PKR* where we found *PKR* to similarly be upregulated by *ADAR* KD, however this effect was significantly reduced with TGF*β* treatment (Fig S5D) and in control cells was entirely blocked with *IFIH1* (MDA5) KD (Fig S5D).

To understand if the effect of TGF*β* to blunt the ISG response is dependent on canonical TGF*β* TFs such as SMAD3, we performed additional experiments with concurrent *SMAD3* KD and evaluated the expression if *ISG15* and *IFIH1*. Interestingly, we observed that *SMAD3* KD itself attenuated the upregulation of both *ISG15* and *IFIH1* (Fig S5E&F) with concurrent *ADAR* KD, suggesting that SMAD3 is playing a role to activate ISG response with *ADAR* KD. However, the effect of TGF*β* treatment appeared to be the same despite *SMAD3* KD (Fig S5E&F). This may suggest that SMAD3 and TGF*β* have distinct yet contrasting roles in regulating ISG induction, implicating alternative, non-canonical TGF*β* signaling pathways to mediate this effect.

In comparing the effect of serum stimulation or TGF*β* treatment on *ADAR* expression, we observed that serum stimulation induced a small increase in *ADAR* expression (∼1.2 fold, Fig S5G)(similar to what we observed *in vivo* during atherosclerosis, Fig S1), whereas TGF*β* treatment produced a somewhat greater 2-fold upregulation of *ADAR* (Fig S5H). At this timepoint, 8 days after siRNA treatment, *ADAR* expression in the *ADAR* KD treatment was found to have normalized in a manner dependent on *IFIH1* (MDA5), however not dependent on PKR, indicating that MDA5 activation is stimulating a compensatory upregulation of *ADAR* (Fig S5H).

To then understand how MDA5 and PKR may be regulating HCASMC response to TGF*β* and phenotypic modulation, we evaluated the expression of the mature SMC contractile marker *ACTA2* as well as TGF*β* responsive gene *PI16*. We observed that TGF*β* stimulation downregulates *ACTA2* in a manner similar to *ADAR* KD. In addition, downregulation of *ACTA2* with *ADAR* KD in control cells was dependent on MDA5 (*IFIH1*) and to a lesser extent *PKR* (Fig S5G). However, the effect of TGF*β* to downregulate *ACTA2* was entirely independent of *ADAR*, *IFIH1*, or *PKR* (Fig S5G). We found that TGF*β* treatment significantly upregulated *PI16*, however this effect was significantly blunted with KD of *ADAR*, in a manner dependent on both *IFIH1* (MDA5) and *PKR* (Fig S5H), suggesting activation of MDA5 and subsequently PKR may regulate SMC response to TGF*β*.

### ADAR1 and MDA5 regulate the transcriptomic and inflammatory response to calcification in vitro

To model SMC calcification *in vitro*, we cultured an immortalized HCASMC line under high calcium/phosphate culture conditions as has been previously reported^40^. In this model, SMCs begin to form deposits that stain with Alizarin red, consistent with calcium crystals and has been proposed to represent vascular calcification^40^. Following siRNA transfection for 48 hours, we cultured cells for 7 days with pro-calcification medium under low serum (1%) conditions and performed bulk RNAseq. After 7 days, calcification deposits became visible, however with *ADAR* KD, there was a significant phenotypic change with increased cell death and increased density of calcium deposits — an effect that was blocked with co-KD of *IFIH1* (Fig S6A&B). Calcification medium under scramble siRNA treatment promoted a significant gene expression response with 584 differentially expressed genes (*FDR <0.05*) (Fig S6C). Top upregulated genes include numerous BMP related signaling genes including *FBLN5*, *FMOD*, and *RANBP3L* (Fig S6D-F), effects that have been previously reported^40^. However, with *ADAR* KD this effect was blunted in an MDA5 dependent mechanism (Fig S6D-F).

Calcification did not overtly activate MDA5 as measured by expression of ISGs (i.e. *ISG15*)(Fig S6G&H), however, *ADAR* KD modified the HCASMC inflammatory response to calcification. Calcification had minimal effect on expression of NFκB (*NFKB1*), however with *ADAR* KD *NFKB1* is modestly upregulated and is further upregulated with calcification in *ADAR* KD cells, an effect blocked with co-KD of *IFIH1* (Fig S6I). This observation revealed a unique relationship where calcification induced a distinct proinflammatory effect only with concurrent *ADAR* KD and MDA5 activation. Similar effects were observed for numerous proinflammatory signaling and matrix proteins including *JUND*, *MMP1*, and *MMP10* (Fig S6J-L).

### SMC specific Adar1 maintains vascular integrity and survival in vivo

Although prior reports have implicated SMC specific RNA editing in mediating phenotypic response^41,42^, the relationship between deficient SMC RNA editing, MDA5 activation, vascular inflammation, and atherosclerosis has not been investigated. To understand how homozygous deletion of *Adar1* within SMCs influence vascular homeostasis, we bred a mouse model with four distinct alleles, including *Adar1*^fl/fl^ (exon 7-9), a tamoxifen inducible SMC specific Cre-recombinase (*Myh11^CreERT^*^2^), a lineage tracing fluorophore (*ROSA*^tdTomato^), on a hyperlipidemic pro-atherosclerosis genetic background (*ApoE^-/-^*) (Fig 2A). A total of 56 mice were treated, comparing *SMC-Adar1^-/-^*, *SMC-Adar1^-/+^* (*Adar1*^fl/WT^, *Myh11^CreERT^*^2^, *ROSA^tdTomato^*, *ApoE^-/-^*), and control (*Adar1*^WT/WT^, *Myh11^CreERT^*^2^, *ROSA^tdTomato^*, *ApoE^-/-^*) genotypes. We observed that at day 4, *SMC-Adar1^-/-^* mice became notably ill as characterized by weight loss, piloerection, and a hunched posture, a phenotype that was not present in *SMC-Adar1^-/+^* or control genotype animals. We observed mortality in the *SMC-Adar1^-/-^* mice at this time and by 14 days, 10/20 (50%) mice had died (Fig 2B). A recent report while this manuscript was in review has further supported this role of SMC specific *Adar1* to maintain survival and vascular integrity^43^. Gross dissection and visual inspection of *SMC-Adar1^-/-^* mice did not reveal prominent findings of the lungs, heart, or intestines however visual appearance of the aorta suggested petechial hemorrhage and erythema (Fig 2C). H&E staining (Fig 2D&E), fluorescent imaging (Fig 2F&G), bright field imaging (Fig 2H), Cd68 immunostaining (Fig 2I), and Masson’s trichrome staining (Fig 2J) of ascending and descending aortic tissue in 6 *SMC-Adar1^-/-^* mice revealed a loss of elastin organization (present in 6/6 mice), dissociation of the matrix at the adventitial aspect with associated ‘ballooning’ out and separation of layers (present in 3/6 mice), with evidence of intramural hemorrhage (present in 2/6 mice) (Fig 2D-H, Fig S7). These sites of ballooning occurred with complete loss of tdTomato SMCs at the adventitial aspect (Fig 2F&G). H&E staining of the aorta confirmed severe loss of organization of the elastin matrix and evidence of intravascular hemorrhage with further discovery of dense cellular infiltrate into the adventitia (Fig 2D&E). These patterns were not observed in control mice (Fig 2E). Immunostaining for CD68 revealed macrophage cell infiltration from the luminal as well as the adventitial aspect of the vessel wall (Fig 2I). Additional Masson’s trichrome stain to evaluate for fibrosis revealed evidence of intra-medial collagen deposition (Fig 2J).

In this model, there was a rapid and early inflammatory response that resulted in profound weight loss and mortality. Although the cause of death was not immediately apparent, as gross examination and H&E sections of liver, lung, and heart were nonrevealing, the acute systemic inflammatory response is likely eliciting death through a multifactorial mechanism akin to a systemic inflammatory response. There was no evidence of aortic rupture as a cause of death. However, mice that survived this period remained underweight (15-18 grams) and ill in appearance until ∼7 months of age. Importantly, no mice that were taken past this 2-week timepoint regained weight or lived a normal lifespan. Dissection of mice at this 7-month timepoint demonstrated dense fecal impaction of the large intestine, consistent with bowel obstruction, a finding not seen at earlier timepoints.

### Haploinsufficiency of MDA5 is sufficient to protect against mortality with SMC Adar1^-/-^ mice

Our *in vitro* work indicates MDA5 to be the principal mediator of downstream signaling with *ADAR1* KD in SMC, however it remains unclear if this relationship is true *in vivo*. In addition, it is unclear if only partial inhibition of MDA5 can prevent overt ISG activation, a model more akin to what might be achieved with future pharmacologic inhibition. To test this, we bred the SMC *Adar1* KO mouse onto a constitutive KO of *Ifih1* (Mda5) on both a heterozygous and homozygous background generating mice that are *SMC-Adar1^-/-^ + Mda5^-/+^* (*Adar1*^fl/fl^, *Ifih1^-/+^, Myh11^CreERT2^*, *ROSAtdTomato*, *ApoE^-/-^*) and *SMC-Adar1^-/-^ + Mda5^-/-^* (*Adar1*^fl/fl^, *Ifih1^-/-^, Myh11^CreERT2^*, *ROSAtdTomato*, *ApoE^-/-^*). Following the same model as shown previously (Fig 2), we performed an additional experiment using litter-mate controlled mice with tamoxifen treatment for a total of 32 mice (12 SMC-*Adar1^-/-^*, 8 *SMC-Adar1^-/-^ + Mda5^-/+^*, and 12 *SMC-Adar1^-/-^ + Mda5^-/-^*). Mice that were homozygous SMC *Adar1* KO with MDA5 WT had severe weight loss and significant mortality, with 10/12 (83%) *SMC-Adar1^-/-^* mice dying prior to the 14-day timepoint and greater than 30% loss of total body weight by 7 days for the 6 mice that survived to the 7-day timepoint (Fig 3A-C). Both *SMC-Adar1^-/-^ + Mda5^-/+^* and *SMC-Adar1^-/-^ + Mda5^-/-^* mice showed minimal weight loss, no mortality, and visually appeared largely unaffected by *Adar1* KO at this 2-week timepoint (Fig 3A-D). *SMC-Adar1^-/-^ + Mda5^-/+^* mice have been taken out to 24 weeks following tamoxifen dosing and continue to show minimal phenotype, demonstrating that MDA5 is required to initiate this severe phenotype with SMC *Adar1* KO. Histological evaluation of aortic tissue from *SMC-Adar1^-/-^, Mda5^-/+^* mice show that aortic cross section, aortic media, and elastin structures are indistinguishable from control mice (Fig S8). Intriguingly, a portion of both *SMC-Adar1^-/-^, Mda5^-/+^* and *SMC-Adar1^-/-^, Mda5^-/-^* mice (2/8 and 4/12, respectively) did lose significant weight within the first week, however had significant weight recovery by week 2 (Fig 3C). This suggests that although MDA5 deletion protects against mortality and any major phenotype, that alternative dsRNA receptors such as RIG-I may be mediating a partial inflammatory response.

**Figure 3.**
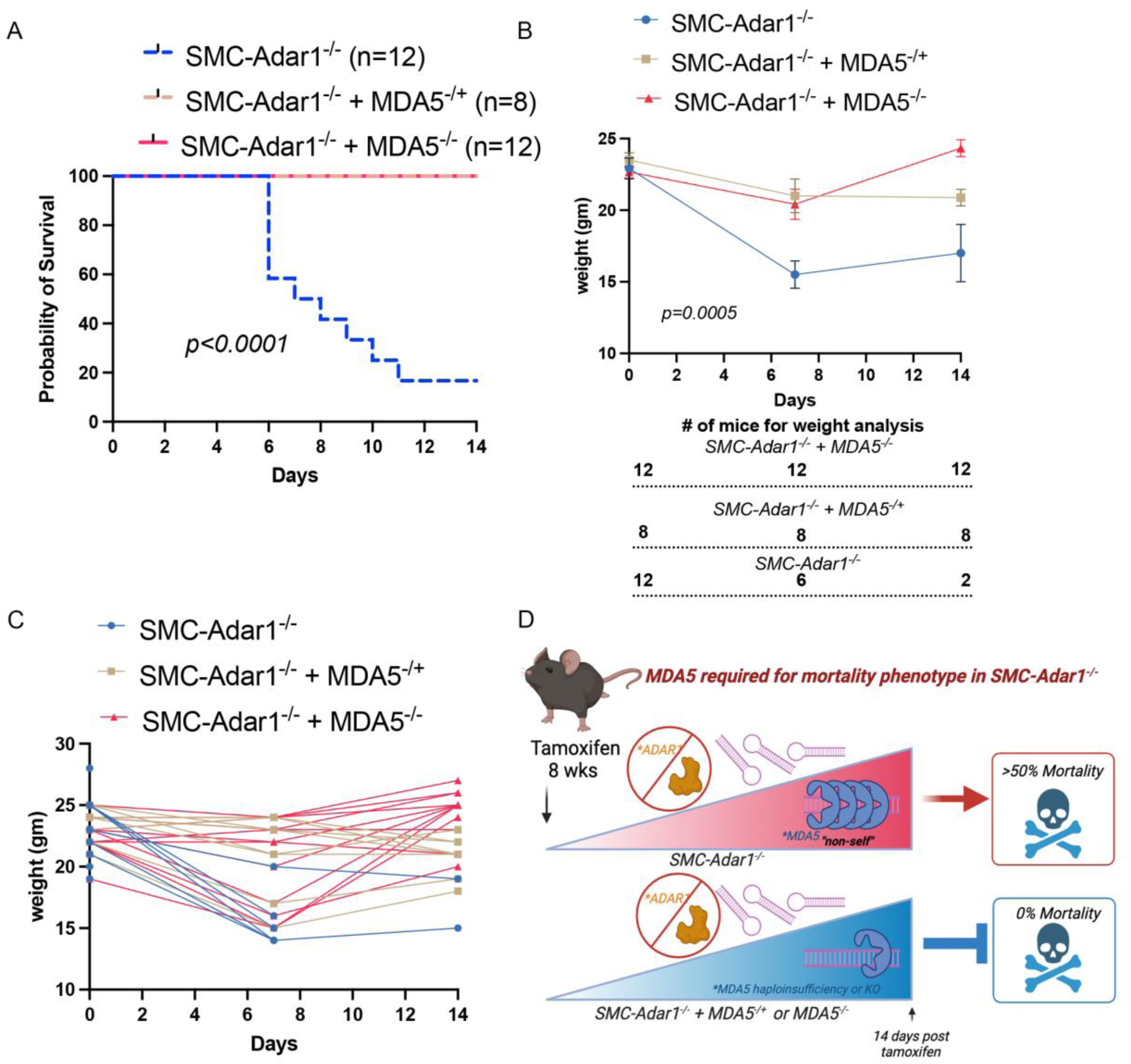
Haploinsufficiency in MDA5 is adequate to prevent mortality in SMC-Adar1^-/-^ mice. (A) Survival curves for *SMC-Adar1^-/-^* (*Adar1*^fl/fl^, *Myh11^CreERT2^*, *ROSAtdTomato*, *ApoE^-/-^*), *SMC-Adar1^-/-^*, *MDA5^-/+^* (*Adar1*^fl/fl^, *Ifih1^-/+^*, *Myh11^CreERT2^*, *ROSAtdTomato*, *ApoE^-/-^*), and *SMC-Adar1^-/-^*, *MDA5^-/-^*(*Adar1*^fl/fl^, *Ifih1^-/-^*, *Myh11^CreERT2^*, *ROSAtdTomato*, *ApoE^-/-^*) genotypes following tamoxifen treatment at 8 weeks of age. (B) Weight curve at 0, 7, and 14 days following tamoxifen treatment between genotypes of surviving mice. (C) Individual weight measures at 0, 7, and 14 days following tamoxifen treatment between genotypes of surviving mice. (D) Schematic diagram of *SMC-Adar1^-/-^* and MDA5 haploinsufficiency or global KO protecting against mortality. *P-*values represent comparison of survival curves Log-rank (Mantel-Cox) test (A-B). (B) Data point represents means (±SEM).

### Single cell RNA sequencing reveals SMC Adar1 controls ISG activation, phenotypic response, and macrophage infiltration into the vessel wall

To further characterize the vascular response of SMC specific *Adar1* KO in mouse, we performed single cell RNA sequencing (scRNAseq) of the aortic root and ascending aorta of *SMC-Adar1^-/-^* mice at day 14 following tamoxifen dosing. With a total of 10 individual scRNAseq captures, we characterized over 50,000 cells with high sequencing depth (>50K reads per cell). Using dimensionality reduction with UMAP as we have done previously^44,45^, we observed *SMC-Adar1^-/-^* aortic cells to have a dramatic transcriptomic change in comparison to control genotype (Fig 4A). Cluster analysis identified 13 main clusters, including 3 SMC clusters — a control SMC cluster marked by mature contractile markers (i.e. *Myh11* and *Cnn1*), and two SMC clusters that represented cells with distinct transcriptomic signatures secondary to SMC *Adar1* KO that are marked by tdTomato (clusters SMC_cAdar1_KD1; SMC_cAdar1_KD2) (Fig 4B&C) (Findmarker gene analysis in Table S3). Genes expressed by cells in these two clusters were found to show gene ontology (GO) enrichment for terms related to ‘response to interferon beta’, ‘response to virus’, ‘defense response to virus’ as top GO terms (Fig S9). Using *Isg15* as a marker of MDA5 activation, clusters SMC_cAdar1_KD1 and SMC_cAdar1_KD2 were marked by notably high expression of *Isg15* (Fig 4D). To validate this finding and characterize the location of *Isg15* expression, we performed RNAscope on aortic tissue from *SMC-Adar1^-/-^* mice. We observed robust *Isg15* expression within the vessel wall that was highest at the luminal aspect of the media, with additional expression within the adventitia (Fig 4E).

**Figure 4.**
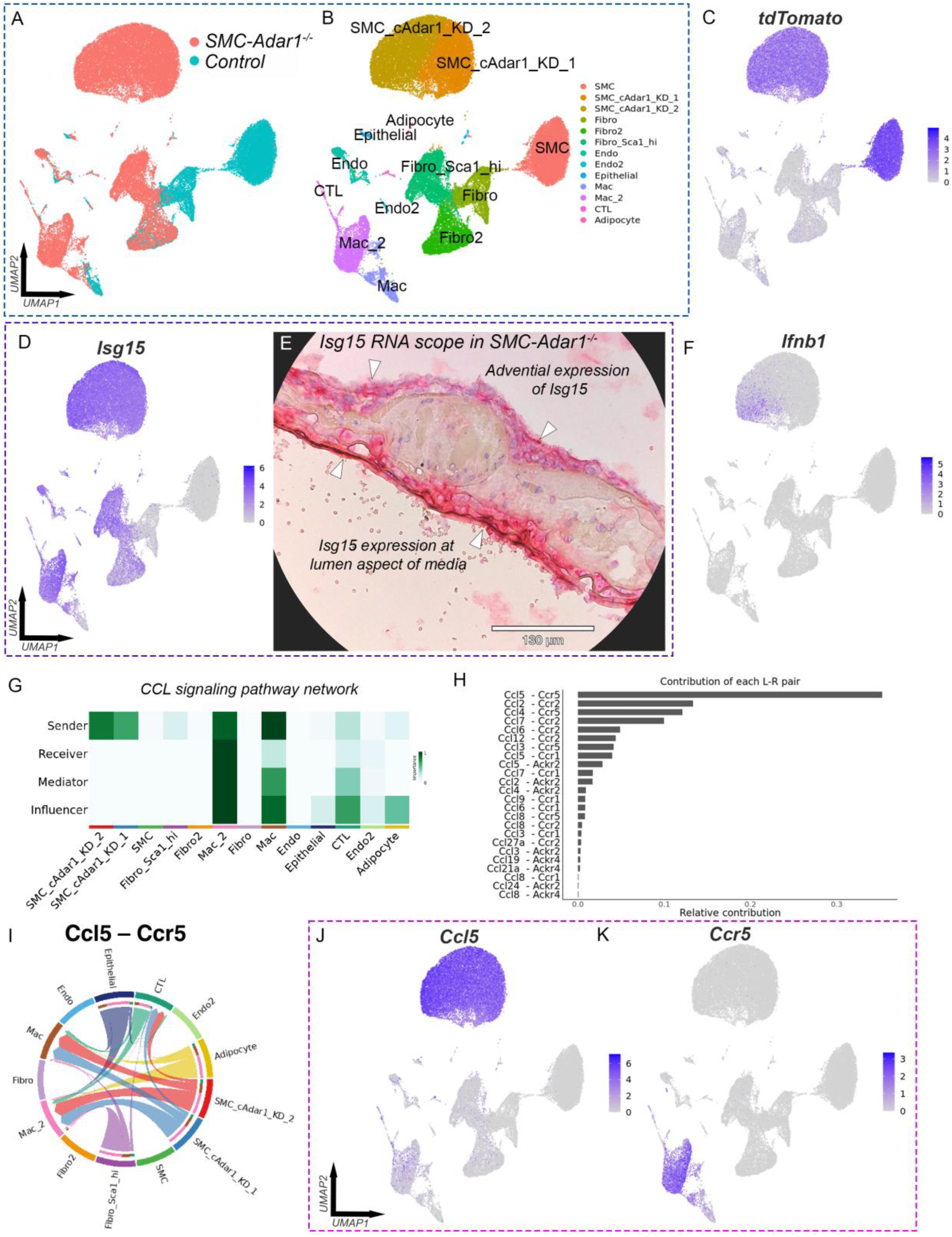
Single cell RNA sequencing reveals SMC Adar1 to control ISG activation, phenotypic response, and macrophage infiltration. (A) scRNAseq UMAP from aortic root and ascending aorta grouped by genotype of *SMC-Adar1^-/-^* (*Adar1*^fl/fl^, *Myh11^CreERT2^*, *ROSAtdTomato*, *ApoE^-/-^*) and control (*Adar1*^WT/WT^, *Myh11^CreERT2^*, *ROSAtdTomato*, *ApoE^-/-^*) genotypes at 2 weeks following tamoxifen treatment. (B) UMAP with clustering and cell type identification of combined *SMC-Adar1^-/-^* and control genotype data. (C) Featureplot of tdTomato showing SMC lineage tracing. (D) Featureplot of *Isg15*. (E) RNAscope of *Isg15* in aortic tissue of *SMC-Adar1^-/-^* mice showing highest *Isg15* expression at the lumen aspect of media, with additional adventitial expression. (F) Featureplot of *Ifnb1*. (G) Heatmap of network centrality scores for CCL signaling network across cell clusters. (H) Contributions of each receptor-ligand pair for CCL signaling. (I) Chord diagram of receptor-ligand interaction between clusters for CCL5:CCR5. (J-K) Featureplot of (J) *Ccl5* and (K) *Ccr5*.

Within tdTomato lineage traced SMCs, conditional *Adar1* KO produced marked gene expression changes with 2,382 differentially expressed genes (*FDR<0.05, log2FC>0.1*), with notable increase in ISGs. In comparing clusters SMC_cAdar1_KD1 and SMC_cAdar1_KD2, SMC_cAdar1_KD2 was marked by higher expression of numerous inflammatory signaling genes (i.e. *Isg15*, *Ccl5*, *Cxcl10*) (Fig S10A).

Importantly, consistent with our *in vitro* findings (Fig S3A-D), there was only modest activation of classical interferon genes with no observed expression of *Ifna1* and minor activation of *Ifnb1* (Fig 4F). Similarly, we observed minimal expression of IL6 family cytokines (i.e. *Il6*) with *SMC-Adar1*^-/-^ (Fig S10B). The minimal expression of these markers appeared limited to the more proinflammatory SMC_cAdar1_KD2 cluster. In addition, consistent with our *in vitro* findings, we observed that SMC *Adar1* KO led to downregulation of mature SMC markers (Fig S10C). By comparing the three major SMC clusters, expression of *Cnn1* and *Myh11* decreased in a pattern consistent with increasing MDA5 activation (Fig S10C).

There was a new distinct fibroblast population seen only with SMC *Adar1*^-/-^ and is marked by ISGs and high expression of *Ly6a* (Sca1) (Fig 4B, Fig S10D), consistent with a proinflammatory adventitial fibroblast population^46^. Similarly, in SMC *Adar1*^-/-^ there was a loss of cells within the ‘Fibro_1’ cluster that was only seen in controls (Fig 4B), suggesting a dynamic shift in gene expression from resident fibroblasts to this new reparative and inflammatory cell cluster. In addition, there was a large infiltrating macrophage population marked by high *Cd68* (resident macrophage cluster – ‘Mac’, infiltrating macrophage cluster – ‘Mac_2’) (Fig 4B, Fig S10E) with an additional small T cell population with SMC *Adar1*^-/-^ marked by high *Gzma*, *Cd3g*, and *Cd52*, suggestive of a cytotoxic T lymphocyte (CTL) population (Fig 4B, Fig S10F). These data indicate that with loss of *Adar1* within SMCs, a profound vascular response occurs throughout the vessel wall that is coordinated by SMCs.

### Receptor-ligand interaction modeling implicates CCL5:CCR5 to coordinate macrophage infiltration in SMC-Adar1^-/-^ mice

To better understand the cell-cell communications that mediate this response, we performed an inference analysis of cell-cell communication focused on receptor-ligand interactions (CellChat)^47^. This analysis revealed that with SMC *Adar1*^-/-^, SMC clusters SMC_cAdar1_KD1 and SMC_cAdar1_KD2 are notably high outward senders of CCL signaling markers (Fig 4G). When interrogating the specific signaling pairs, Ccl5:Ccr5 was the dominant interaction (Fig 4H). A circos plot depicting Ccl5:Ccr5 interaction strength revealed dominant outward interaction arising from SMC_cAdar1_KD1 and SMC_cAdar1_KD2 clusters with key receiving clusters being resident macrophage, infiltrating macrophage, and T cell populations (Fig 4I). Feature plots of Ccl5 and Ccr5 further revealed the distinct expression of Ccl5 within SMCs following *Adar1* KO and Ccr5 within macrophage populations (Fig 4J&K). Evaluation of the next top 3 interaction pairs (Ccl2:Ccr2; Ccl4:Ccr5, Ccl7:Ccr2) suggested that following Ccl5:Ccr5 interaction, Ccl2 and Ccl7 arising from SMC_cAdar1_KD2, Fibro_Sca1_hi, and macrophage cell populations appeared to interact with Ccr2 within macrophage and T lymphocyte populations (Fig S10G-L). These data suggested that Ccl4 from resident and infiltrating macrophage clusters further signaled through Ccr5 on these same cell clusters (Fig S10J&K). Activation of CCR5 has been previously identified as a key regulator of antiapoptotic signaling in macrophage cells in response to viral infection^48^. Our observation provides a unique insight that following the loss of SMC *Adar1*, Ccl5:Ccr5 acts as a dominant regulator of macrophage infiltration and macrophage cell survival.

### Homozygous deletion of MDA5 (Ifih1) prevents transcriptomic effect of SMC-Adar1 KO seen with single cell RNA sequencing

Our data indicates that loss of SMC *Adar1* elicits severe vascular effect and mortality due to dsRNA activation of MDA5 (Fig 3). Further, our *in vitro* HCASMC data suggests that loss of *ADAR1* with concurrent MDA5 KD produces only a minor effect on cellular transcriptome (Fig S2). However, it is unclear if this is true *in vivo* and there remains question as to if loss of RNA editing by ADAR1 may mediate significant transcriptomic effect despite concurrent MDA5 deletion. To investigate this, we performed additional scRNAseq captures in 9 *SMC-Adar1^-/-^, Mda5^-/-^* mice at this same timepoint of 14-days post tamoxifen as previously performed, for a total of 6 scRNAseq captures. Data was integrated together with control and *SMC-Adar1^-/-^* captures (16 total sequencing libraries). Remarkably, UMAP visualization of single cell transcriptomic data from *SMC-Adar1^-/-^, Mda5^-/-^* mice show nearly identical pattern compared to control (S11A). Infiltrating macrophage and T cell populations and Sca1-hi fibroblast populations are no longer present. When evaluating expression of key ISGs such as *Isg15* and *Ccl5*, homozygous *Mda5* (*Ifih1*) KO prevents the entirety of ISG induction across all cell types (S11B-C) and when analysis is focused on tdTomato clusters alone (S11D-E). We interrogated expression of specific SMC marker genes such as *Myh11* and *Cnn1* in tdTomato lineage traced cells, where we observe that downregulation of contractile markers seen with *SMC-Adar1* KO is prevented with *Mda5* KO (S11F-G), indicating this effect is due to activation of MDA5. *Findmarker* differential gene expression analysis and heatmap visualization of marker genes from the three genotypes show near identical patterns between control and *SMC-Adar1^-/-^, Mda5^-/-^* mice (S11H)(*Findmarker* analysis in Table S4). These data demonstrate most effect of *SMC-Adar1* KO is secondary to MDA5 activation. A small gene set where expression is observed to be downregulated in both *SMC-Adar1^-/-^* and *SMC-Adar1^-/-^, Mda5^-/-^* mice compared to control (S11H), suggesting that a small fraction of genes may be downregulated with loss of SMC Adar1 RNA editing independent of MDA5. A second set of genes appear to be downregulated with *SMC-Adar1^-/-^* however are increased with *SMC-Adar1^-/-^, Mda5^-/-^* (S11H), suggesting a role for MDA5 in regulating the expression of these genes independent of *Adar1*.

### SMC specific haploinsufficiency of Adar1 activates the ISG response in atherosclerosis and increases vascular chondromyoctye formation

Although conditional homozygous deletion of *Adar1* in SMCs caused a profound and rapid onset phenotype *in vivo* in a manner dependent on Mda5, we aimed to understand if haploinsufficiency of *Adar1* in SMCs, where mice appear to have no discernable phenotype, may lead to MDA5 activation, SMC phenotypic modulation, and progression of atherosclerosis. To do this, we used the *SMC-Adar1^-/+^* (*Adar1*^fl/WT^, *Myh11^CreERT2^*, *ROSAtdTomato*, *ApoE^-/-^*) mice with genotype controls (*Adar1*^WT/WT^, *Myh11^CreERT2^*, *ROSAtdTomato*, *ApoE^-/-^*), dosed with tamoxifen at 7.5 weeks of age, and placed on high fat diet for 16 weeks (Fig 5A). In this model, mice developed pronounced atherosclerosis as we have shown previously^32,33,44,45^. *SMC-Adar1^-/+^* and control mice had no differences in weight or lipid analysis (cholesterol, LDL, HDL, triglycerides) at this 16-week timepoint (Fig S12). To understand the cellular resolution of the vessel wall, at 16 weeks, we performed scRNAseq of the aortic root and ascending aorta. With a total of 12 scRNAseq captures analyzing over 70,000 cells comparing *SMC-Adar1^-/+^* and control genotypes, we observed expected distinct cell clusters of SMCs, fibroblasts, endothelial cells, macrophage cells, and T cells (Fig 5B). Lineage traced SMCs demonstrated distinct SMC clusters as we have previously observed and have been consistent with phenotypic modulation^33,44,45^, with a mature SMC cluster, a fibromyocyte (FMC) cluster, and a calcification promoting chondromyocyte (CMC) cluster of cells (Fig 5B). Importantly, *SMC-Adar1^-/+^* mice showed a large and distinct new SMC population residing between the mature SMC population and the FMC population (Fig 5B). Characterization of this population by *findmarker* analysis demonstrated this cluster to express numerous markers of ISG activation (i.e. *Ifit3*, *Ifit1*, *Isg15*) consistent with MDA5 activation. Analysis of *Isg15* as a marker of ISG activation revealed notable expression within this SMC cluster, termed ‘SMC_ISG’ (Fig 5B&C)(*Findmarker* results in Table S5).

**Figure 5.**
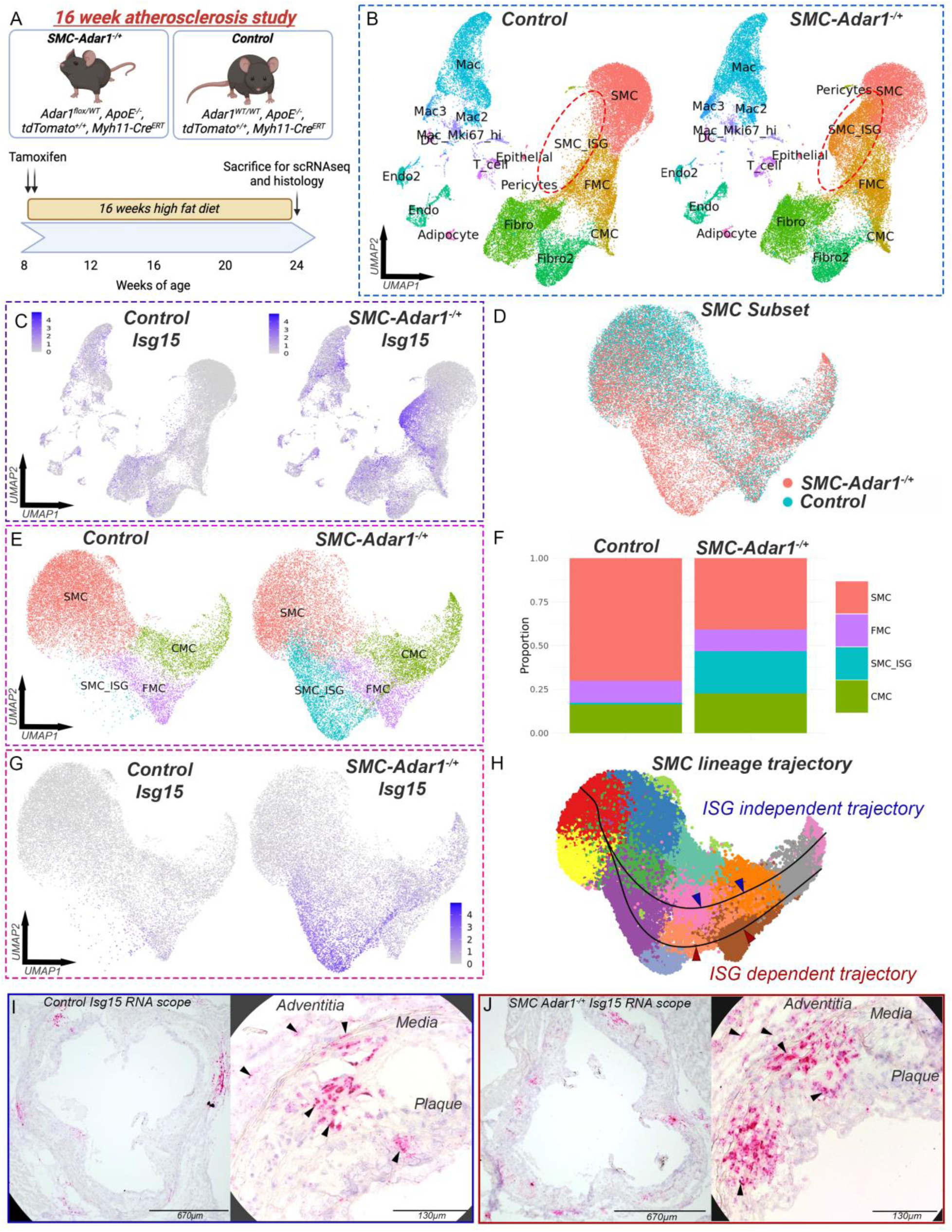
SMC specific haploinsufficiency of Adar1 activates the ISG response in atherosclerosis increasing vascular chondromyoctye formation. (A) Schematic diagram of 16 week high fat diet (HFD) atherosclerosis study. (B) UMAP of scRNAseq data from atherosclerotic aortic root and ascending aorta at 16 weeks high fat diet split by control (*Adar1*^WT/WT^, *Myh11^CreERT2^*, *ROSAtdTomato*, *ApoE^-/-^*) and *SMC-Adar1^-/+^* (*Adar1*^fl/WT^, *Myh11^CreERT2^*, *ROSAtdTomato*, *ApoE^-/-^*) genotypes. (C) Featureplot of *Isg15* split by genotype. (D) UMAP of SMC subset analysis grouped by genotype. (E) UMAP of SMC subset cluster analysis split by genotype showing SMC, SMC_ISG (interferon stimulated gene), fibromyocyte (FMC), and chondromyocyte (CMC) cell clusters. (F) Stacked bar chart showing cell populations of SMC subset for SMC, SMC_ISG, FMC, and CMC clusters. (G) Featureplot of *Isg15* in SMC subset split by genotype. (H) Lineage trajectory analysis using slingshot within SMC subset showing two distinct lineages (SMC_ISG dependent and independent) from SMC to CMC. (I&J) RNA scope for *Isg15* in aortic root sections in control (I) and *SMC-Adar1^-/+^* (J) mice.

To focus on the lineage traced SMC population, we performed an analysis on the ‘SMC’, ‘SMC_ISG’, ‘FMC’, and ‘CMC’ clusters. With subsequent dimensionality reduction and repeat clustering, UMAP visualization highlighted the marked effect of SMC *Adar1* haploinsufficiency in SMC phenotypic states in atherosclerosis (Fig 5D&E). Top 10 marker genes and GO term enrichment for ‘SMC’, ‘SMC_ISG’, ‘FMC’, and ‘CMC’ clusters reveals the SMC_ISG cluster to be marked by ISGs including *Ifit3, Isg15, Ifit1,* and *Irf7* and GO terms highlighting “response to virus”, “response to interferon-beta”, and “defense response to virus” (Fig S13A&B). Evaluation of SMC marker *Myh11* demonstrated downregulation of mature contractile markers with phenotypic modulation (Fig S14A) with associated upregulation of FMC and CMC markers (i.e. *Tnfrsf11b*, *Col2a1*, and *Acan*) (Fig S14B-D). Separation by genotype further highlighted that with *SMC-Adar1^-/+^* mice, there was a shift away from mature SMCs, marked increase in the SMC_ISG cluster (1.12% SMC_ISG in control vs 24.7% SMC_ISG in *SMC-Adar1^-/+^*; *p<E-100*), and an increase in CMC (16.3% CMC in control vs 22.6% CMC in *SMC-Adar1^-/+^*; *p=7.29E-100*)(Fig 5E&F). Comparison of the expression of CMC markers *Col2a1* and *Acan* between genotypes demonstrated an increase in CMC cells with *SMC-Adar1^-/+^* mice consistent with this finding (Fig S14E&F). This finding suggests that ISG activation from SMC *Adar1* haploinsufficiency in atherosclerosis may be contributing to CMC formation and vascular calcification.

### Lineage trajectory analysis of vascular chondrogenesis reveals a distinct ISG dependent trajectory

To investigate the potential for this SMC lineage relationship, we performed pseudotime inference and cellular trajectory analysis with Slingshot^49^. By increasing clustering resolution we generated 14 high resolution clusters of all tdTomato lineage traced SMCs and performed Slingshot inference to understand the cellular trajectory between mature SMC to CMC clusters. In this analysis, we observed only 2 distinct trajectories between SMC and CMC (Fig 5H). Importantly, one trajectory excludes all SMC_ISG clusters, while a second is dependent on SMC_ISG clusters (Fig 5H). This suggests two distinct cellular trajectories to CMC formation, with one trajectory dependent on ISG activation, while a second is ISG independent.

To better understand the top genes driving this ISG dependent trajectory, we performed *findmarker* analysis on all high-resolution clusters and evaluated the top markers for each cluster in sequence of the ISG dependent pathway (Fig S15A). This analysis highlights that as SMC lose expression of contractile markers such as *Myh11* and *Acta2*, they begin to express numerous ISGs. Initially, there is high expression of *Irf7*, *Isg15*, *Ifit1*, which is then followed by increasing expression of *Ly6a*, *C3*, *Cxcl12*, and other FMC markers such as *Dcn* and *Tnfrsf11b*, before SMCs have marked expression of key CMC genes such as *Col2a1*, *Chad*, *Fmod*, and *Spp1* (Fig S15A). To further understand the pathways driving this ISG dependent trajectory, we selected the gene markers of the high-resolution clusters that are distinct to the ISG trajectory and performed gene ontology pathway analysis (Fig S15B). Here, we identified top viral immune response pathways including *‘Coronavirus Disease’*, *‘Influenza A’*, and *‘Epstein-Barr Virus infection’*, driven by genes such as *Jun, Ccl2*, *Il6*, *Nfkb1*, and this analysis also reveals key JAK/STAT signaling molecules such as *Stat3* and *Jak1* (Fig S15B). In addition to immune pathways, *‘Focal Adhesion’* and *‘Regulation of Actin Cytoskeleton’* are highlighted, driven by genes such as *Rock1*, *Itgb1*, *Egfr,* PDGFs, and *Map2k1*, suggesting a distinct interrelationship between activation of JAK/STAT and key immune signaling pathways influencing focal adhesion pathways (Fig S15B).

To determine the location of MDA5 activation within the plaque and vessel wall, we performed RNA scope on aortic root sections for *Isg15*. In control mice, we observed scattered *Isg15* expression within the media, the base of the plaque and extending up to the region beneath the cap, as well as in the adventitia (Fig 5I). However, in *SMC Adar1^-/+^* mice we observed a marked increase in *Isg15* signal, with intense *Isg15* expression within the media, throughout the plaque, and within the adventitia (Fig 5J). Signal of *Isg15* RNAscope positive area in the vessel wall was quantified (Fig S16A) and location of *Isg15* signal was compared to signal of chondromyocyte marker *Col2a1*, where we reveal that cells that express *Isg15* reside in spatially distinct regions compared to *Col2a1* (Fig S16B-C). These data suggest ISG activation in atherosclerosis in control mice, however this effect is markedly increased with haploinsufficiency of *Adar1* within the vascular SMC.

### SMC specific haploinsufficiency in Adar1 increases plaque size and calcification in atherosclerosis

We performed histological analysis of the aortic root for *SMC-Adar1^-/+^* and control genotype mice after 16 weeks of high fat diet (n=13 each group). Using fluorescent imaging of tdTomato and autofluorescence (elastin), we quantified plaque area and plaque area normalized to total area of the internal elastic lamina area (IEL) (Fig 6A-D). We observed that *SMC-Adar1^-/+^* mice had a significant increase in plaque area, an effect that remained following normalization to overall area of the IEL (Fig 6A-D). Notably, quantification of the area of the media (area between IEL and external elastic lamina, EEL), we observed *SMC-Adar1^-/+^* mice to have a significant reduction, an effect that remained following normalization to the total area of the EEL (Fig 6E&F). When plaque area was normalized to media area, the observed effect with *SMC-Adar1^-/+^* mice was further enhanced (Fig 6G). In addition, we observed total area of the EEL to be increased (Fig 6H), suggesting an increased outward remodeling of the plaque. Given our observation from single cell data that suggested increased chondromyocyte formation, we evaluated the effect of SMC *Adar1* haploinsufficiency on calcification by Ferangi Blue Chromogen staining (Fig 6I-L). We observed *SMC-Adar1^-/+^* mice to have a significant increase in total Ferangi Blue staining by area (Fig 6K), an effect that remained with normalization to plaque area (Fig 6L). These findings demonstrate that SMC *Adar1* haploinsufficiency accelerate the formation atherosclerotic plaque area and calcification.

**Figure 6.**
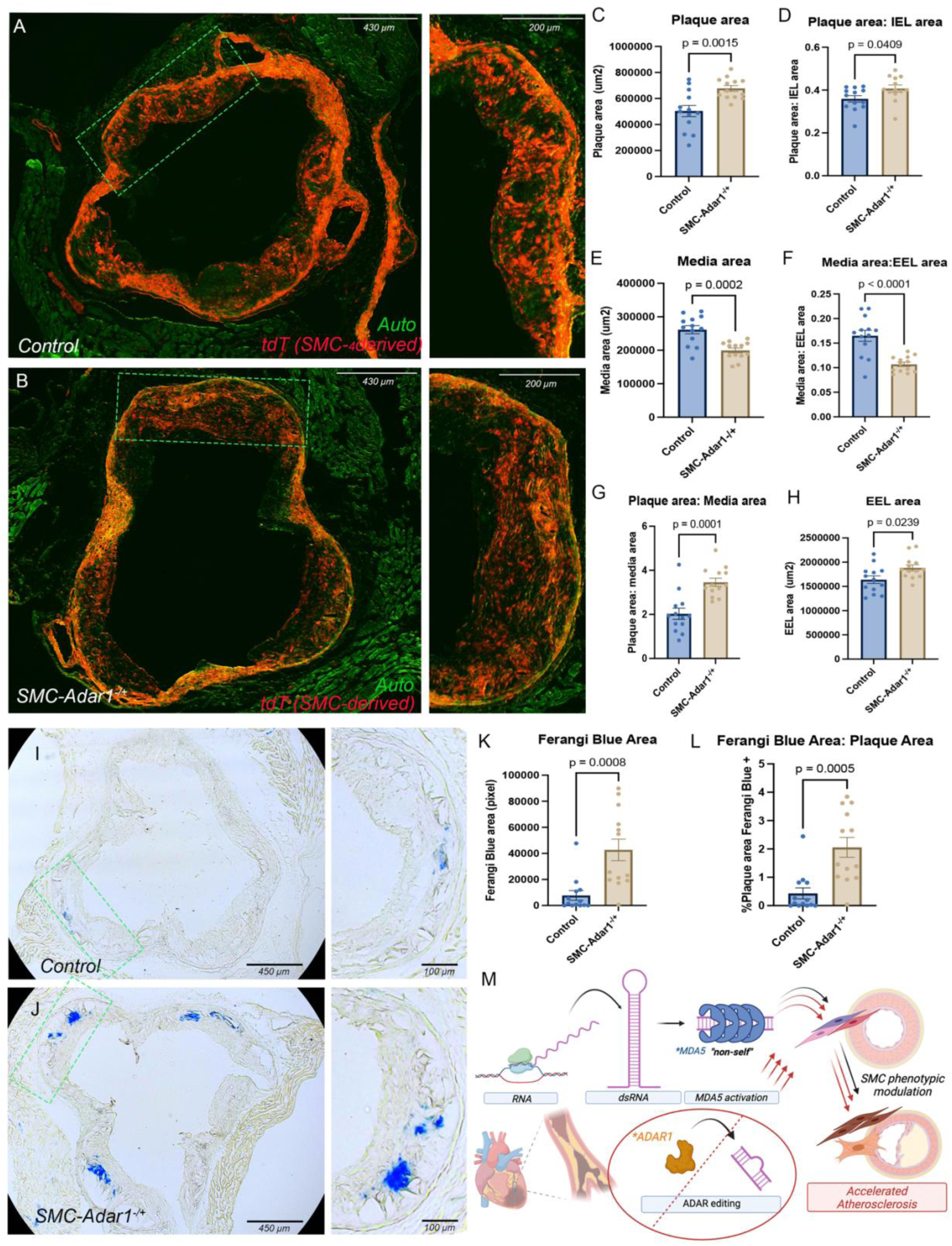
SMC specific haploinsufficiency in Adar1 increases plaque size and calcification in atherosclerosis. Representative fluorescent images of aortic root plaque histology in *SMC-Adar1^-/+^* and control genotypes (A&B) with quantification of plaque area (C), plaque area normalized to internal elastic lamina area (IEL) (D), media area (E), media area normalized to external elastic lamina area (EEL) (F), plaque area normalized to media area (G), and EEL area (H). Representative images of Ferangi Blue calcification staining in *SMC-Adar1^-/+^* and control genotypes (I&J), quantification of Ferangi Blue area (K), and Ferangi Blue area normalized to plaque area (L). (M) Schematic diagram showing mechanism of dsRNA formation, MDA5 activation, and impaired RNA editing by ADAR1 to accelerate SMC phenotypic modulation, calcification, and atherosclerosis. Data represent mean +/- SEM, N = 13 per group. *P-* values represent unpaired two-tailed T-test.

### SMC specific haploinsufficiency of Adar1 increased SMC lineage traced cell content in plaque without a change in acellular area

Our data suggests that impaired RNA editing in vascular SMC leads to increased plaque area and decreased media thickness, implicating increased SMC migration out of the media and into the plaque. To investigate this, we quantified tdTomato positive area within plaque as a marker of SMC lineage traced cells integrating into the plaque. Here, we observed that in *SMC Adar1^-/+^* mice, the percent tdTomato area within plaque is increased compared to control mice (p=0.0024)(Fig S17A-B). To then understand if this increased SMC migration into the plaque results in a similar increase in tdTomato area within the cap of the plaque, we measured the tdTomato area of the top 30μm segment of the cap (Fig S17D). Here, we did not observe a significant difference, although there was a greater mean tdTomato area in the SMC Adar1^-/+^ mice (p=0.0749).

To evaluate if there is increased acellular area within the plaque, we performed Masson’s Trichrome stains on lesions and quantified acellularity by tracing the plaque and subtracting all area that has stain to then be able to quantify all acellular area. Here, when normalizing acellular area by plaque area, we observed no difference between genotypes (Fig S17E-G).

### SMC specific haploinsufficiency in Adar1 has minimal effect on macrophage content in atherosclerosis

With a subsetted analysis on all non-tdTomato lineage traced cell populations, we observed that *SMC-Adar1^-/+^* mice have a small increase in total macrophage cell populations in this 16-week atherosclerosis model (39.7% vs 35.2% of all non-tdTomato cell population) (χ^2^ *p=3.66E-26*) with minimal change in cell numbers across all other cell types (Fig S18A&B). As we have shown that CCL signaling and Ccl5:Ccr5 appeared to be the dominant driver of macrophage recruitment in *SMC-Adar1^-/-^* mice, we investigated if these signaling molecules were similarly upregulated in this 16-week *SMC-Adar1^-/+^* atherosclerosis model. When performing receptor-ligand interaction analysis using CellChat^47^, we observed the ‘SMC_ISG’ cluster to have absent CCL signaling and did not appear to act as a sender, receiver, mediator, or influencer (Fig S18C&D). We then performed Cd68 immunohistochemical staining on aortic root sections to evaluate for Cd68 positive area within the plaque between control and *SMC-Adar1^-/+^* mice (Fig S18E-I). We observed that the total Cd68 positive area within the plaque was significantly greater in the *SMC-Adar1^-/+^* mice (Fig S18G), however this effect only trended towards significance when normalized to plaque area (Fig S18H) or area of the total vessel at the internal elastic lamina (Fig S18I). The lack of a major effect on macrophage content at this timepoint for this 16-week atherosclerosis *Adar1* haploinsufficiency model is in contrast to our 2-week SMC-*Adar1* KO model (Fig 2) and likely reflects a ‘dose-dependent’ response of SMC *Adar1* gene deletion, MDA5 activation, and *Ccl5* expression.

### Haploinsufficiency of Mda5 prevents ISG activation and chondromyocyte formation in atherosclerosis with SMC-Adar1^-/+^

Homozygous as well as heterozygous deletion of *Mda5* (*Ifih1*) prevents mortality seen with *SMC-Adar1* KO (Fig 3) and homozygous deletion of *Mda5* (*Ifih1*) prevents the entirety of ISG activation in our single cell RNA sequencing analysis (Fig S11), raising the question as to if deletion of *Mda5* can similarly prevent ISG induction and prevent CMC formation observed with atherosclerosis with *SMC-Adar1* haploinsufficiency. To investigate this, we further bred *SMC-Adar1^-/+^* mice onto *Mda5* (*Ifih1*) heterozygous background (*Adar1*^fl/WT^, *Ifih1^-/+^, Myh11^CreERT2^*, *ROSAtdTomato*, *ApoE^-/-^*) mice and performed additional 16-week HFD atherosclerosis studies following tamoxifen dosing at 7.5 weeks following our established protocol (Fig 5). We chose to use the *Mda5* (*Ifih1*) heterozygous background over homozygous deletion given haploinsufficiency is more akin to what may be achieved from future therapeutic inhibition. We performed additional single cell RNA sequencing on the aortic root and ascending aorta with 9 mice for a total of 6 additional scRNAseq captures. Following data integration with a total of more than 96K cells with >50,000 reads per cell (18 total scRNAseq libraries), we evaluated how *Mda5* haploinsufficiency affects cellular phenotype and lineage trajectory during atherosclerosis compared to control and *SMC-Adar1^-/+^* genotypes. With UMAP visualization, we demonstrate near identical overlap between the control and *SMC-Adar1^-/+^*, *Mda5^-/+^* genotypes (Fig 7A), suggesting Mda5 haploinsufficiency to largely prevent global transcriptomic effect of SMC *Adar1* haploinsufficiency. With evaluation of key ISGs such as *Isg15*, we observe that *Mda5* haploinsufficiency prevents the entirety of ISG induction (Fig 7B). With SMC subset analysis and visualization of SMC clusters (‘SMC’, ‘SMC_ISG’, ‘FMC’, and ‘CMC’) we observe a clear effect where *SMC-Adar1^-/+^*, *Mda5^-/+^* genotype eliminates the ‘SMC_ISG’ cluster seen with *SMC-Adar1^-/+^* mice (Fig 7C), with further reduction in CMC formation (Fig 7C). Upon evaluation of key marks of SMC, SMC_ISG, FMC, and CMC populations, we observe that *Mda5* haploinsufficiency decreases the expression of CMC markers such as *Col2a1* and *Acan* (Fig 7D-E) with near elimination of expression of ‘SMC_ISG’ marker *Isg15* (Fig 7F). In quantifying the proportion of cells that belong to each SMC cluster in the control, *SMC-Adar1^-/+^*, and *SMC-Adar1^-/+^*, *Mda5^-/+^* genotypes, we observe that the control and *SMC-Adar1^-/+^* genotypes have 17% and 24% CMCs whereas the *SMC-Adar1^-/+^*, *Mda5^-/+^* genotype significantly reduces this proportion to 12% (Fig 7G). This significant reduction in CMC population in the *SMC-Adar1^-/+^*, *Mda5^-/+^* genotype is significantly lower than not only the *SMC-Adar1^-/+^* genotype, but also the control genotype (Fig 7G). This may suggest that even in control genotype, Mda5 is being activated and contributing to CMC formation. Differential *findmarker* gene expression analysis in the tdTomato population and heatmap visualization shows most marker genes increased in *SMC-Adar1^-/+^* genotype compared to control are further decreased in the *SMC-Adar1^-/+^*, *Mda5^-/+^* genotype, suggesting the gene expression effect is dependent on Mda5 (Fig 7H). However, similar to our observation in the SMC *Adar1* KO + *Mda5* KO study (Fig S11), a portion of genes appear to be regulated by Adar1 independent of Mda5 and similarly a portion of genes appear to be regulated by Mda5 independent of Adar1 (Fig 7H). Upon evaluation of top ISGs (ie. *Isg15*, *Ifit3*, *Irf7*, *Stat1*), we observe a consistent pattern of increased expression in *SMC-Adar1^-/+^* genotype compared to control which is then blocked in entirety with the *SMC-Adar1^-/+^*, *Mda5^-/+^* genotype (Fig 7I). Importantly, when evaluating key markers of SMC phenotypic transition and FMC/CMC formation (ie. *Spp1*, *Fn1*, *Tnfrsf11b*, *Dcn*) we see a baseline moderate level of expression in the control group, a significant increase in the *SMC-Adar1^-/+^* genotype, and then a decrease in expression to a level lower than the control in the *SMC-Adar1^-/+^*, *Mda5^-/+^* genotype (Fig 7J). This is strong evidence to suggest that Mda5 is playing a significant role in regulating SMC phenotypic transition in atherosclerosis even with genetically intact *Adar1* RNA editing, consistent with our noted finding of increasing ISG activation with SMC phenotypic transition in both human and mouse in atherosclerosis (Fig 1) and human genetics data.

**Figure 7.**
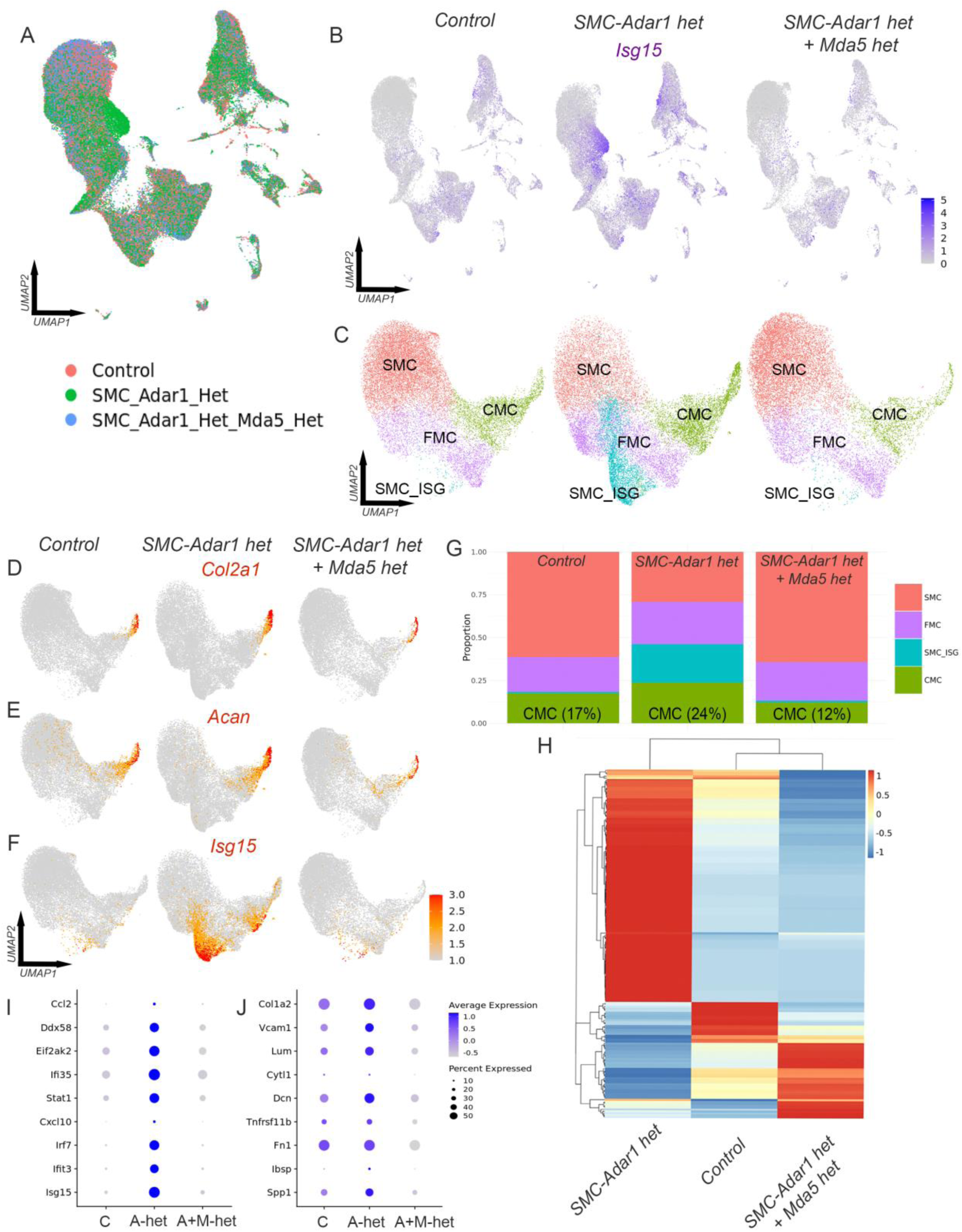
Haploinsufficiency of Mda5 prevents ISG activation and chondromyocyte formation in atherosclerosis with SMC-Adar1^-/+^. (A) UMAP of integrated scRNAseq data set between control (*Adar1^WT/WT^, Myh11^CreERT2^, ROSAtdTomato, ApoE^-/-^*), *SMC-Adar1 het* (*Adar1^flWTl^, Myh11^CreERT2^, ROSAtdTomato, ApoE^-/-^*), and *SMC-Adar1 het + Mda5 het* (*Adar1^flWTl^, Ifih1^-/+^, Myh11^CreERT2^, ROSAtdTomato, ApoE^-/-^*) genotypes following 16 weeks high fat diet. (B) Featureplot of *Isg15* split by genotype. (C) UMAP tdTomato+ subset analysis with SMC, SMC_ISG, FMC, and CMC clusters split by genotype. Featureplots split by genotype for (D) *Col2a1*, (E) *Acan*, and (F) *Isg15*. (G) Stacked bar charts showing proportion of cells within ‘SMC’, ‘SMC_ISG’, ‘FMC’, and ‘CMC’ clusters split by genotype. (H) Heatmap visualization for normalized expression of marker genes for control, *SMC-Adar1 het*, and *SMC-Adar1 het + Mda5 het* genotypes in tdTomato+ subset analysis. Dotplot gene expression splity by genotype in SMC subset for (I) ISG genes and (J) FMC/CMC marker genes.

We measured the overall expression of *Ifih1* (Mda5) across genotypes where featureplot visualization and mean normalized expression values show a pattern of increasing expression of *Ifih1* (Mda5) with *SMC-Adar1^-/+^* (1.5 fold increase for all cells, 3.6 fold increase for SMC) (Fig S19). We observed a 65% and 58% reduction in *Ifih1* (Mda5) expression in *SMC-Adar1^-/+^*, *Mda5^-/+^* genotype compared to control in all cells and SMC subset respectively (Fig S19). This data suggests that Mda5 haploinsufficiency not only produces a ∼50-60% reduction in expression but prevents an upregulation seen with *SMC-Adar1^-/+^*.

### ISG expression correlates to markers of SMC phenotypic modulation, plaque vulnerability, and plaque calcification in large human atherosclerosis dataset (Athero-Express)

Our group’s prior work has revealed edQTLs that reduce ADAR1 RNA editing to correlate with increased ISG activation in human atherosclerotic plaque^12^, and our current analysis of human scRNAseq datasets have suggested the vascular SMC to have a unique requirement for ADAR1 RNA editing and that ISG activation in SMCs occurs with SMC phenotypic modulation (Fig 1). Although our comprehensive human *in vitro* and murine *in vivo* models suggest a novel mechanism where impaired ADAR1 editing leads to altered SMC phenotype driving increasing atherosclerosis and calcification through activation of MDA5 in an ‘axis’ of ADAR1-dsRNA-MDA5, there remains a question as to if this relationship between ISG activation, adverse plaque phenotype, and atherosclerotic plaque calcification remains true in larger human populations. To address this question, we utilized our unique human database of carotid endarterectomy samples (Athero-Express) where 1093 patient samples underwent bulk RNA sequencing and paired with detailed histological characterization of plaque and clinical phenotypic information (Fig 8A)^50^. Here, we evaluated the expression of key ISGs, markers of SMC phenotypic modulation/calcification, and related their expression to plaque phenotype.

**Figure 8.**
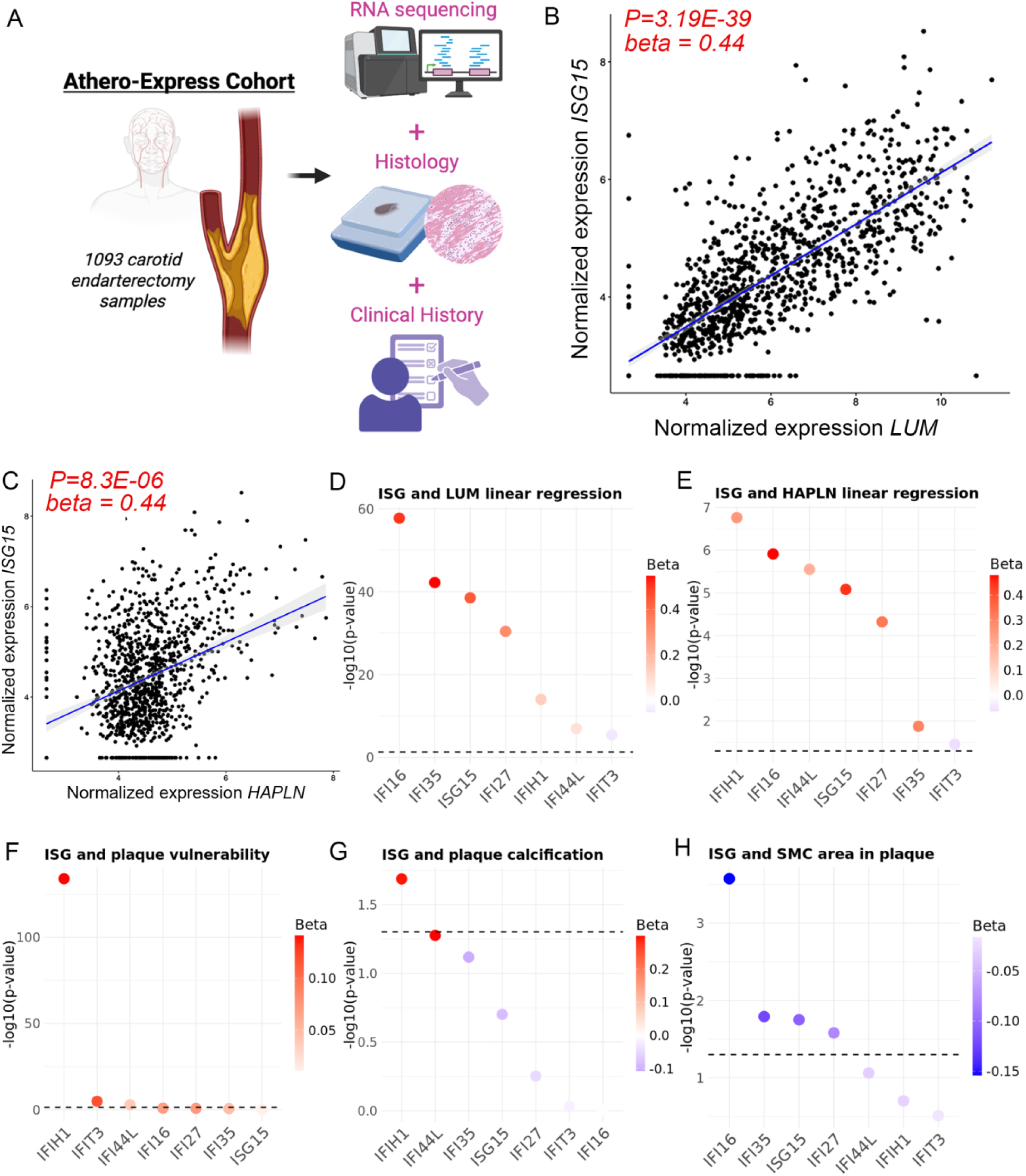
ISG expression in human atherosclerotic plaque dataset (Athero-Express) correlates to markers of SMC modulation, plaque vulnerability, plaque calcification, and decreasing SMC investment into plaque. (A) Schematic diagram of Athero-Express cohort and data evaluation. Linear relationship analysis between chondromyocyte markers *LUM* (B) and *HAPLN* (C) to *ISG15*. Dot plots displaying linear regression analysis between ISGs and *LUM* (D) and *HAPLN* (E) showing -log10(p value) on Y axis and gene on X axis with beta coefficient scaled by color. Dot plots displaying linear regression analysis between ISGs and plaque phenotype corrected for co-variates for (F) plaque vulnerability, (G) calcification, and (H) SMC area within plaque.

The patient demographics for this dataset has been previously reported^50^, briefly, the patients enrolled are predominantly male (75.3%), mean age is 68.4yrs, and the majority of patients have had symptomatic carotid stenosis (43.2% with stroke, 24.2% with transient ischemic attack, 17.3% with ocular symptoms) with only 15.3% asymptomatic. We first evaluated the distribution of RNA expression for 7 key ISGs identified from our prior scRNAseq datasets (*ISG15*, *IFIH1*, *IFIT3*, *IFI16*, *IFI27*, *IFI35*, *IFI44L*). We revealed a significant distribution of expression ranging from very low to quite elevated with a similar level of distribution between these 7 ISGs (i.e. *IFIH1* and *ISG15*, Fig S20A-B). We evaluated the expression between men and women by normalized patient density by RNA expression and show there is no difference in expression between sexes (i.e. *IFIH1* and *ISG15*, Fig S20C-D). To understand the relationship in expression between these ISGs, we performed correlation and heat map analysis and showed an overall positive correlation between ISGs with an unanticipated exception of *IFIT3*, where *IFIT3* expression showed a slight negative correlation to other ISGs (Fig S20E).

To understand the relationship of ISG expression and CMC formation we performed linear regression analysis between individual ISGs and CMC markers *LUM* and *HAPLN* (Fig 8B-E). We observed a strong positive relationship between expression of ISGs and *LUM* and *HAPLN*, where, for example, the expression of *ISG15* showed a significant linear relationship to *LUM* with a beta of 0.44 (*P=3.19E-39*) (Fig 8B) with a similar relationship to *HAPLN* (Fig 8C). When evaluating the linear regression P-value and beta coefficient across each ISG, we observe *IFI16* to have the strongest relationship to *LUM* while *IFIH1* to have the strongest relationship to *HAPLN* (Fig 8D&E) with a consistent pattern of increasing ISGs to have a significant linear relationship to CMC markers. There was a notable exception of *IFIT3* that had a weak yet significant negative relationship to *LUM* and *HAPLN*, perhaps suggesting distinct biology for *IFIT3* (Fig 8D&E).

Given this relationship of ISG expression to CMC markers, we aimed to understand if ISG expression correlates to plaque histological phenotype. Following the adjustment for co-variates including age, sex, year of surgery, HTN, DM2, smoking, LDL-C, lipid lowering and anti-platelet therapies, eGFR, DMI, history of CVD (see methods), we evaluated the correlation of ISG expression to overall plaque vulnerability (0-5 score), plaque calcification, total SMC area within the plaque, and total macrophage area within the plaque. We observed that expression of ISGs *IFIH1*, *IFI44L*, and *IFIT3* had a significant positive relationship to plaque vulnerability (Fig 8F). Notably, increasing expression of *IFIH1* was associated with increased vulnerability of plaque with a p-value of significance of *9.71E-135* (Fig 8F). We observed a similar positive relationship between expression of *IFIH1* and measure of plaque calcification (*P=0.02*) and a trend of significance for *IFI44L* (*P=0.052*) (Fig 8G).

A notable observation from our *in vivo* data shows that MDA5 activation can lead to ISG activation and SMC cell death in the SMC *Adar1* homozygous KO model with macrophage recruitment into the vessel wall, whereas the SMC *Adar1* heterozygous model in atherosclerosis can lead to increased SMC recruitment into plaque with minimal effect on macrophage recruitment. To understand how ISG activation relates to SMC and macrophage investment into plaque in human disease, we evaluated the relationship of ISG expression and the overall SMC and macrophage area within plaque. Intriguingly, we observed a significant negative relationship between ISG expression and SMC investment into plaque, where increasing expression of *ISG15*, *IFI35*, *IFI16*, and *IFI27* were each significantly associated with decreasing SMC plaque area (i.e. *IFI16 P=2.6E-4*)(Fig 8H). Importantly, we found ISG expression to have little relationship to macrophage area within plaque, however, we did observe a significant negative relationship between *IFI44L* expression and macrophage area within plaque (Fig S20F).

These data together show that ISG activation in human plaque correlates to increasing SMC phenotypic modulation and CMC formation, increasing plaque vulnerability and calcification, while leading to decreased SMC investment within plaque with minimal effect on macrophage recruitment.

## DISCUSSION

Both academic and industry leaders are leveraging the power of human genetics to drive discovery in the goal to define causal disease biology^51^. The rapid pace of identifying novel genomic loci that meet genome wide level significance for complex disease has resulted in notable challenges of target prioritization within a locus^52^, as well as prioritization for therapeutic drug development^53^. However, the association of edQTL and CAD risk, where impaired RNA editing increases CAD risk while increasing ISG activation, implicates a common disease pathway that is largely dependent on one gene product — MDA5. Strikingly, 20% of CAD risk loci examined colocalized with edQTL loci^12^. With a new understanding of the genetic determinants of RNA editing and dsRNA sensing as an underlying etiology of coronary artery disease and other inflammatory disorders^12^, there is a significant need to define how this mechanism influences disease risk.

Here, we provide a crucial link between the human genetics that implicate RNA editing and dsRNA sensing by MDA5 in atherosclerosis and the biological underpinnings of this disease association. Analysis of human atherosclerotic plaque scRNAseq data indicate a pattern where SMCs express immunogenic RNA at high levels and that ISG activation — suggestive of MDA5 activation with inadequate RNA editing — occurs with SMC phenotypic modulation in both human and mouse data. We further show that deletion of SMC specific *Adar1* in adult mice results in loss of vascular integrity and significant mortality due to activation of MDA5 (Figs 2 and 3). Further, scRNAseq analysis of aortic tissue reveals distinct ISG activation and a complex transcriptional response and inflammatory cell recruitment within the vessel wall (Fig 4), an effect blocked in its near entirety with deletion of MDA5 (Fig S11). This highlights the absolute requirement for *Adar1* within the vascular SMC, but its role appears to be primarily to prevent dsRNA activation of MDA5. Importantly, in 16-week atherosclerosis studies in *Adar1* haploinsufficient mice, scRNAseq reveals ISG activation in SMCs in a cell type and context specific mechanism driving phenotypic modulation, vascular chondromyocyte formation, and to a lesser extent macrophage recruitment (Fig 5), and that this coincides with increased plaque area, decreased thickness of the media, and increased calcification (Fig 6). This effect of ISG activation and chondromyocyte formation in SMC *Adar1* haploinsufficiency is prevented with only haploinsufficiency of Mda5, suggesting the importance of Mda5 in mediating this effect (Fig 7). To corroborate this finding in human datasets, we utilized the Athero-Express cohort of carotid endarterectomy samples where patient data with careful histological examination are paired with bulk RNA sequencing in 1093 patients^50^. Impressively, ISG expression are positively associated with increasing plaque vulnerability, increasing plaque calcification, and decreased SMC investment into the plaque, likely indicative of ISG activation contributing to SMC cell death within the plaque (Figs 8 & S20).

These findings highlight an important new concept, that the need for RNA editing to evade MDA5 activation not only occurs in a cell type specific manner, but a cellular context specific manner. Two prior studies have investigated the role of SMC *Adar1* in haploinsufficiency in SMC phenotypic modulation, focused on carotid injury^42^ and an angiotensin II model of abdominal aortic aneurysm (AAA)^41^. In these reports, the authors reported that loss of SMC Adar1 prevented phenotypic modulation, however no investigation into potential MDA5 activation was performed. Our data reported here appears in contrast to this finding, however our findings implicate that the effect of loss of ADAR1 on phenotypic transition is dependent on MDA5 activation and occurs in a ‘dose dependent’ mechanism where increasing MDA5 activation has a greater effect. In the SMC *Adar1^-/+^* model, there is no immediate activation of MDA5, however in specific cellular contexts immunogenic RNA formation appears to ‘max out’ the RNA editing capacity of the single *Adar1* allele leading to MDA5 activation. This effect is overt in a 16-week high fat diet model of atherosclerosis; however, this was perhaps not evident in either of the prior short term carotid injury or AAA models. However, a recent discovery that MDA5 (*IFIH1*) is a locus in a large GWAS of AAA raises an intriguing hypothesis that ADAR1 and MDA5 are causal mechanisms in AAA formation^54^.

Our data indicate that SMC phenotypic modulation is a cellular context that requires heightened RNA editing and that deficient RNA editing with SMC *Adar1* heterozygous mice leads to MDA5 activation that accelerates SMC phenotypic changes and disease state. Although our data suggests that expression of *ADAR1* increases slightly with SMC phenotypic modulation *in vitro* and in atherosclerosis (Figs S1 and S5), this likely reflects inadequate compensatory response leading to an overall accumulation of dsRNA and MDA5 activation in specific cell states, influencing SMC phenotype. These data reveal trajectories to SMC formation of the vascular chondromyocyte (CMC), where ISG activation through MDA5 facilitates the formation of CMCs and vascular calcification. This effect of MDA5 activation on SMC phenotypic state is consistent with prior work investigating the role of ADAR1 and MDA5 on cellular proliferation and cell death^13^. ADAR1, through its role of editing dsRNA and preventing MDA5 activation, is crucial in regulating proliferation and deterring endogenous dsRNA from triggering transcriptional shutdown and apoptotic cell death through activation of PKR (*EIF2AK2*)^13^. It was previously reported that with initial KD of *ADAR1*, cell proliferation was found to be increased, however with mounting MDA5 activation there was ultimately PKR activation and transcriptional shut down and cell death^13^. This effect, where initial loss of ADAR1 stimulates proliferation and cell state transitions, is perhaps consistent with modifying a cell state similar to SMC to FMC transition in atherosclerosis in our SMC *Adar1^-/+^* model. However, with overwhelming MDA5 activation as seen in our SMC *Adar1^-/-^* model, there is transcriptional shut down, cell death, and major vascular effect. This biology may explain why in our human atherosclerosis dataset ISG expression was associated with decreasing SMC investment into plaque (Fig 8). This is somewhat in contrast to our *in vivo* SMC *Adar1^-/+^* model of atherosclerosis where we see ISG activation leading to increased SMC migration into the plaque, however this may reflect the countering biology where overt ISG activation can precipitate cell death and plaque instability.

Importantly, although classical inflammatory mechanisms have been implicated in the pathogenesis of atherosclerosis and tremendous interest has been placed in the potential to inhibit these pathways to treat disease^55^, only a small minority of GWAS loci in CAD implicate inflammatory pathways^2–5^. This finding raises the question as to the overall weight of effect for disease causality for such inflammatory mediators (i.e. IL6, TNFα, and IL1β). The relatively minimal benefit of IL1β antagonism in CAD^56^ may suggest an overall limited potential to treat disease through inhibition of these inflammatory mediators. However, dsRNA sensing by MDA5 and ISG activation as an overarching mechanism provides a new avenue of understanding as to how inflammatory mechanisms are driving disease biology. Our data demonstrates that ISG activation occurs in SMCs during atherogenesis, however the exact function of these ISGs remains unclear. The function of ISGs has been previously reviewed^57^, however, ultimately IFN and ISGs serve key roles to regulate cellular viral immune response, likely through driving transcriptional shutdown in a PKR dependent mechanism of cells infected by a viral pathogen. Importantly, Type I interferon induces JAK/STAT intracellular signaling pathways^57^, a finding we observed in our ISG dependent cellular trajectory in SMC phenotypic transition (Fig 5H & Fig S15). This finding suggests that JAK/STAT is an important regulator of this trajectory. Specific ISGs have distinct roles in fighting viral infection, for example on inhibiting viral entry (IFITM1/2/3), mRNA synthesis (IFI16), protein synthesis (PKR), and mRNA degradation (ISG20)^57^. The mechanism by which specific ISGs may modify atherosclerosis within the vessel wall remains unclear. However, an important observation from human genetics is that edQTL loci, which ultimately implicates dsRNA formation, MDA5 activation, and ISG induction, to be a greater driver of disease heritability than all eQTLs in CAD^12^. This data presented here further implicates a specific mechanism of disease while highlighting a unique opportunity to leverage this biology for therapeutic development by prioritizing the inhibition of dsRNA sensing and downstream activation of MDA5.

## MATERIALS AND METHODS

### Data availability

All raw and processed sequencing data are deposited in the National Center for Biotechnology Information Gene Expression Omnibus. Bulk and single cell RNA sequencing data is available through accession IDs GSE254862, GSE255661, and GSE280641.

### Mouse Models

To generate SMC-specific deletion of *Adar* we used the *Adar^flox/flox^* allele (exon 7-9 floxed; MGI allele: Adar^tm1.1Phs^; MGI:3828307) as we have reported previously^58^, crossed onto the validated SMC-specific mouse Cre allele (*Myh11*^CreERT2^) combined with the floxed tandem dimer tomato (tdT) fluorescent reporter protein gene knocked into the ROSA26 locus (B6.Cg-Gt(ROSA)26Sortm14(CAGtdTomato)Hze/J) and *ApoE*^-/-^ alleles (*Adar*^flox/flox^, *Myh11*^CreERT2^, *ROSAtdT*/+, *ApoE*^-/-^). To induce homozygous or heterozygous deletion of *Adar1* in SMCs (*SMC-Adar1^-/-^*: *Adar1*^fl/fl^, *Myh11^CreERT2^*, *ROSAtdTomato*, *ApoE^-/-^* or *SMC-Adar1^-/+^*: *Adar1*^fl/WT^, *Myh11^CreERT2^*, *ROSAtdTomato*, *ApoE^-/-^*), mice receive 2 doses of tamoxifen (200mg/kg, P.O.) at 7.5-8 weeks of age (dose 1 - day 0; and dose 2 - day 2) and started on a high fat diet as we have done previously.^44,45^ To evaluate the effect of MDA5 KO on *Adar1* KO background, we crossed the *Ifih1*^-/-^ mouse (B6.Cg-*Ifih1^tm1.1Cln^*/J) that we have used previously^58^, onto the above background in a heterozygous manner to generate SMC *Adar1*^-/-^ x Mda5^-/+^ mice (Adar^flox/flox^, *Ifih1*^-/+^, Myh11^CreERT2^, ROSAtdT/+, ApoE^-/-^), SMC *Adar1*^-/-^ x Mda5^-/-^ mice (*Adar*^flox/flox^, *Ifih1*^-/-^, *Myh11*^CreERT2^, *ROSAtdT*/+, *ApoE*^-/-^), or SMC *Adar1*^-/+^ x MDA5^-/+^ mice (*Adar*^flox/WT^, *Ifih1*^-/+^, *Myh11*^CreERT2^, *ROSAtdT*/+, *ApoE*^-/-^).

### Mouse aortic root/ascending aorta cell dissociation

Immediately after sacrifice, mice were perfused with phosphate buffered saline (PBS). The aortic root and ascending aorta were excised, up to the level of the brachiocephalic artery. Tissue was washed three times in PBS, placed into an enzymatic dissociation cocktail (2 U ml−1 Liberase (5401127001; Sigma–Aldrich) and 2 U ml−1 elastase (LS002279; Worthington) in Hank’s Balanced Salt Solution (HBSS)) and incubated at 37 °C for 45min, and subsequently minced. The cell suspension was further dissociated with 1ml pipette 20X and single cell suspension was confirmed under microscope. Cells were then pelleted by centrifugation at 500g for 5 min. The enzyme solution was then discarded, and cells were resuspended in fresh HBSS and passed through a 35um filter into a FACS tube. To increase biological replication, a single mouse was used to obtain single-cell suspension, and three mice were used in combination for each scRNA capture. 3 separate pairs of isolation were performed for control, *SMC-Adar1^-/+^*, *SMC-Adar1^-/-^*, *SMC-Adar1^-/-^*, *Mda5^-/-^,* or *SMC-Adar1^-/+^*, *Mda5^-/+^* mice. Cells were FACS sorted based on tdTomato expression. tdT+ cells (considered to be of SMC lineage) and tdT− cells were then captured on separate but parallel runs of the same scRNA-Seq workflow (gating strategy and threshold identical to those published in previous work by Wirka et al^33^), and datasets were later combined for all subsequent analyses.

### Mouse aortic root histology

IHC was performed according to standard protocol. Primary antibodies for CD68 rabbit polyclonal antibody (1:400 dilution; ab125212; Abcam) with secondary rabbit-on-rodent HRP polymer (RMR622; BioCare Medical) for CD68 evaluation. Hematoxylin and eosin (H&E) and Trichrome staining was performed using standard protocols using and H&E and Trichrome Stain Kits (Abcam, ab245880 and ab150686 respectively). The processed sections were visualized using a Leica DM5500 microscope objective magnifications, and images were obtained using Leica Application Suite X software. For quantification of atherosclerosis plaque area, sections obtained at equal distance measured from the superior margin of the coronary sinus were used for comparison to ensure equal level sections were obtained. Images with FITC autofluorescence and tdTomato with red channel representing SMC lineage traced cells were obtained. Areas of interest were quantified using ImageJ (National Institutes of Health) software and compared using a two-sided t-test. Lesion size was defined by the area encompassing the intimal edge of the lesion to the border of SMC-marker positive intima-media junction. Calcification was evaluated using Ferangi Blue Chromogen Kit 2 (Biocare Medical, FB813) following manufacturers protocol. For RNAscope, slides were processed and hybridized according to the manufacturer’s protocol, with reagents from ACD Bio. All area quantification was performed in a genotype blinded fashion with image J using length information embedded in exported files. All biological replicates for each staining were performed simultaneously on position-matched aortic root sections to limit intra-experimental variance.

### Single-cell capture and library preparation and sequencing

All single-cell capture and library preparation was performed in the Quertermous lab, sequencing was performed by Medgenome Inc (Foster City, CA). Cells were loaded into a 10x Genomics microfluidics chip and encapsulated with barcoded oligo-dT-containing gel beads using the 10x Genomics Chromium controller according to the manufacturer’s instructions. Single-cell libraries were then constructed according to the manufacturer’s instructions (Illumina). Libraries from individual samples were multiplexed into one lane before sequencing on an Illumina platform with targeted depth of 50,000 reads per cell for RNA or 400 million paired end reads per sequencing library.

### Analysis of scRNA-Seq data

Fastq files from each experimental time point and mouse genotype were aligned to the reference genome (mm10) individually using CellRanger Software (10x Genomics). The dataset was then analyzed using the R package Seurat 30^32^ and datasets were merged in Seurat using standard Merge function. The dataset was trimmed of cells expressing fewer than 2000 genes, and genes expressed in fewer than 50 cells. The number of genes, number of unique molecular identifiers and percentage of mitochondrial genes were examined to identify outliers. As an unusually high number of genes can result from a ‘doublet’ event, in which two different cell types are captured together with the same barcoded bead, cells with >9000 genes were discarded. Cells containing >7.5% mitochondrial genes were presumed to be of poor quality and were also discarded. The gene expression values then underwent library-size normalization and normalized using established LogNormalize function in Seurat. No additional batch correction was performed. Principal component analysis was used for dimensionality reduction, followed by clustering in principal component analysis space using a graph-based clustering approach via Louvain algorithm. UMAP was then used for two-dimensional visualization of the resulting clusters. Pseudotime trajectory analysis was performed using Slingshot^49^ with available software. Analysis, visualization and quantification of gene expression and generation of gene modulescores were performed using Seurat’s built-in function such as “FeaturePlot”, “VlnPlot”, “AddModuleScore”, and “FindMarker.” Inference analysis of cell-cell communication was performed using CellChat^47^. Standard CellChat protocol was used where overexpressed genes, interactions, and communication probability was computed and subsequently aggregated and visualized across cell clusters.

For immunogenic RNA score, we generated an AddModuleScore ‘score’ that summates the expression of all ‘immunogenic’ RNA from a putative list that was generated by combining our data from experimental *in vitro* models^27^ along with human genetic edQTL analysis^12^ and complete list of immunogenic RNA is included in Table S1.

### HCASMC culture and *in vitro* genomic studies

Cells were cultured in smooth muscle growth medium (Lonza; catalog number: CC-3182) supplemented with human epidermal growth factor, insulin, human basic fibroblast growth factor and 5% FBS, according to the manufacturer’s instructions. All HCASMC lines were used at passages 4–8. HCASMCs immortalized with hTert transgene were acquired from Dr. Clint Miller (UVA) were used for all calcification assays with HCASMCs between passages 31 and 35. siRNA knockdowns were performed using Lipofectamine RNAiMax (Life Technologies) using manufacturer’s recommended protocol at 50pg siRNA / 100,000cells.

### Phenotypic transition, calcification, and TGF***β*** assays

For phenotypic transition assays, HCASMC are cultured in 5% BSA and then subjected to serum starvation for 2 days. Following 2 days of serum starvation (0% BSA) HCASMCs develop a mature SMC phenotype, reintroduction of serum (5%) stimulates a more proliferative phenotype and RNA/protein is isolated 48hrs following reintroduction of serum. For calcification assays, HCASMCs are cultured in 1% BSA with 10mM beta-glycerophosphate and 8mM CaCl2 for 7 to 10 days. For TGF*β* stimulation assays, we cultured HCASMC to confluency, treated with siRNA for 48 hours, and transitioned cells to a serum free condition for 72 hours. We then stimulated cells with TGF*β* (10ng/mL) or control for 72 hours before we isolated RNA and performed quantitative PCR.

### RNA isolation and Bulk RNA sequencing

RNA was isolated from disrupted HCASMC cells using standard RNeasy mini kit protocol (Qiagen). Isolated RNA underwent quality assessment, cDNA conversion, and library construction by Novogene (San Jose, CA). cDNA libraries were sequenced using Illumina Novoseq 6000 achieving 20-30 million paired-end reads per sample. RNAseq data were processed using a workflow starting from fastq files and code is available on github (https://github.com/zhaoshuoxp/Pipelines-Wrappers/blob/master/RNAseq.sh). In brief, the quality control was performed using FastQC, mapping to the human genome hg19/GRCh37 was performed using STAR second pass mapping to increase the percentage of mapped reads, and counting was done with featureCounts using GENCODE gtf annotation. Next, differential analysis was performed using DESeq2.

### RNA editing analysis

RNA editing analysis in bulk RNA seq data was performed following a pipeline as we have done previously^12^. Briefly, we compiled a list of reference editing sites for quantification. By incorporating known sites in the RADAR database^59^, tissue-specific sites identified in GTEx V6p17, and recently published hyper-editing sites^60^, we finalized a list of 2,802,572 human editing sites. To quantify the editing levels, we computed the ratio of G reads divided by the sum of A and G reads at each site. Because the bulk RNA-seq data from HCASMC cells was not strand specific, the reads cannot be automatically assigned to sense and antisense strands, so that the quantification of editing level may be affected by expression of either of the two overlapping genes. Therefore, we developed a method to measure the RNA editing level of a given site using the reads derived from the corresponding transcript (sense for A-to-G edits and antisense for T-to-C edits), with less influence by the other overlapping transcript. For reads that are edited, they would be derived from the sense transcript if the edits are A-to-G, and from the antisense transcript if the edits are T-to-C. For the rest of the reads that are unedited, we estimated the numbers derived from the sense versus antisense transcripts in proportion to their expression levels. Overall RNA editing frequency was reported as percentage of total RNA editing sites across the genome.

### Athero-Express Bulk RNA isolation/sequencing and histological staining and analysis

Data from the Athero-Express Biobank study were used for the ISG expression validation in human carotid endarterectomy samples. The AE is an ongoing biobank study including endarterectomy patients from two tertiary hospitals in the Netherlands. Patients provided written informed consent, the medical ethics committee of the respective hospitals approved of the study (protocol number 22/018), and the study adheres to the Declaration of Helsinki^61^. A total of 1,093 plaque segments were selected from two subsequent experiments, Athero-Express RNA Study 1 and 2 (n = 622 and n = 471, respectively). RNA was isolated using in-house standardized protocols. Library preparation utilized the CEL-seq2 method. This method yielded the highest mappability reads to the annotated genes. It captures the 3’ end of polyadenylated RNA species and includes unique molecular identifiers (UMIs), which allows for direct counting of the unique RNA molecules. This plaque sample processing for RNA isolation and library preparation has previously been described in detail^50^. Both AERNAS1 and AERNAS2 where mapped against the cDNA reference of all transcripts in GRCh38.p13 and Ensembl 108 (GRCh38.p13/ENSEMBL_GENES_108). These include raw read counts of all non-ribosomal, protein coding genes with existing HGNC gene name. All read counts are corrected for UMI sampling by ‘raw.genecounts=round(-4096*(log(1-(raw.genecounts/4096))))’.

For histological analysis, stained slides are digitally stored up to 40X magnification. Semi-quantitatively scoring was performed by two independent observers, and a third independent observer if interpretations between the first two differed. Multiple stains were used to phenotype each plaque segment. Collagen content was assessed using picrosirius red and elastin von Gieson staining. Smooth muscle cells were identified using alpha actin, and macrophages using CD68 staining. Hematoxylin–Eosin (H&E) staining was used to evaluate calcification and the lipid core. In combination with Fibrin (Mallory’s phosphotungstic acid-hematoxylin), H&E was also used to detect the presence of intraplaque haemorrhage (IPH). The scoring protocol of the semi quantitative scoring has been described previously^61,62^. The alpha actin (smooth muscle cell) and CD68 (macrophage) are also analysed quantitatively by computerized analysis. To assess plaque vulnerability, we use the plaque vulnerability index (PVI), a histological score incorporating collagen, macrophages, smooth muscle cells, lipid content and intraplaque haemorrhage^61^. Each trait was scored as stable or unstable, yielding a cumulative vulnerability score from 0 to 5.

The multivariable linear regression analyses between the ISG Expression and plaque characteristics were performed in R (version 4.4.1) using the ‘lm()’function to fit a linear model. The bulk RNA-Seq counts of the ISG genes were transformed using variance stabilization transformation. For each ISG gene and phenotype (smooth muscle cells, macrophages, calcification, and plaque vulnerability index), we modelled the gene expression against the phenotype and corrected for age, gender, hypertension, diabetes and smoker status, LDL cholesterol levels, use of lipid-lowering drugs, use of antiplatelets drugs, estimated glomerular filtration rate based on the MDRD formula, body mass index, history of cardiovascular diseases, level of stenosis, and year of surgery. For the linear regression analysis between the ISG expression and CMC markers (*LUM & HAPLN*). The bulk RNA-Seq counts of both the ISG genes and CMC markers were transformed using variance stabilization transformation. We modelled each ISG gene against both CMC markers and corrected for Age, Sex and Year of surgery.

## List of Supplementary Materials

### Supplemental Figures

- Supplemental Figure 1. *SMC ADAR1 and MDA5 expression in atherosclerosis in mice*
- Supplemental Figure 2. *ADAR1 regulates RNA editing in human coronary SMCs and RNA editing requirement is dependent on cell context*.
- Supplemental Figure 3. *Loss of ADAR1 in human coronary artery SMCs (HCASMCs) regulates viral sensing transcriptomic response in vitro*
- Supplemental Figure 4. Global RNA editing is decreased in HCASMC following phenotypic modulation in vitro
- Supplemental Figure 5. *Serum and TGF β stimulation upregulate ADAR1 expression with differential effect on ISG induction in HCASMC*.
- Supplemental Figure 6. *Loss of ADAR1 regulates transcriptomic and inflammatory response to calcification in vitro*
- Supplemental Figure 7. *Quantification of histological findings for control and SMC-Adar1 KO mice*
- Supplemental Figure 8. *Aortas from SMC Adar1 KO with Mda5 KO are indistinguishable from control*
- Supplemental Figure 9. *SMC Adar1 KD induces antiviral defense transcriptional pathways*
- Supplemental Figure 10. *Loss of SMC Adar1 coordinates distinct response throughout vessel wall*
- Supplemental Figure 11. *Homozygous deletion of MDA5 (Ifih1) prevents transcriptomic effect of SMC-Adar1 KO*
- Supplemental Figure 12. *SMC Adar1 haploinsufficiency has no effect on weight or cholesterol in atherosclerosis high fat diet model*
- Supplemental Figure 13. *SMC specific haploinsufficiency of Adar1 in SMC subset analysis shows ISG activation with phenotypic modulation in atherosclerosis*
- Supplemental Figure 14. *SMC specific haploinsufficiency of Adar1 increases vascular chondromyoctye formation*
- Supplemental Figure 15. ISG dependent trajectory analysis from SMC to CMC implicates distinct gene ontologies
- Supplemental Figure 16. *Isg15 RNAscope reveals increased Isg15 signal in plaque of SMC Adar1^-/+^ mice*
- Supplemental Figure 17. *SMC specific haploinsufficiency in Adar1 increased SMC lineage traced cell content in plaque without change in acellular area*
- Supplemental Figure 18. *SMC specific haploinsufficiency of Adar1 has minimal effect on macrophage infiltration in atherosclerosis*
- Supplemental Figure 19. *Mda5 haploinsufficiency reduces Mda5 expression and prevents upregulation in SMC Adar1 het background*.
- Supplemental Figure 20. *Variability of ISG expression between patient carotid endarterectomy samples with no difference between sexes in Athero-Express cohort*.

## Author Contributions

Conceptualization: C.W., Q.L., P.C., J.B.L., and T.Q.

Methodology: C.W., Q.L., J.M.

Investigation: C.W., Q.L., J.M., H.G., D.G., W.G., M.W., Q.Z., M.R., D.L., B.P., A.B., R.K., T.N., T.P., M.M., C.M., S.L.

Visualization: C.W.

Funding acquisition: C.W., J.B.L., T.Q.

Project administration: C.W., J.B.L., T.Q.

Supervision: J.B.L., T.Q.

Writing — original draft: C.W., J.B.L., T.Q.

Writing — review and editing: C.W., Q.L., J.B.L., T.Q.

