## Supplemental Figures for "Smooth muscle expression of RNA editing enzyme *ADAR1* controls activation of RNA sensor MDA5 in atherosclerosis"

- Supplemental Figure 1. *SMC ADAR1 and MDA5 expression in atherosclerosis in mice*
- Supplemental Figure 2. *ADAR1 regulates RNA editing in human coronary SMCs and RNA editing requirement is dependent on cell context.*
- Supplemental Figure 3. *Loss of ADAR1 in human coronary artery SMCs (HCASMCs) regulates viral sensing transcriptomic response in vitro*
- Supplemental Figure 4. Global RNA editing is decreased in HCASMC following phenotypic modulation in vitro
- Supplemental Figure 5. *TGFβ attenuates ISG induction in HCASMC.*
- Supplemental Figure 6. *Loss of ADAR1 regulates transcriptomic and inflammatory response to calcification in vitro*
- Supplemental Figure 7. *Quantification of histological findings for control and SMC-Adar1 KO mice*
- Supplemental Figure 8. *Aortas from SMC Adar1 KO with Mda5 KO are indistinguishable from control*
- Supplemental Figure 9. *SMC Adar1 KO induces antiviral defense transcriptional pathways*
- Supplemental Figure 10. *Loss of SMC Adar1 coordinates distinct response throughout vessel wall*

- Supplemental Figure 11. *Homozygous deletion of MDA5 (Ifih1) prevents transcriptomic effect of SMC Adar1 KO*
- Supplemental Figure 12. *SMC Adar1 haploinsufficiency has no effect on weight or cholesterol in atherosclerosis high fat diet model*
- Supplemental Figure 13. *SMC specific haploinsufficiency of Adar1 in SMC subset analysis shows ISG activation with phenotypic modulation in atherosclerosis*
- Supplemental Figure 14. *SMC specific haploinsufficiency of Adar1 increases vascular chondromyoctye formation*
- Supplemental Figure 15. ISG dependent trajectory analysis from SMC to CMC implicates distinct gene ontologies
- Supplemental Figure 16. *Isg15 RNAscope reveals increased Isg15 signal in plaque of SMC Adar1*$^{-/+}$ *mice*
- Supplemental Figure 17. *SMC specific haploinsufficiency in Adar1 increased SMC lineage traced cell content in plaque without change in acellular area*
- Supplemental Figure 18. *SMC specific haploinsufficiency of Adar1 has minimal effect on macrophage infiltration in atherosclerosis*
- Supplemental Figure 19. *Mda5 haploinsufficiency reduces Mda5 expression and prevents upregulation in SMC Adar1 het background.*
- Supplemental Figure 20. *Variability of ISG expression between patient carotid endarterectomy samples with no difference between sexes in Athero-Express cohort.*

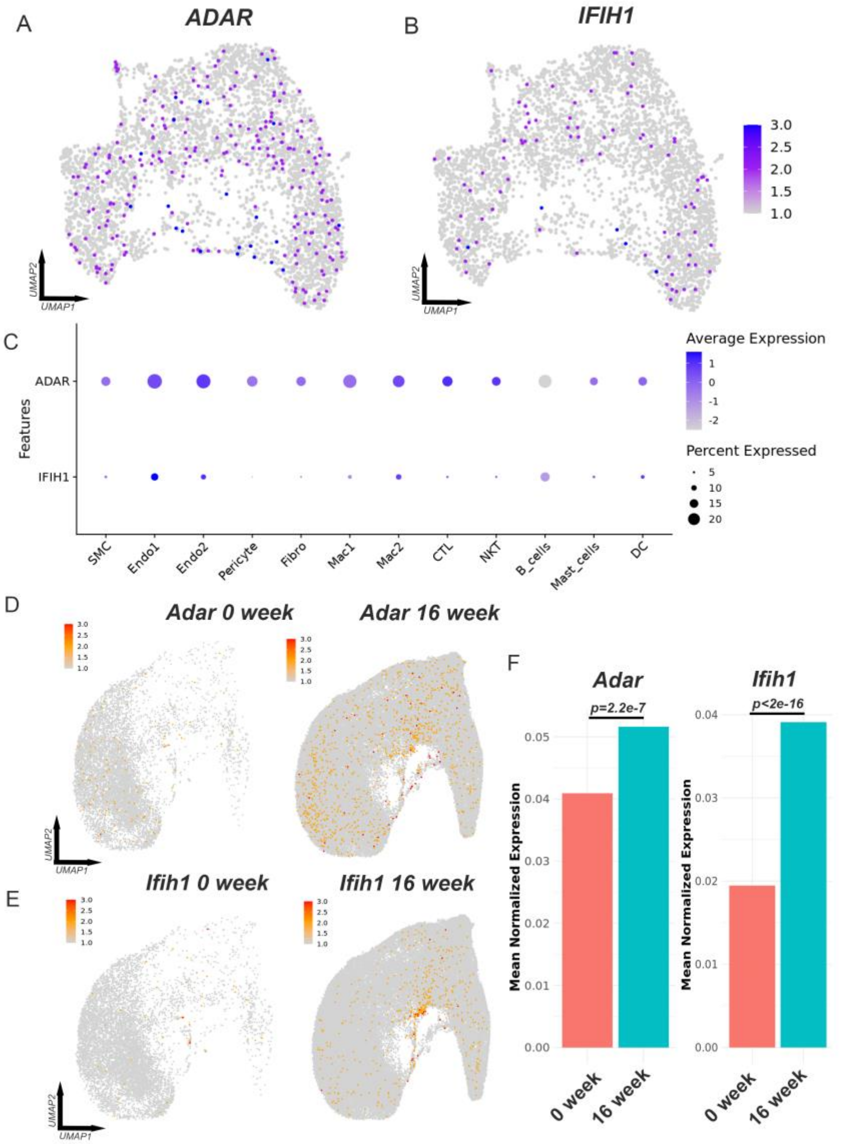


**Supplemental Figure 1.** ***SMC ADAR1 and MDA5 expression increase in atherosclerosis.*** Feature plot of (A) *ADAR* and (B) *IFIH1* in SMC subset analysis of human carotid atherosclerotic plaque (Alsaigh et al., 2022). (C) Dotplot showing expression of *ADAR* and *IFIH1* across celltypes in human carotid plaque dataset. (D-E) In mouse comprehensive integrated dataset of lineage traced SMCs (from Sharma et al., 2024), featureplot of *Adar* (D) and *Ifih1* (E) at 0 and 16 weeks of high fat diet. (F) Bar chart of extracted normalized expression of *Adar* and *Ifih1* in lineage traced SMCs at 0 and 16 weeks high fat diet.

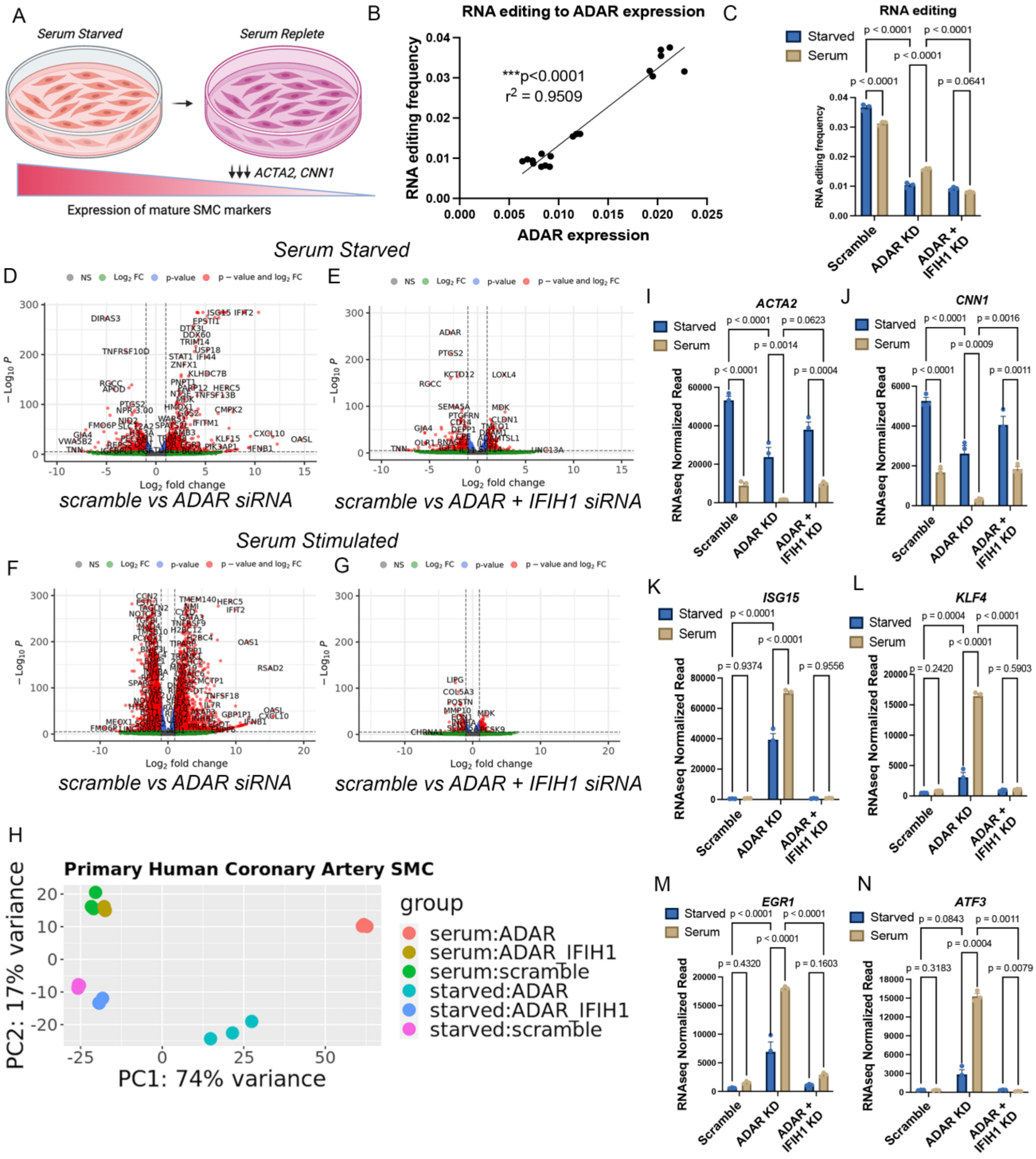


**Supplemental Figure 2.** ***ADAR1 regulates RNA editing in human coronary SMCs and RNA editing requirement is dependent on cell context.*** (A) Schematic of SMC phenotypic modulation in vitro assay. (B) Linear regression analysis of ADAR1 expression and global RNA editing frequency in primary human coronary artery SMCs (HCASMCs) following siRNA KD of *ADAR*. (C) Grouped bar chart of RNA editing frequency with cellular treatment of scramble, *ADAR*, and *ADAR* + *IFIH1* (MDA5) siRNA. (D) Volcano plot of bulk RNAseq DE gene analysis comparing scramble vs

*ADAR* siRNA in serum starved conditions, (E) volcano plot comparing scramble vs *ADAR* + *IFIH1* siRNA in serum starved conditions, (F) volcano plot comparing scramble vs *ADAR* siRNA in serum fed conditions, and (G) volcano plot comparing scramble vs *ADAR* + *IFIH1* siRNA in serum fed conditions. (H) Principal component analysis (PCA) of bulk RNAseq data of principal components 1 and 2 for each HCASMC treatment. Grouped bar chart of normalized RNAseq reads across HCASMC treatments for (I) *ACTA2*, (J) *CNN1*, (K) *ISG15*, (L) *KLF4*, (M) *EGR1*, and (N) *ATF3*. *P*-values represent simple linear regression (B) or (C, I-N) two-way ANOVA with multiple comparisons post hoc analysis. N = 3 RNA seq libraries per condition.

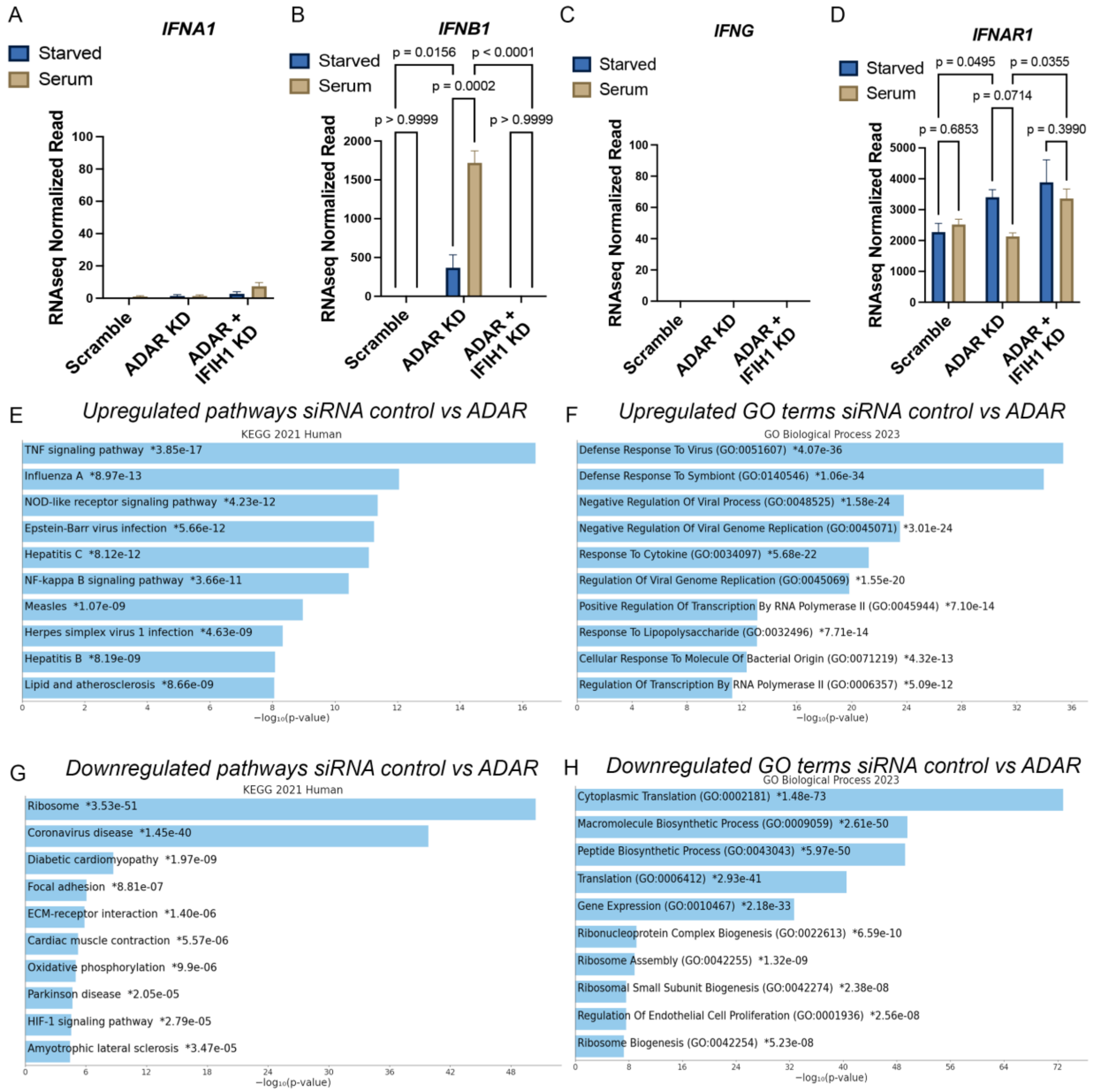


**Supplemental Figure 3.** ***Loss of ADAR1 in human coronary artery SMCs (HCASMCs) regulates viral sensing transcriptomic response in vitro*****.** Grouped bar chart of normalized RNAseq reads across primary HCASMC treatments for (A) *IFNA1*, (B) *IFNB1*, (C) *IFNG*, and (D) *IFNAR1*. KEGG pathways (E) and GO terms (F) of top 1000 upregulated genes with siRNA control vs *ADAR* KD. KEGG pathways (G) and GO terms (H) of top 1000 downregulated genes with siRNA control vs *ADAR* KD. *P*-values represent two-way ANOVA with multiple comparisons post hoc analysis.

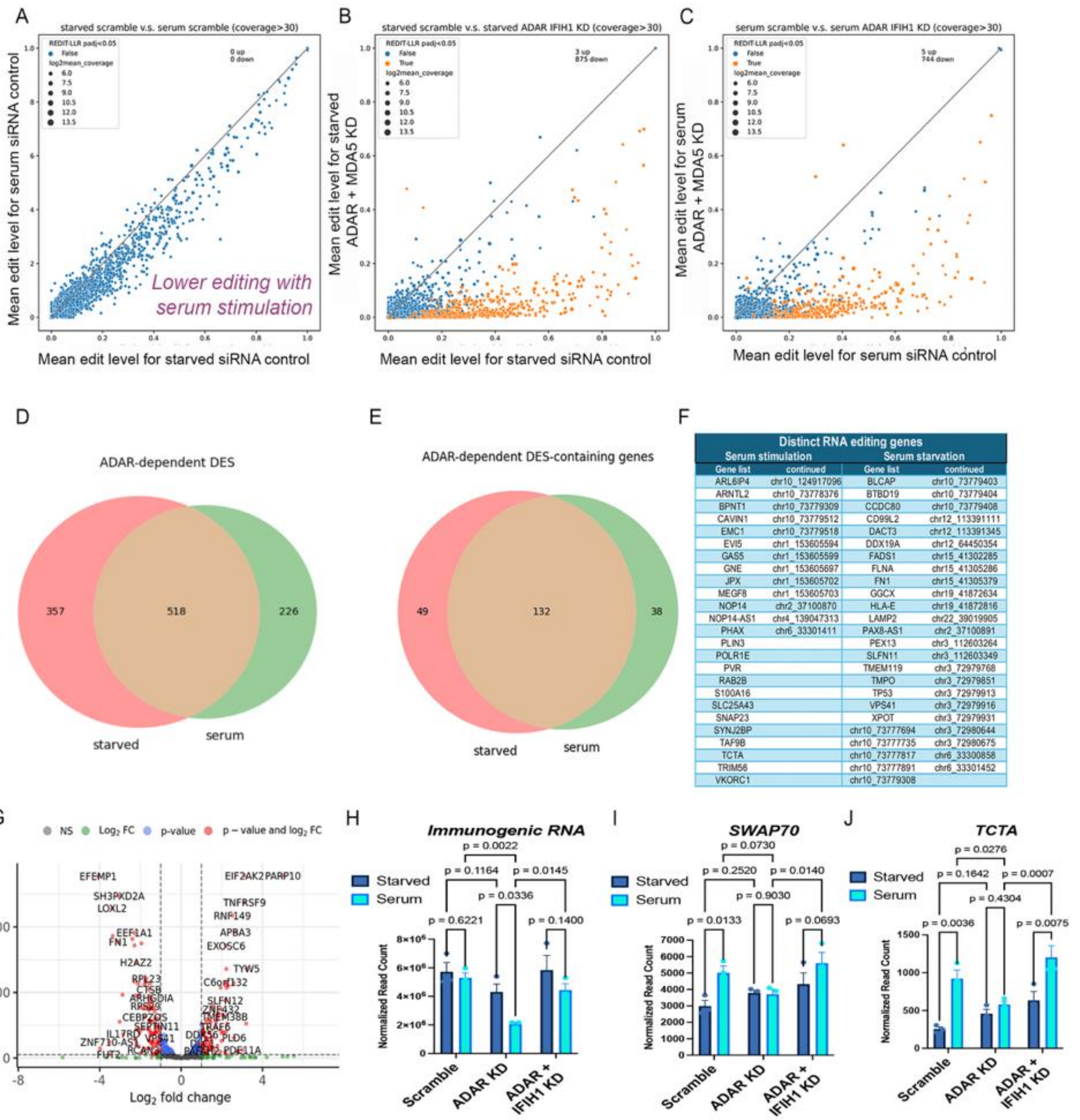


| Distinct RNA editing genes | | | |
|---|---|---|---|
| Serum stimulation | | Serum starvation | |
| Gene list | continued | Gene list | continued |
| ARL6IP4 | chr10_124917096 | BLCAP | chr10_73779403 |
| ARNTL2 | chr10_73778376 | BTBD19 | chr10_73779404 |
| BPNT1 | chr10_73779309 | CCDC80 | chr10_73779408 |
| CAVIN1 | chr10_73779512 | CD99L2 | chr12_113391111 |
| EMC1 | chr10_73779518 | DACT3 | chr12_113391345 |
| EVI5 | chr1_153605594 | DDX19A | chr12_64450354 |
| GAS5 | chr1_153605599 | FADS1 | chr15_41302285 |
| GNE | chr1_153605697 | FLNA | chr15_41305286 |
| JPX | chr1_153605702 | FN1 | chr15_41305379 |
| MEGF8 | chr1_153605703 | GGCX | chr19_41872634 |
| NOP14 | chr2_37100870 | HLA-E | chr19_41872816 |
| NOP14-AS1 | chr4_139047313 | LAMP2 | chr22_39019905 |
| PHAX | chr6_33301411 | PAX8-AS1 | chr2_37100891 |
| PLIN3 | | PEX13 | chr3_112603264 |
| POLR1E | | SLFN11 | chr3_112603349 |
| PVR | | TMEM119 | chr3_72979768 |
| RAB2B | | TMPO | chr3_72979851 |
| S100A16 | | TP53 | chr3_72979913 |
| SLC25A43 | | VPS41 | chr3_72979916 |
| SNAP23 | | XPOT | chr3_72979931 |
| SYNJ2BP | | chr10_73777694 | chr3_72980644 |
| TAF9B | | chr10_73777735 | chr3_72980675 |
| TCTA | | chr10_73777817 | chr6_33300858 |
| TRIM56 | | chr10_73777891 | chr6_33301452 |
| VKORC1 | | chr10_73779308 | |

**Supplemental Figure 4.** ***Global RNA editing is decreased in HCASMC following phenotypic modulation in vitro*.** In HCASMC, editing site specific comparison of RNA editing frequency for siRNA control in serum starved (X axis) vs serum stimulated (Y axis)(A). Editing site specific comparison of RNA editing frequency for siRNA control versus siRNA *ADAR* + *IFIH1* (MDA5) in serum starved (B) and serum stimulated (C) conditions. Venn diagram of ADAR-dependent differentially edited sites (DES) (D) and DES-containing genes (E). Table of distinct ADAR-dependent DES-containing genes for serum stimulated and serum starved conditions (F). Volcano plot of DE gene analysis for immunogenic RNA between scramble siRNA treated cells under serum starved and serum stimulated conditions (G). Grouped bar charts of normalized RNAseq reads for (H) all immunogenic RNA, (I) *SWAP70*, and (J) *TCTA*.

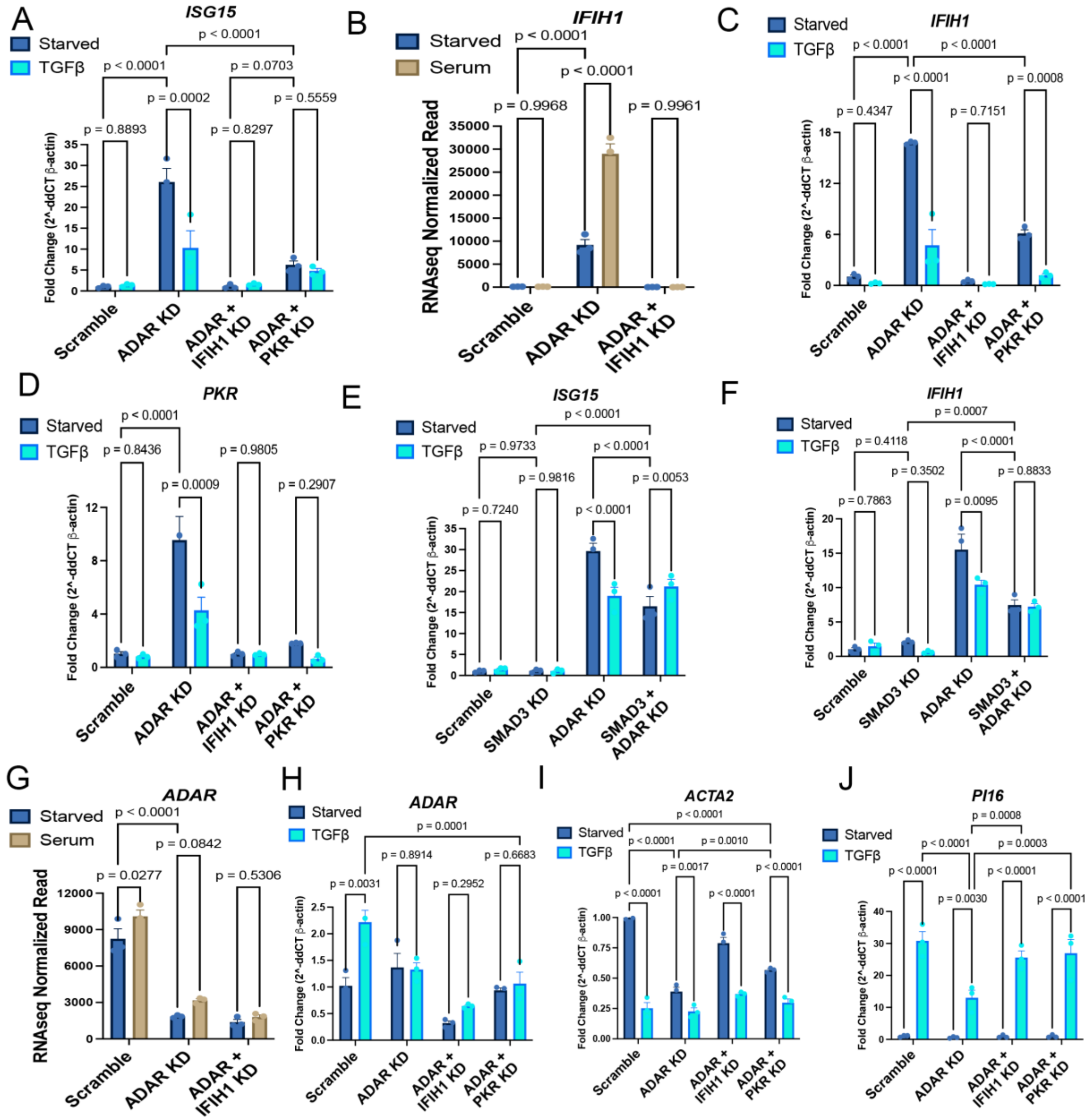


**Supplemental Figure 5.** ***TGFβ attenuates ISG induction in HCASMC.*** Grouped bar charts of normalized RNAseq reads or fold change expression in relation to $\beta$-actin by qPCR for (A) *ISG15* with *TGFβ* stimulation (10ng/mL 72 hours), (B) *IFIH1* in serum stimulated vs starved conditions, (C) *IFIH1* with *TGFβ* stimulation, (D) *PKR* with *TGFβ* stimulation, (E) *ISG15* with *TGFβ* stimulation and *SMAD3* KD, (F) *IFIH1* with *TGFβ* stimulation and *SMAD3* KD, (G) *ADAR* in serum stimulated vs starved conditions, (H) *ADAR* with *TGFβ* stimulation (E) *ACTA2* with *TGFβ* stimulation, and (H) *PI16* with *TGFβ*. *P*-values represent two-way ANOVA with multiple comparisons post hoc analysis. N = 3 per group.

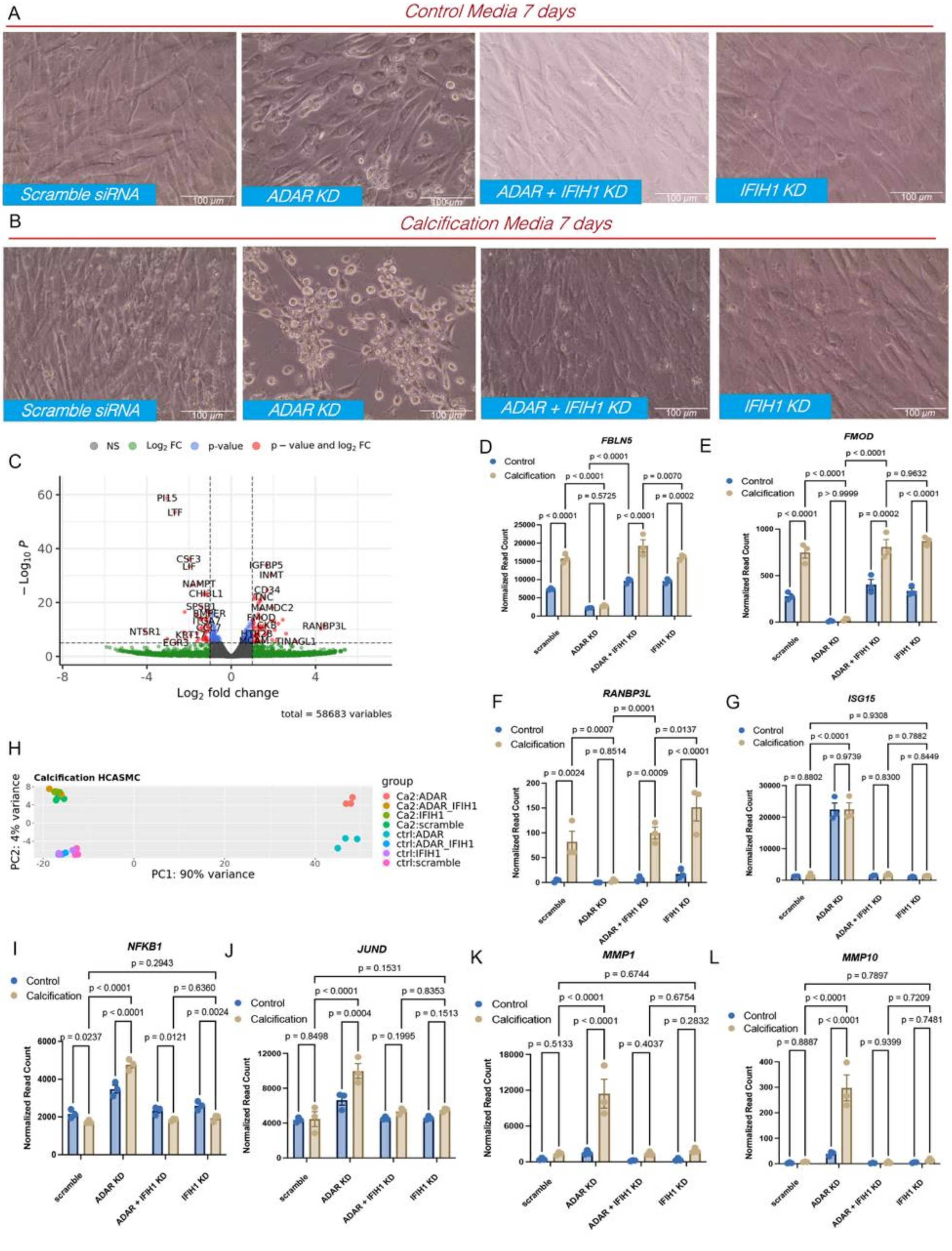
A
Control Media 7 days
Scramble siRNA
ADAR KD
ADAR + IFIH1 KD
IFIH1 KD
B
Calcification Media 7 days
Scramble siRNA
ADAR KD
ADAR + IFIH1 KD
IFIH1 KD
C
NS Log2 FC p-value p – value and log2 FC
PI15
LTF
CSF3
LIF
NAMPT
CHI3L1
IGFBP5
INMT
CD34
TNC
MAMDC2
FMOD
RANBP3L
NTSR1
EGR3
TINAGL1
Log2 fold change
total = 58683 variables
D
FBLN5
E
FMOD
Control
Calcification
Normalized Read Count
scramble
ADAR KD
ADAR + IFIH1 KD
IFIH1 KD
F
RANBP3L
G
ISG15
H
Calcification HCASMC
PC1: 90% variance
PC2: 4% variance
group
Ca2:ADAR
Ca2:ADAR_IFIH1
Ca2:IFIH1
Ca2:scramble
ctrl:ADAR
ctrl:ADAR_IFIH1
ctrl:IFIH1
ctrl:scramble
I
NFKB1
J
JUND
K
MMP1
L
MMP10

**Supplemental Figure 6. *Loss of ADAR1 regulates transcriptomic and inflammatory response to calcification in vitro*.** Images of hTert immortalized human coronary artery SMCs following treatment with scramble, *ADAR*, *ADAR* + *IFIH1*, and *IFIH1* siRNA and 7 days of culture in control medium (A) and calcification medium (B). (C) Volcano plot of DE gene analysis between scramble siRNA treated cells under control and calcification medium. (D-G) Grouped bar charts of normalized RNAseq reads for (D) *FBLN5*, (E) *FMOD*, (F) *RANBP3L*, (G) *ISG15*. (H) Principal component analysis (PCA) for PC1 and PC2 of bulk RNAseq data from each cell treatment. Grouped bar charts of normalized RNAseq reads for (I) *NFKB1*, (J) *JUND*, (K) *MMP1*, (L) *MMP10*. *P*-values represent two-way ANOVA with multiple comparisons post hoc analysis. N = 3 RNA seq libraries per condition.

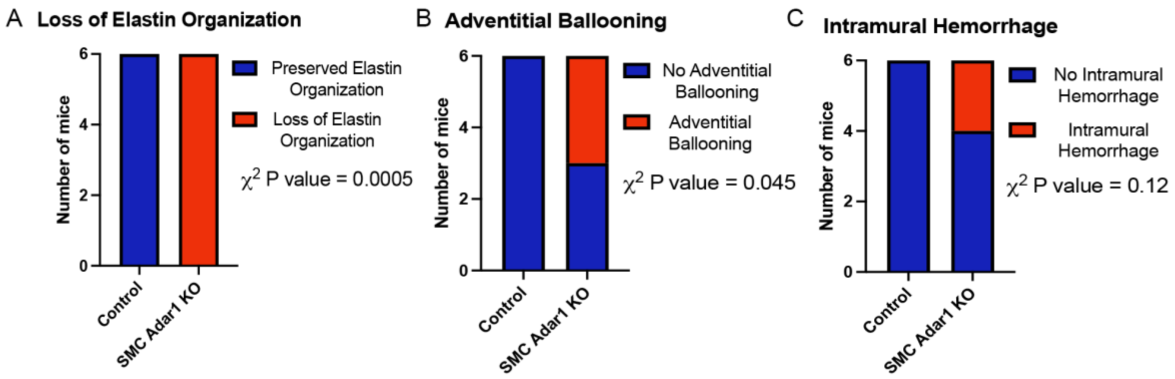


**Supplemental Figure 7. Quantification of histological findings for control and *SMC-Adar1 KO* mice**. (A) loss of elastin organization, (B) adventitial ballooning, and (C) intramural hemorrhage. P values represent $\chi^2$ statistical comparison between groups. N = 6 per group.

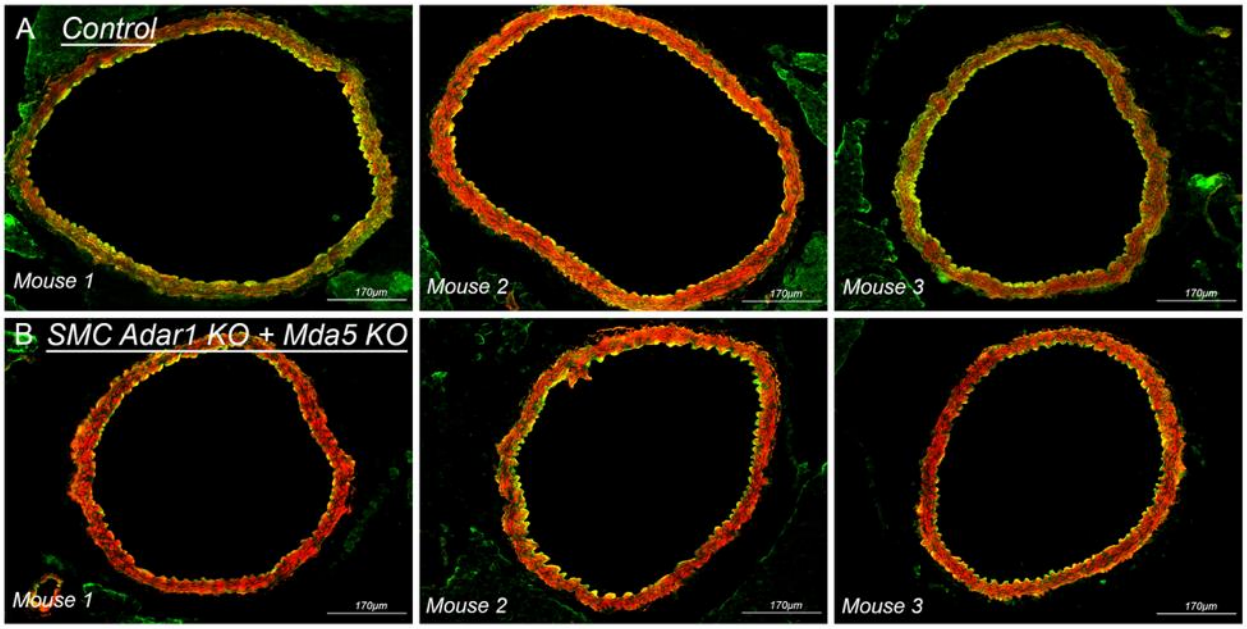


**Supplemental Figure 8.** ***Aortas from SMC Adar1 KO with Mda5 KO are indistinguishable from control.*** Representative fluorescent images of aorta from (A) control (Myh11$^{CreERT2}$, ROSAtdTomato, ApoE$^{-/-}$) and (B) SMC-Adar1 KO + Mda5 KO (Adar1$^{fl/fl}$, *Ifih1*$^{-/-}$, Myh11$^{CreERT2}$, ROSAtdTomato, ApoE$^{-/-}$) mice at 2 weeks post tamoxifen. Red represents tdTomato and green represents autofluorescence. Scale bar = 170$\mu$m.

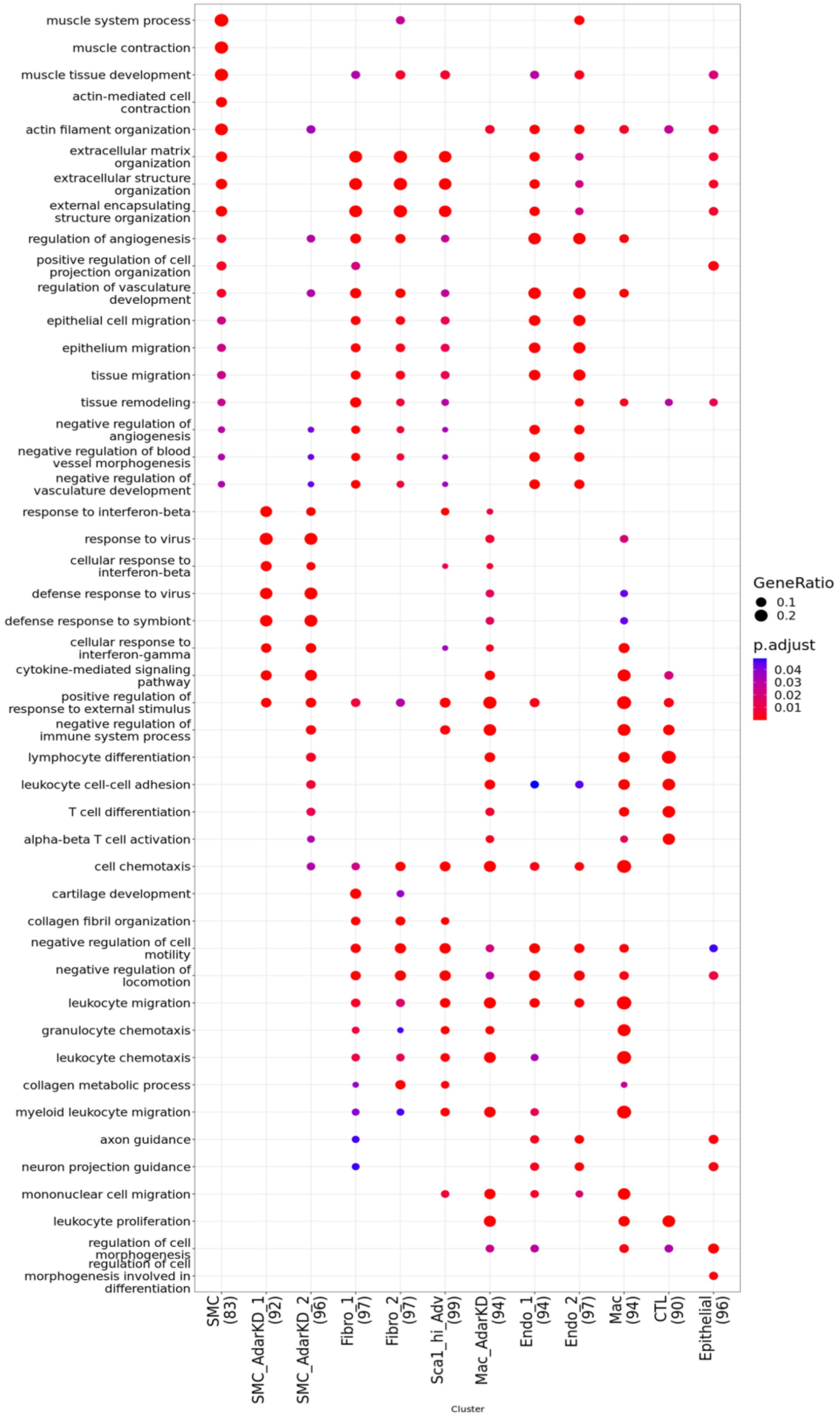


**Supplemental Figure 9.** ***SMC Adar1 KD induces antiviral defense transcriptional pathways.*** GO term enrichment analysis of scRNAseq data clusters at 2 weeks following tamoxifen treatment for SMC-Adar1$^{KO}$ (Adar1$^{fl/fl}$, Myh11$^{CreERT2}$, ROSAtdTomato, ApoE$^{-/-}$) and control (Myh11$^{CreERT2}$, ROSAtdTomato, ApoE$^{-/-}$) genotypes. Parentheses reflect distinct genes included in pathway enrichment analysis.

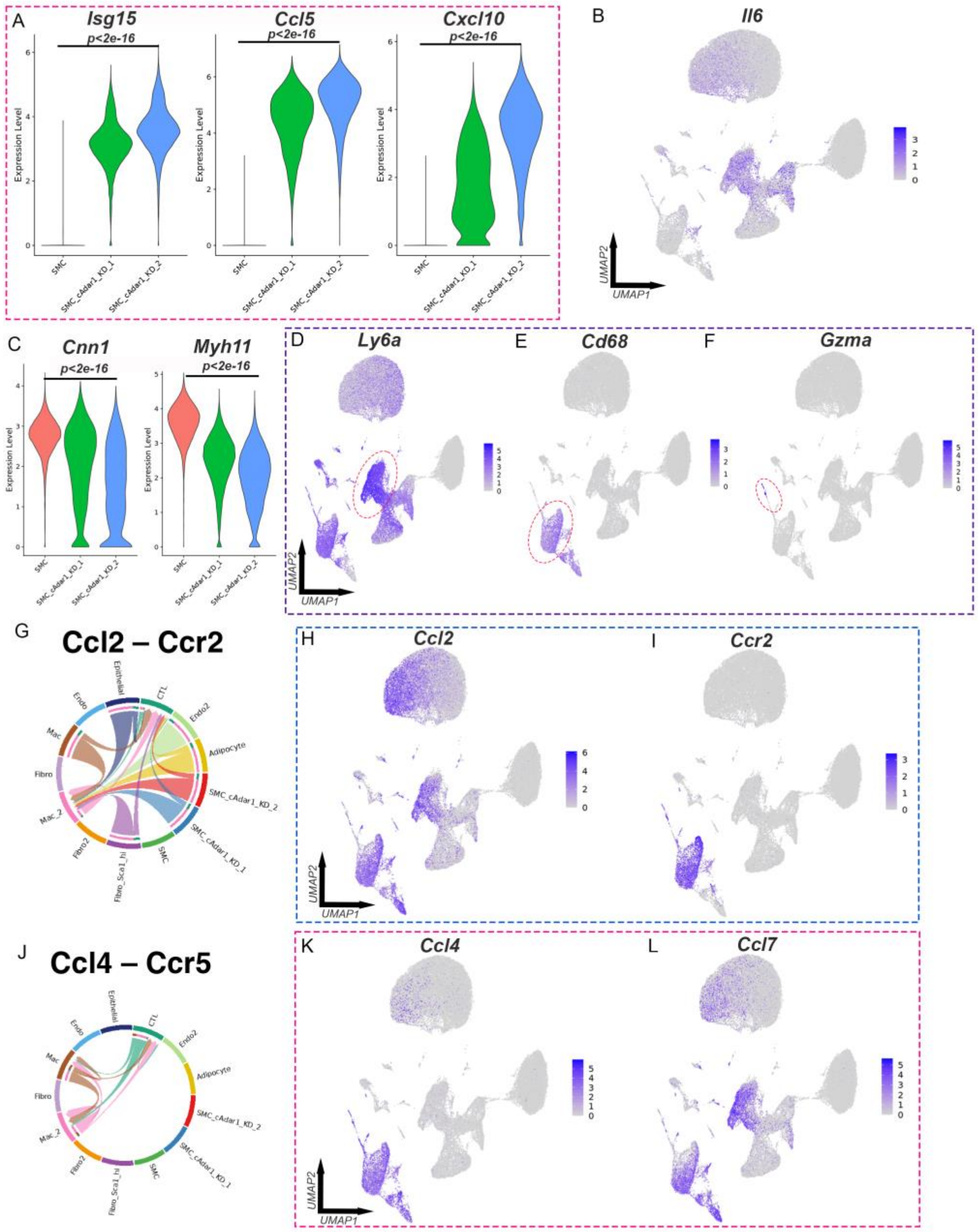


**Supplemental Figure 10.** ***Loss of SMC Adar1 coordinates distinct response throughout vessel wall.*** (A) Violin plot of *Isg15*, *Ccl5*, and *Cxcl10* within SMC, SMC_cAdar1_KD_1, and SMC_cAdar1_KD_2 clusters of scRNAseq data clusters at 2 weeks following tamoxifen treatment for *SMC-Adar1*$^{-/-}$ (*Adar1*$^{fl/fl}$*, Myh11*$^{CreERT2}$*, ROSAtdTomato, ApoE*$^{-/-}$) and control (*Adar1*$^{WT/WT}$*, Myh11*$^{CreERT2}$*, ROSAtdTomato, ApoE*$^{-/-}$) genotypes. (B) Featureplot of *Il6*, (C) *Ly6a* (Sca1), (D) *Cd68*, and (E) *Gzma*. (G) Violin plot of *Cnn1* and (G) *Myh11* in SMC, SMC_cAdar1_KD_1, and SMC_cAdar1_KD_2 clusters. (F) Chord diagram of receptor-ligand interaction between clusters for CCL2:CCR2. (G) Featureplot of *Ccl2* and (H) *Ccr2*. (I) Chord diagram of receptor-ligand interaction between clusters for CCL4:CCR5. (J) Featureplot of *Ccl4* and (K) *Ccl7*.

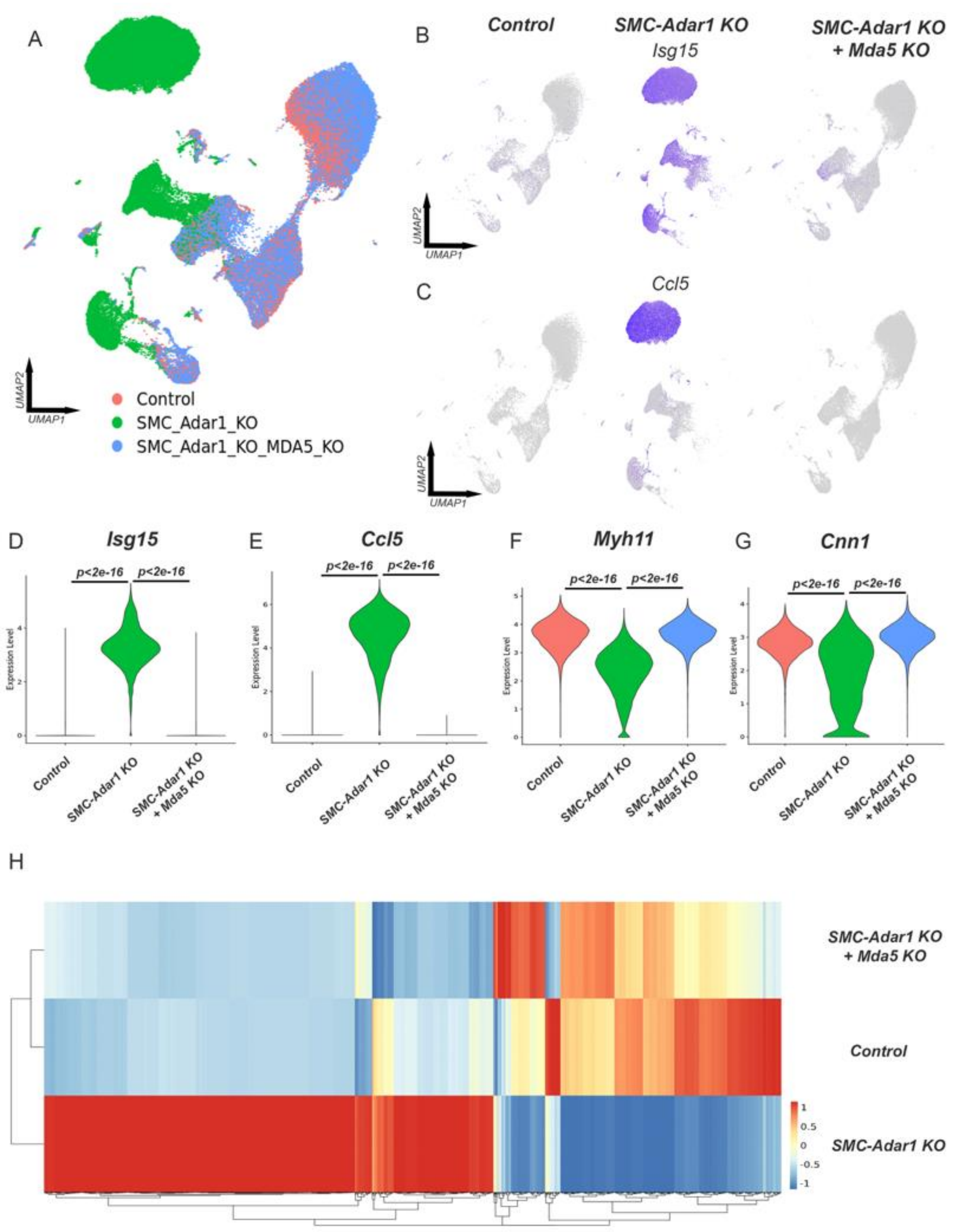


**Supplemental Figure 11.** ***Homozygous deletion of MDA5 (Ifih1) prevents transcriptomic effect of SMC-Adar1 KO.*** (A) UMAP of integrated scRNAseq data set between control (*Adar1$^{WT/WT}$, Myh11$^{CreERT2}$, ROSAtdTomato, ApoE$^{-/-}$*), *SMC-Adar1 KO* (*Adar1$^{fl/fl}$, Myh11$^{CreERT2}$, ROSAtdTomato, ApoE$^{-/-}$*), and *SMC-Adar1 KO, Mda5 KO* (*Adar1$^{fl/fl}$, Ifih1$^{-/-}$, Myh11$^{CreERT2}$, ROSAtdTomato, ApoE$^{-/-}$*) genotypes. Featureplot of (B) *Isg15* and (C) *Ccl5* split by genotype. Violin plot of tdTomato+ subset analysis split by genotype for (D) *Isg15*, (E) *Ccl5*, (F) *Myh11*, (G) *Cnn1*. (H) Heatmap visualization for normalized expression of marker genes for control, *SMC-Adar1 KO*, and SMC-*Adar1 KO + Mda5 KO* genotypes in tdTomato+ subset analysis.

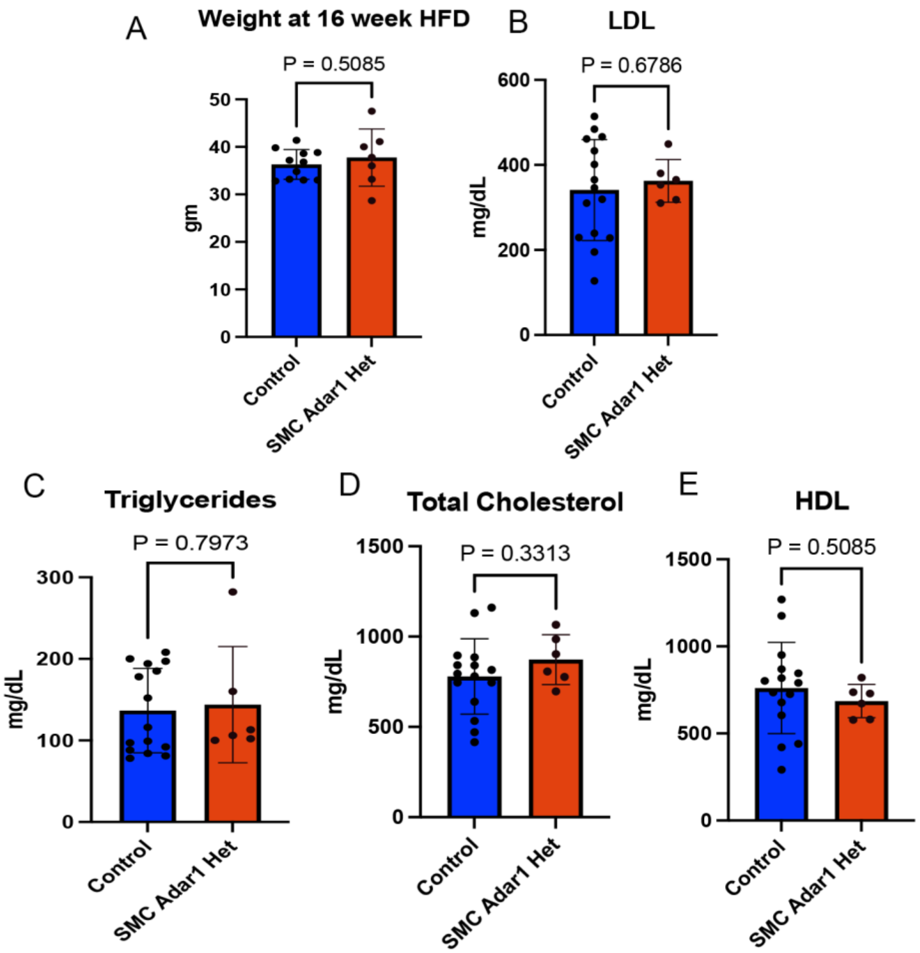


**Supplemental Figure 12.** ***SMC Adar1 haploinsufficiency has no effect on weight or cholesterol in atherosclerosis high fat diet model.*** SMC Adar1$^{-/+}$ versus control following 16 weeks high fat diet in atherosclerosis model for (A) weight (gm), (B) LDL cholesterol (mg/dL), (C) triglycerides (mg/dL), (D) total cholesterol (mg/dL), (E) HDL cholesterol (mg/dL). N = 13 (control) and N = 7 (SMC Adar1$^{-/+}$). P values represent T test for comparison.

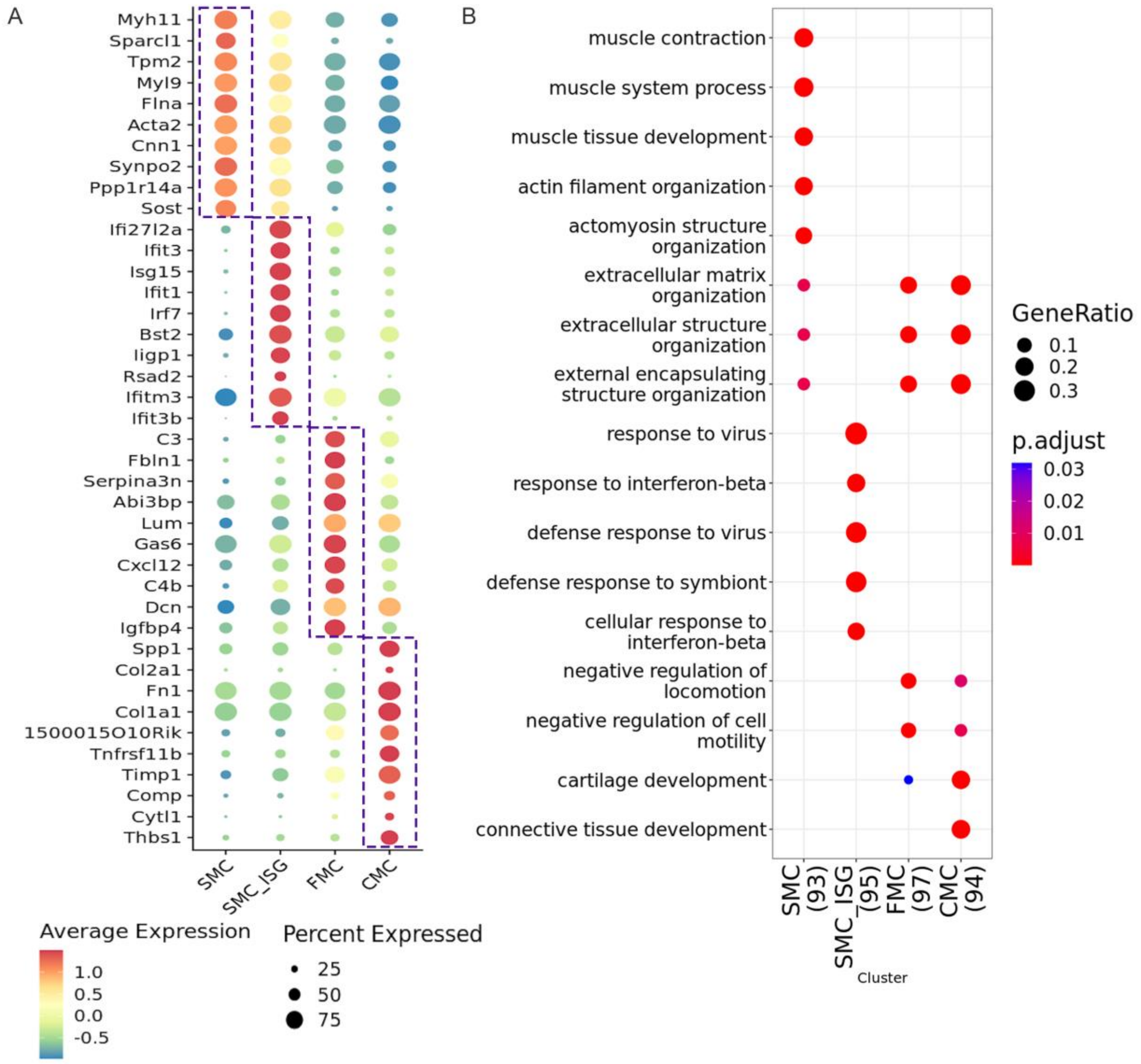


**Supplemental Figure 13.** ***SMC specific haploinsufficiency of Adar1 in SMC subset analysis shows ISG activation with phenotypic modulation in atherosclerosis.*** (A) Dotplot of top 10 genes for each of the 4 SMC clusters. (B) Top GO term enrichment for gene markers for each SMC cluster. Parentheses reflect distinct genes included in pathway enrichment analysis.

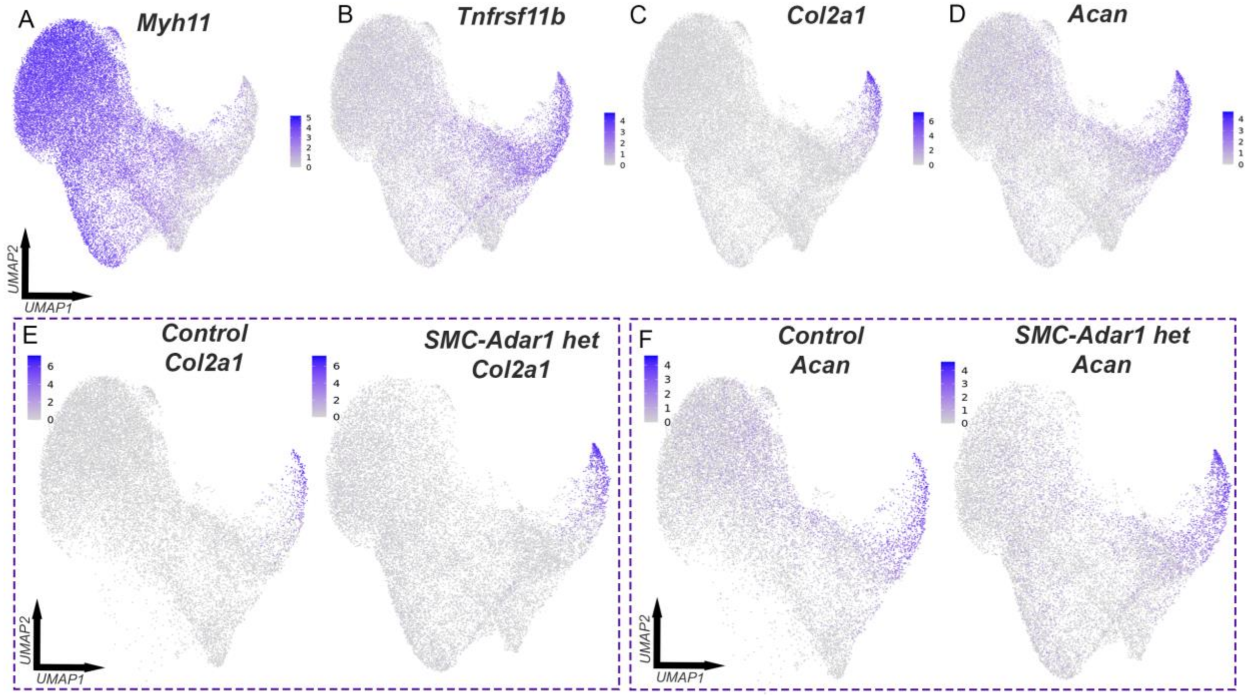


**Supplemental Figure 14.** ***SMC specific haploinsufficiency of Adar1 increases vascular chondromyoctye formation*** (A) Featureplot of *Myh11*, (B) *Tnfrsf11b*, (C) *Col2a1*, and (D) *Acan* of scRNAseq SMC subset data from atherosclerotic aortic root and ascending aorta at 16 weeks high fat diet in *SMC-Adar1*$^{-/+}$ (*Adar1*$^{fl/WT}$, *Myh11*$^{CreERT2}$, *ROSAtdTomato*, *ApoE*$^{-/-}$) and control (*Adar1*$^{WT/WT}$, *Myh11*$^{CreERT2}$, *ROSAtdTomato*, *ApoE*$^{-/-}$) genotypes. (E) Featureplots of *Col2a1* and (F) *Acan* split by genotype.

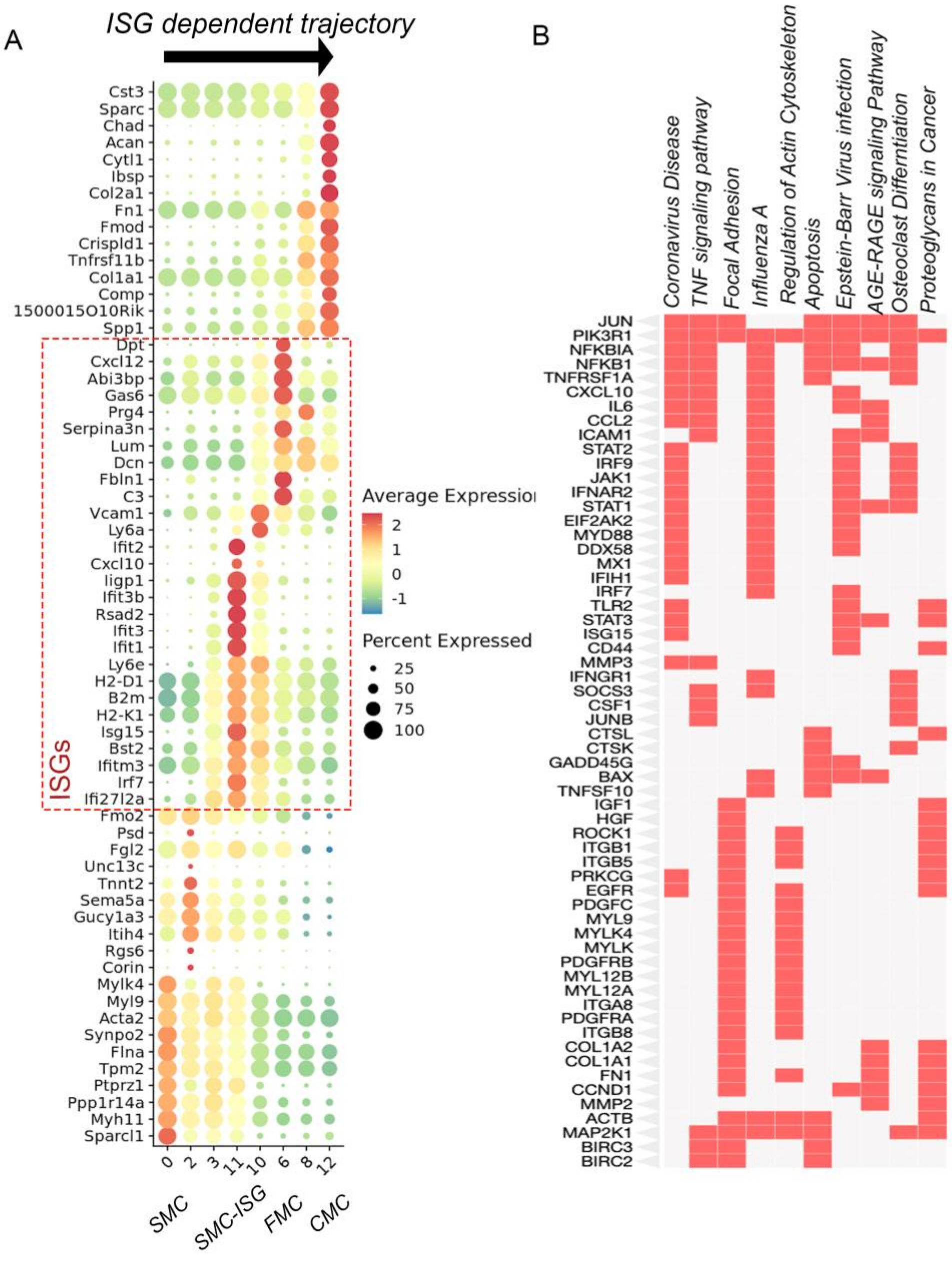


**Supplemental Figure 15.** ***ISG dependent trajectory analysis from SMC to CMC implicates distinct gene ontologies.*** (A) Top genes for each ISG dependent trajectory cluster in DotPlot. (B) Top gene ontology pathways for marker genes of distinct clusters from ISG dependent trajectory.

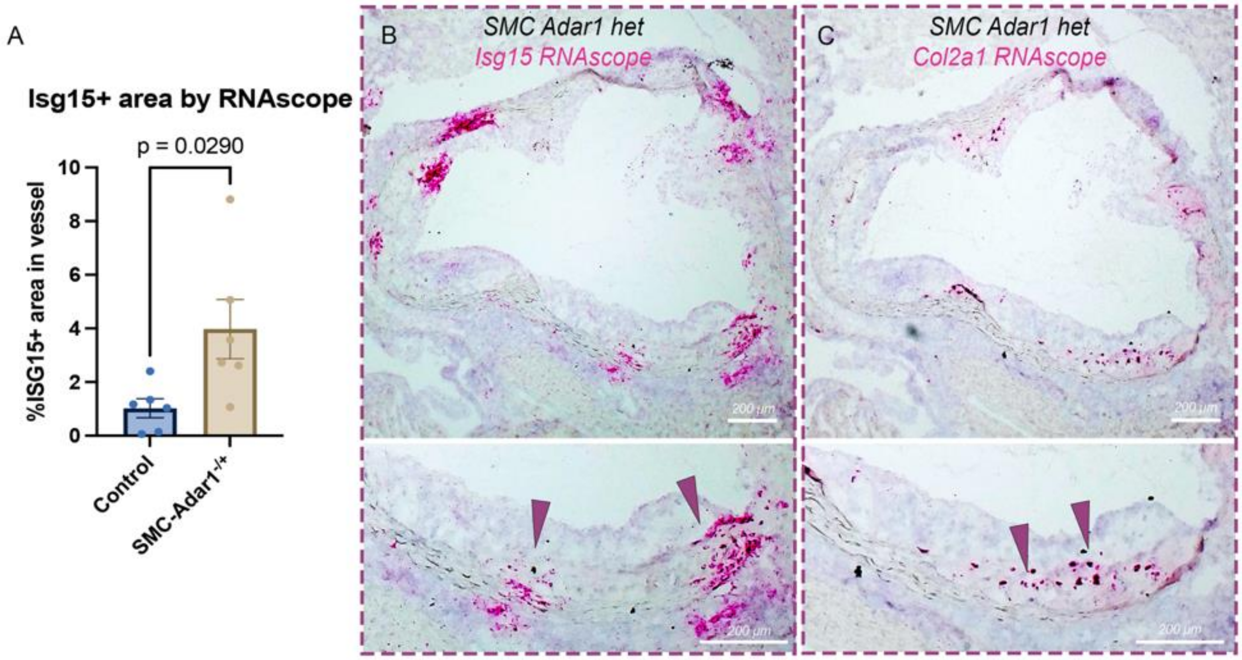


**Supplemental Figure 16. *Isg15 RNAscope reveals increased Isg15 signal in plaque of SMC Adar1$^{-/+}$ mice.*** (A) Quantification of *Isg15* + area within the vessel wall between control and *SMC Adar1$^{-/+}$* mice following 16 week of high fat diet (n = 6 per group). (B-C) Representative images of *Isg15* (B) and *Col2a1* (C) RNAscope. P values represent T test for comparison.

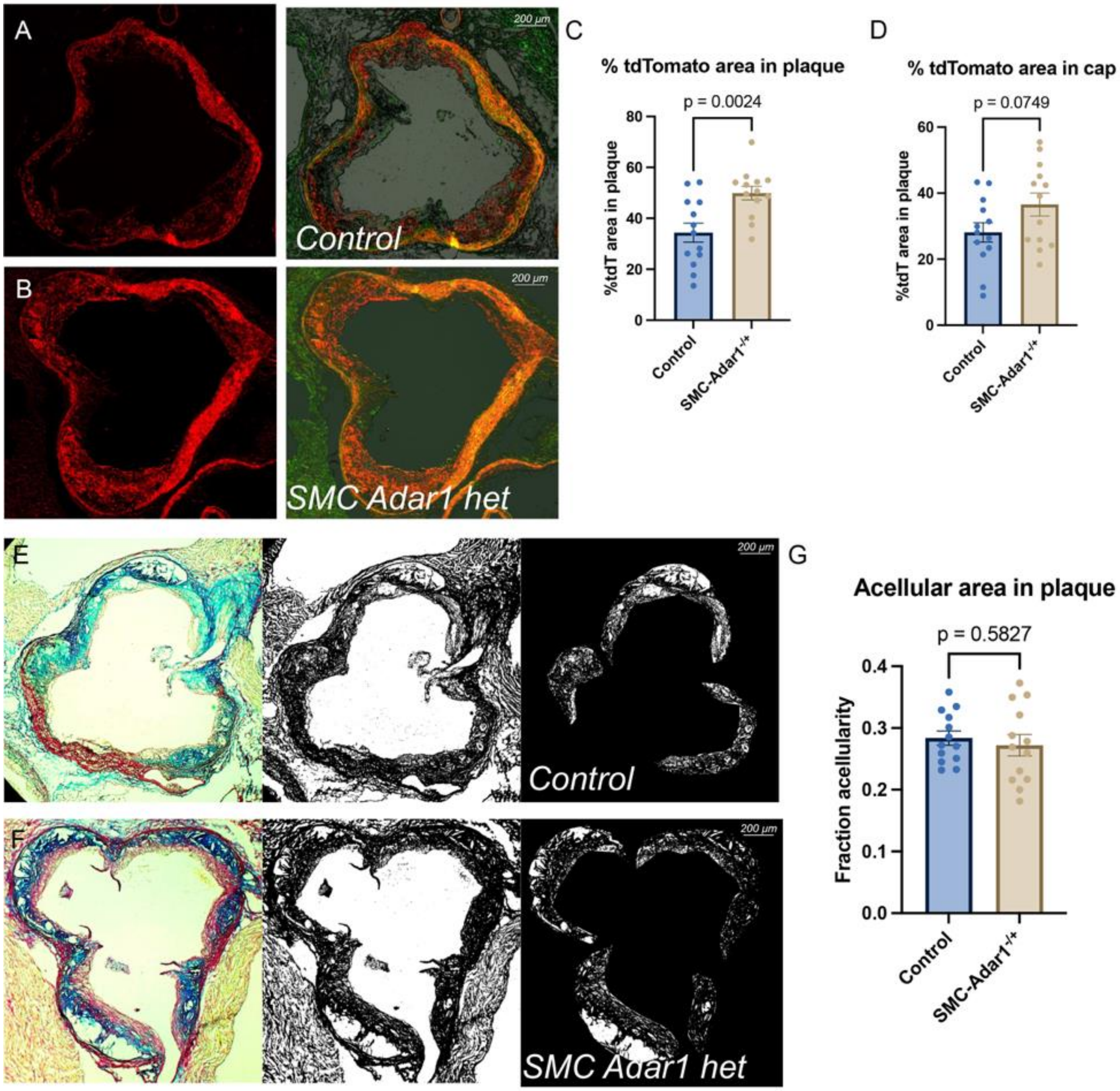


**Supplemental Figure 17.** ***SMC specific haploinsufficiency in Adar1 increased SMC lineage traced cell content in plaque without change in acellular area.*** Representative images of tdTomato (SMC derived) and overlay images with FITC and brightfield images in control (*Adar1*$^{WT/WT}$, *Myh11*$^{CreERT2}$, *ROSAtdTomato*, *ApoE*$^{-/-}$) (A) and SMC Adar1 het (*Adar1*$^{fl/WT}$, *Myh11*$^{CreERT2}$, *ROSAtdTomato*, *ApoE*$^{-/-}$)(B) genotypes. Quantification of the percentage of tdTomato positive area in the plaque (C) and in the top 30 $\mu$m segment of the plaque representing the cap (D). Masson's Trichrome staining with threshold analysis and acellular area quantification in control (E) and SMC-Adar1$^{-/+}$ mice (F) with quantification (G). N = 13 per group. P values represent T test for comparison.

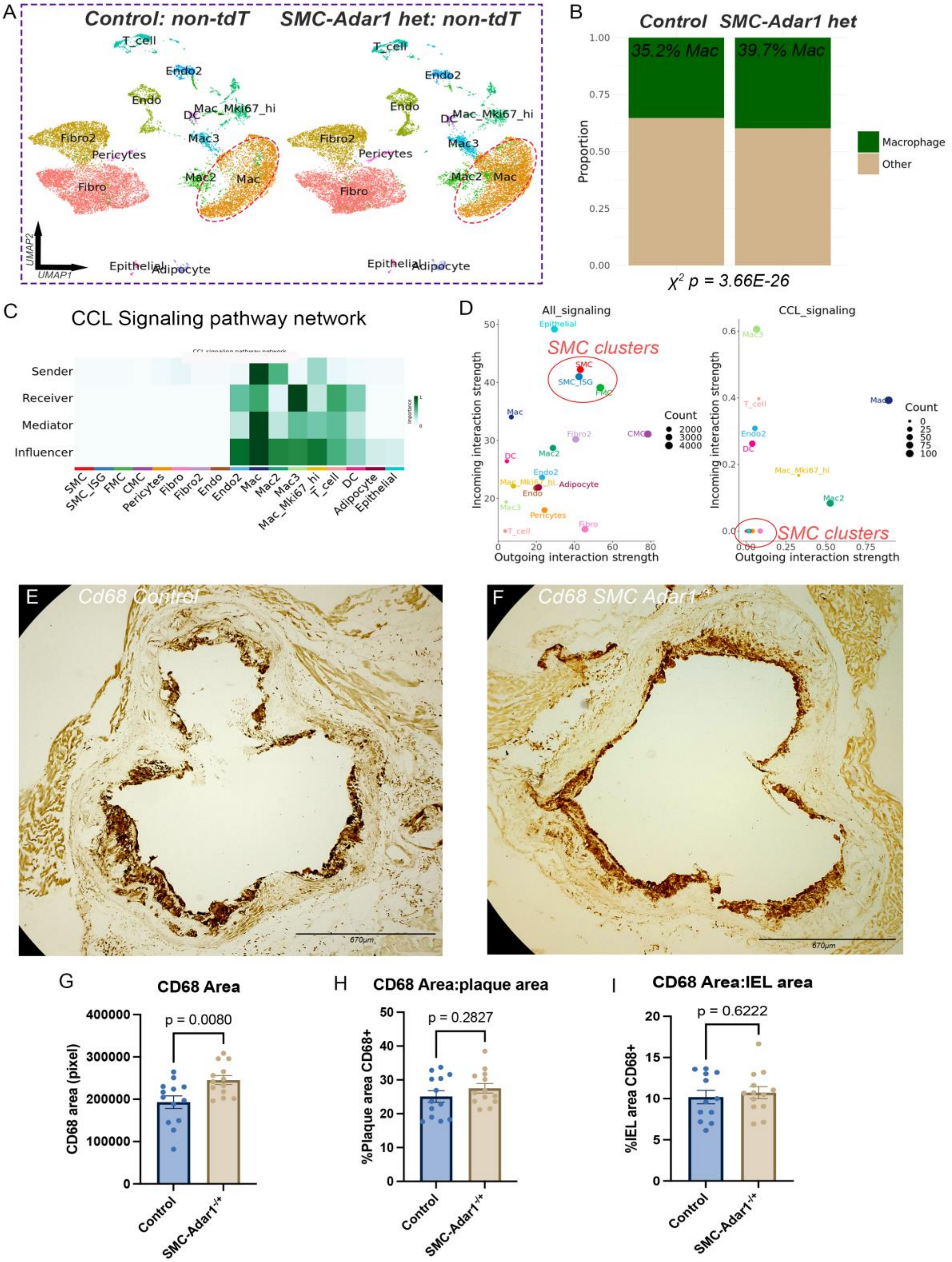


**Supplemental Figure 18.** ***SMC specific haploinsufficiency of Adar1 has minimal effect on macrophage infiltration in atherosclerosis.*** (A) UMAP of scRNAseq data of non-tdTomato lineage traced cells from atherosclerotic aortic root and ascending aorta at 16 weeks high fat diet split by control and *SMC-Adar1*$^{-/+}$ genotypes. (B) Stacked bar chart comparing proportion of macrophage cell population between control and *SMC-Adar1*$^{-/+}$ genotypes in the non-tdT+ cells. (C) Heatmap of network centrality scores for CCL signaling network across cell clusters. (D) Dot plot of outgoing and incoming interaction strength across cell clusters for all signaling networks (left) and CCL signaling network (right). (E-F) Cd68 immunohistochemical staining within the aortic root following in control (E) and *SMC- SMC-Adar1*$^{-/+}$ (F) mice. (G) Total area of Cd68 positive stain showing a significant increase in Cd68 in *SMC-Adar1*$^{-/+}$ compared to control, however this effect is not significant when normalizing to (H) plaque area, or (I) vessel area at the internal elastic lamina (IEL). *P*-values represent chi-squared test (B) and unpaired two-tailed T-test (G-I).

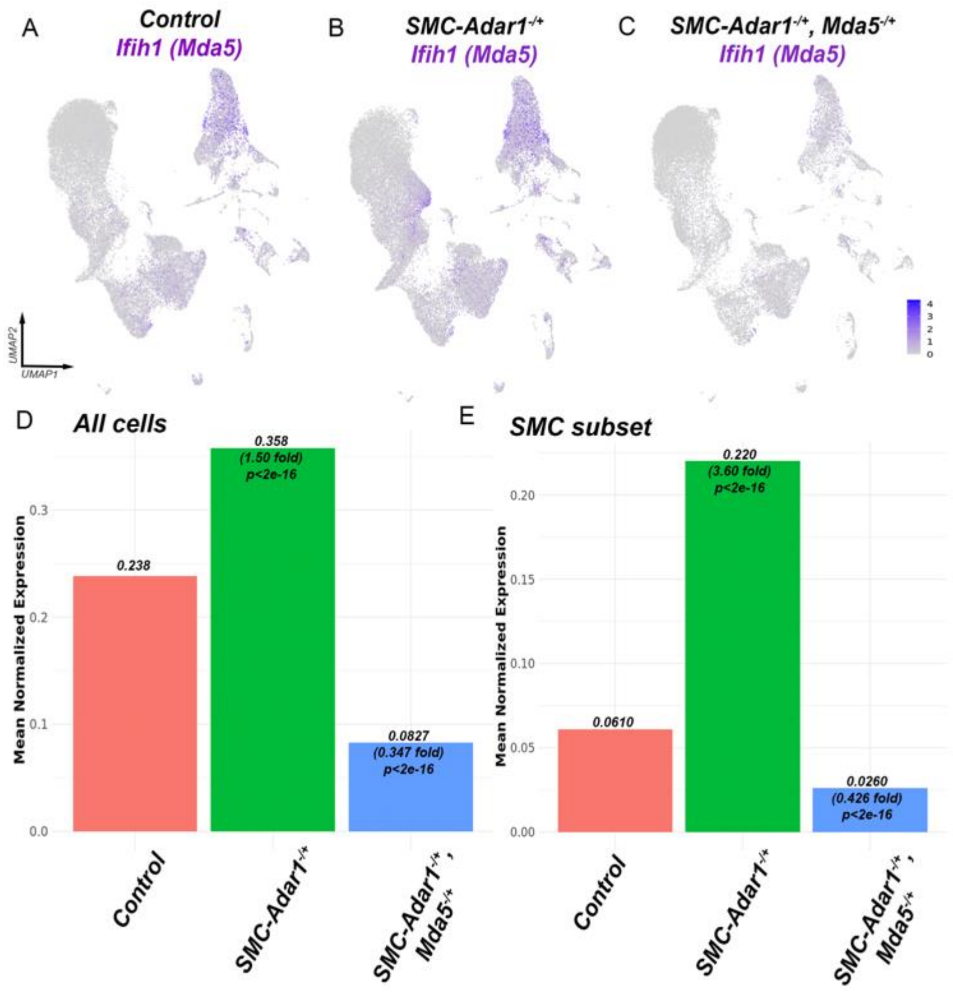


**Supplemental Figure 19. *Mda5 haploinsufficiency reduces Mda5 expression and prevents upregulation in SMC Adar1 het background.*** Featureplot of *Ifih1* (Mda5) for all cells from atherosclerotic aortic root and ascending aorta at 16 weeks high fat diet split by control (A), *SMC-Adar1*$^{-/+}$ (B), and *SMC-Adar1*$^{-/+}$, *Mda5*$^{-/+}$ (C) genotypes. Extracted normalized expression data of *Ifih1* (Mda5) for (D) all cells and (E) SMC subset displaying normalized expression values, fold change from control genotype, and p value for significance.

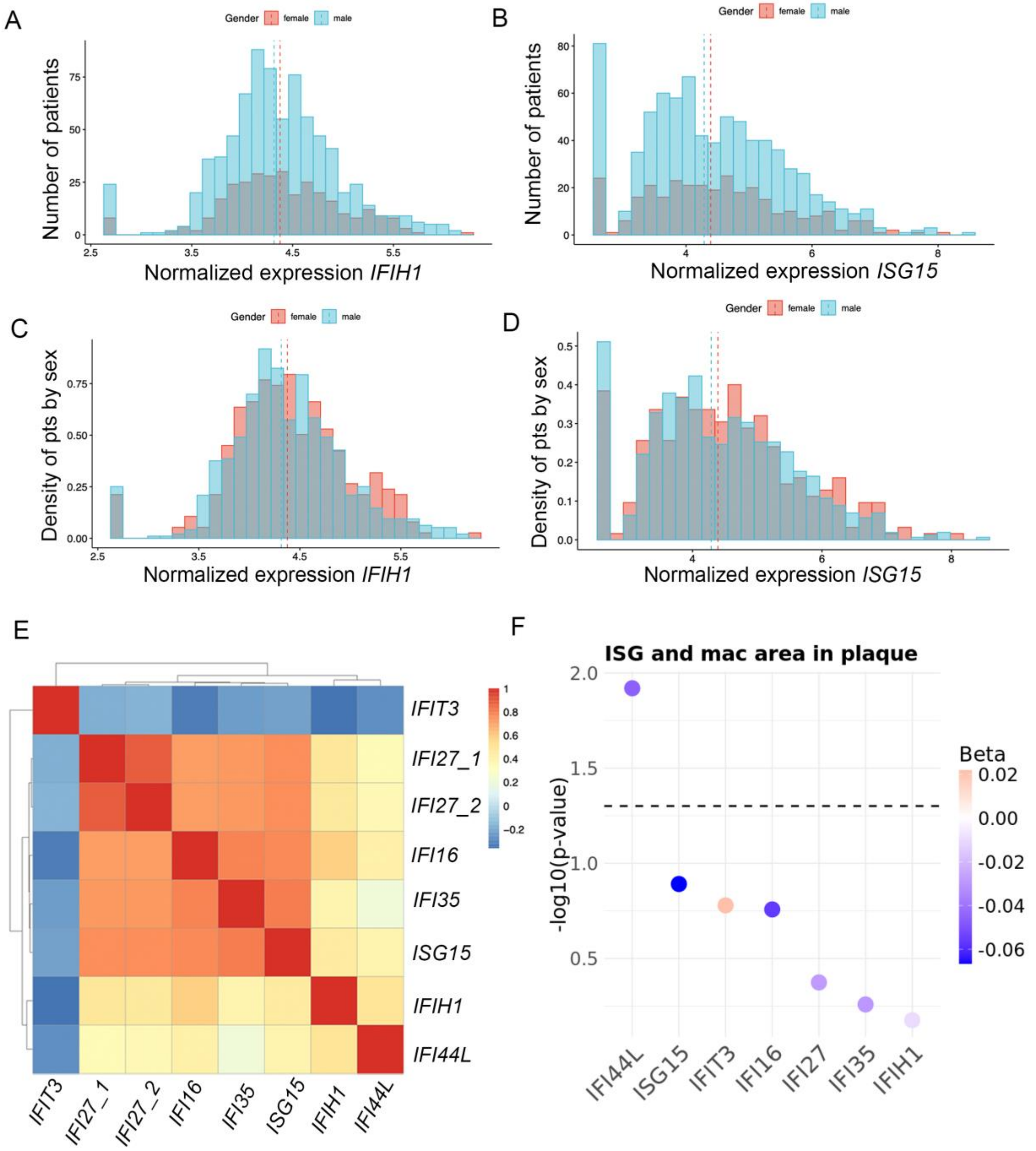


**Supplemental Figure 20.** ***Variability of ISG expression between patient carotid endarterectomy samples with no difference between sexes in Athero-Express cohort.*** Histogram plot of patient distribution for (A) normalized *IFIH1* and (B) *ISG15* expression split by sex (male – blue, female – red). Histogram plot of patient density by sex for (C) normalized *IFIH1* and (D) *ISG15* expression. Heatmap plot of correlation beta value for ISGs (E). (F) Dot plot displaying linear regression analysis between ISGs and macrophage area within plaque corrected for co-variates.
